# Massively parallel quantification of CRISPR editing in cells by TRAP-seq enables better design of Cas9, ABE, CBE gRNAs of high efficiency and accuracy

**DOI:** 10.1101/2020.05.20.103614

**Authors:** Xi Xiang, Kunli Qu, Xue Liang, Xiaoguang Pan, Jun Wang, Peng Han, Zhanying Dong, Lijun Liu, Jiayan Zhong, Tao Ma, Yiqing Wang, Jiaying Yu, Xiaoying Zhao, Siyuan Li, Zhe Xu, Jinbao Wang, Xiuqing Zhang, Hui Jiang, Fengping Xu, Lijin Zou, Huajing Teng, Xin Liu, Xun Xu, Jian Wang, Huanming Yang, Lars Bolund, George M. Church, Lin Lin, Yonglun Luo

## Abstract

The CRISPR RNA-guided endonucleases Cas9, and Cas9-derived adenine/cytosine base editors (ABE/CBE), have been used in both research and therapeutic applications. However, broader use of this gene editing toolbox is hampered by the great variability of efficiency among different target sites. Here we present TRAP-seq, a versatile and scalable approach in which the CRISPR gRNA expression cassette and the corresponding surrogate site are captured by **T**argeted **R**eporter **A**nchored **P**ositional **Seq**uencing in cells. TRAP-seq can faithfully recapitulate the CRISPR gene editing outcomes introduced to the corresponding endogenous genome site and most importantly enables massively parallel quantification of CRISPR gene editing in cells. We demonstrate the utility of this technology for high-throughput quantification of SpCas9 editing efficiency and indel outcomes for 12,000 gRNAs in human embryonic kidney cells. Using this approach, we also showed that TRAP-seq enables high throughput quantification of both ABE and CBE efficiency at 12,000 sites in cells. This rich amount of ABE/CBE outcome data enable us to reveal several novel nucleotide features (e.g. preference of flanking bases, nucleotide motifs, STOP recoding types) affecting base editing efficiency, as well as designing improved machine learning-based prediction tools for designing SpCas9, ABE and CBE gRNAs of high efficiency and accuracy (>70%). We have integrated all the 12,000 CRISPR gene editing outcomes for SpCas9, ABE and CBE into a CRISPR-centered portal: The Human CRISPR Atlas. This study extends our knowledge on CRISPR gene and base editing, and will facilitate the application and development of CRISPR in both research and therapy.

## INTRODUCTION

Clustered Regularly Interspaced Short Palindromic Repeats (CRISPR) and CRISPR-associated protein 9 (Cas9) are essential adaptive immune components in most bacteria. The system has successfully been harnessed for programmable RNA-guided genome editing in prokaryotes, humans and many other living organisms [1-5]. The *Streptococcus pyogenes* Cas9 (SpCas9) is the most extensively studied and broadly applied Cas9 protein, amongst other Cas9 orthologs (e.g. SaCas9, StCas9, NmCas9) [6-9] and Cas proteins (e.g. Cas12a, Cas13) [10, 11]. Guided by a programmable small RNA molecule (also known as gRNA), the SpCas9 protein introduces a double-stranded DNA break (DSB) to the DNA target site, which constitutes a complementary protospacer sequences and a canonical protospacer adjacent motif (PAM) [2]. The classical CRISPR gene editing is achieved by reparation of the DSBs in living organisms by the endogenous DNA repair mechanisms, predominantly by the NHEJ and MMEJ pathways in mammalian cells. This process generates indels (deletions or insertions) to the repaired site [12]. It is thus essential to have data from CRISPR editing in cells to develop accurate prediction rule sets of CRISPR activity.

The simplicity of the CRISPR system, the flexibility for modifying the Cas9 protein, and the increasing efforts from CRISPR scientists and pharmaceutical companies have extensively broadened the CRISPR-Cas9-based gene editing toolkits. We are now enabled to epigenetically perturb endogenous gene expression [13, 14], fluorescently label endogenous DNA elements[15] and site-specifically edit single nucleotides [16-20]. The CRISPR base editors, which comprise two major classes: adenine base editors (ABE) and cytosine base editors (CBE), have increasingly evolved as attracting tools for gene editing. These base editors are created by fusing a catalytically dead Cas9 (dCas9) or Cas9 nickase (nCas9) to either an adenine deaminase or a cytidine deaminase [18, 19]. Without introducing double stranded DNA breaks, the ABE and CBE base editors, respectively, can efficiently create an A to G (or T to C on the complementary strand) and C to T (or G to A on the complementary strand) substitution within a small editing window of the target site [16, 21-23]. Albeit all these fantastic developments and applications of the CRISPR-Cas9 gene editing theme, there is still an urgent need of methods and high throughput data on the Cas9-induced DBS repair outcomes, as well as ABE and CBE efficiencies, to ensure a successful CRISPR gene editing outcome. Such cataloged data of Cas9 and base editor efficiencies will allow the selection of experimentally validated gRNAs, as well as for developing better rules for *in silico* Cas9, ABE, and CBE gRNAs design.

Quantification of gRNA activity at the endogenous sites in cells is limited by scale. *In vitro* approaches (in a test-tube) can overcome the scale but fails to recapitulate the effects of genome and epigenome architectures and cellular DNA repair mechanisms on CRISPR editing [24, 25]. Methods based on integrating synthetically barcoded DNA constructions were developed for large-scale measuring of Cas9-induced DSB repair outcomes of gRNAs in cells [26-28]. Currently, we lack large-scale ABE and CBE editing data for developing better rules for designing base editing gRNAs. In this study, we developed an assay system for massively parallel quantification of a large-scale CRISPR gRNAs activities in human cells. We optimized the design and procedures for generation and in-cell CRISPR editing of synthetically barcoded DNA constructs. Each construct contains a unique gRNA expression cassette and the corresponding surrogate target site. Using this method, Targeted Reporter Anchored Positional Sequencing (hereafter referred as **TRAP-seq**), we demonstrated the applicability of TRAP-seq for massively parallel quantification of the SpCas9-induced DSB repair outcomes, ABE and CBE efficiency and profiles for 12,000 gRNAs in human embryonic kidney cells.

## RESULTS

### Design and functional validation of the lentivirus-based TRAP-seq system

To streamline vector cloning, gRNA expression and delivery into cells, we firstly designed a lentivirus-based system with four main features: (1) A human U6 promoter; (2) Golden-Gate Assembly (GGA) based cloning with a *lac Z* marker for precise and efficient insertion of an expression cassette; (3) A green fluorescent protein (GFP) marker for measuring viral titer and real-time tracking of viral delivery; (4) A puromycin selection gene for enrichment of stably transduced cells (**Fig. 1a and S1**). Essentially, this lentivirus system allows conventional GGA-based insertion of a synthetic DNA containing a gRNA spacer, scaffold and the corresponding surrogate target site after the U6 promoter. As current microarray-based method can only faithfully synthesize oligo pools of max 170 bp, we optimized the DNA design to contain a 102bp gRNA expression cassette (20bp spacer + 82bp scaffold) and a 37bp surrogate target site, flanked by a 31bp GGA cloning site and PCR handles (**Fig. 1a and S1**). We and several other groups previously demonstrated that such a surrogate target site can faithfully recapitulate the endogenous editing efficiency and indel profile [27, 29]. To further validate the 37 bp surrogate target site, we firstly generated HEK293T cells expressing a doxycycline (Dox)-inducible SpCas9 [1], an adenine base editor (ABE7.10) [19] or a cytosine base editor (CBE, BE4-Gam) [30]. Next, we performed ICE-based analysis of three different sites (*AAVS1, INHCB, TYMP*) in the HEK293T-SpCas9, HEK293T-ABE and HEK293T-CBE cells. The results validated that the CRISPR editing efficiency and outcomes from the surrogate sites were closely correlated **(*r***^***2***^ ***= 0.96 – 0.99*)** with those from the endogenous genome sites (**Fig. S2**). For simplification, we hereafter named the system as Targeted Reporter Anchored Positional Sequencing (TRAP-seq), the 170bp synthetic oligo/DNA as TRAP oligo/DNA, and the 37bp surrogate target site as TRAP site.

**Figure 1.**
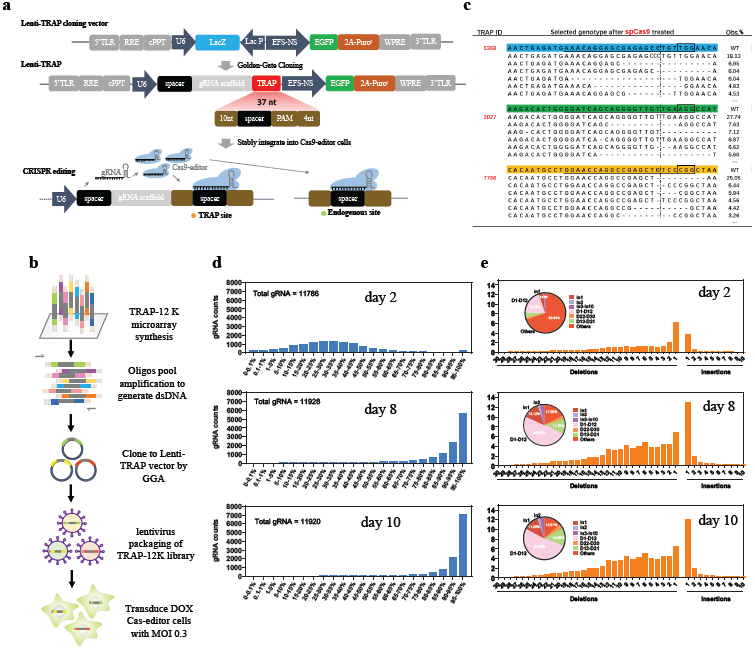
High Throughput Quantification of SpCas9 efficiency in cells by TRAP-seq. **a.** Schematic illustration of the TRAP-seq system. TLR, long terminal repeat; RRE, Rev Response Element; cPPT, central polypurine tract; U6, human U6 promoter; EFS-NS, short EFS promoter derived from EF1a. Figure not drown to scale. **b.** Schematic illustration of oligo pool synthesis, PCR amplification, gold-gate assembly, lentivirus packaging, and transduction of the 12K TRAP-seq library. **c.** Representative quantification of top 5 indel types for 3 TRAP sites. Dash line indicates the DSB site. Results for the 12,000 TRAP sites can be found at the CRISPR atlas database. **d.** Bar plots of SpCas9 editing efficiency of all TRAP sites measured by targeted amplicon sequencing. Results are shown for Dox-induced HEK293T-SpCas9 cells from 2, 8 and 10 days after transduction. Corresponding results for Dox-free HEK293T-SpCas9 are shown in Fig. S10. **e.** Bar plots of indel profiles (1-20 bp deletion, 1-10 bp insertion) for all TRAP sites introduced by SpCas9 in the Dox-induced HEK293T-SpCas9 cells from at 2, 8, and 10 days post transduction. Pie chat quantified the proportion of major indel groups: 1bp insertion (ins), 2bp insertion, 3-10 bp insertion, 1-12 bp deletion, 13-21bp deletion and 22-30 bp deletion. Other indels and wild-type reads are presented together as “others”. Corresponding results for Dox-free HEK293T-SpCas9 are shown in Fig. S11 and S12.

### Generation of 12K TRAP-seq lentiviral library

We next generated a 12K TRAP-seq library comprising 12,000 TRAP oligos by microarray synthesis (**Fig. 1b**). The library targets 3,834 human protein-coding genes (**Table S1**) [31]. The gRNA spacers were selected from the iSTOP database [32]. Out optimized workflow (also seen in methods) for PCR amplification of the 12K TRAP-seq oligos and cloning into the lentivirus-based TRAP-seq vector system is illustrated in **Fig. S3a.** A serial of optimizations in PCR conditions, GGA reactions and lentiviral packaging were carried out to avoid unspecific amplification (**Fig. S3b, c**), maximize successful ligation (**Fig. S4**) and properly quantify viral titer by FACS (**Fig. S5**), respectively.

To analyze the coverage of each TRAP oligo in our 12K TRAP-seq library, as well as to assess if the whole procedure of vector cloning and lentivirus packaging/transduction affected the overall TRAP representation, we performed targeted amplicon sequencing of the TRAP DNA in the 12K TRAP-seq oligo library, GGA plasmid DNA and wildtype HEK293T cells transduced with the 12K TRAP-seq lentivirus with a multiplexity of infection (MOI) of 0.3. With a constant sequencing depth (> 1,000X), all 12,000 TRAP oligos were detected in the 12K TRAP-seq library and the majority of TRAP oligos (> 90%) were evenly distributed (**Fig. S6a**). Most importantly, over 99% of the TRAP oligos were present in the 12K TRAP-seq plasmids and lentivirus preparation with high correlation of representation for each TRAP oligo (**r**^**2**^ **= 0.86-0.91, Fig. S6b**), suggesting that our optimized PCR, GGA, lentivirus packaging and transduction methods faithfully retained the complexity of the 12K TRAP-seq library without causing dramatic over/under-representation of the TRAP oligos.

### Quantification of SpCas9 editing at 12,000 sites by TRAP-seq

To demonstrate applicability of the 12K TRAP-seq lentivirus library, we firstly investigated massively parallel quantification of SpCas9 editing activity at all 12,000 TRAP sites. As schematically shown in **Fig. S7**, we transduced the Dox inducible HEK293T-SpCas9 cells with the 12K TRAP-seq lentivirus (MOI = 0.3 and transduction coverage = 4,690 cells per TRAP). Puromycin selection and Dox addition started two days after transduction to achieve maximum transduction and gene editing efficiency (**Fig. S8**). To enable comparison and identification of CRISPR-introduced indels, we also transduced wildtype HEK293T cells with the 12K TRAP-seq lentivirus with same MOI and transduction coverage. We harvested genomic DNA from the transduced cells at three time points: 2, 8, and 10 days after transduction (**Fig. S7 and Fig. S8**). Targeted PCRs were performed with a pair of universal primers specifically amplifying the TRAP DNA, followed by targeted deep sequencing with a DNA Nanoball sequencing technology [33]. With a constant sequencing depth (**Fig S9**), the proportional representation of each TRAP correlated better in the Dox-free HEK293T-SpCas9 cells (**r = 0.95**) than that in the Dox-induced HEK293T-SpCas9 cells (**R = 0.88**). Similar to CRISPR knockout screening pool libraries [34, 35], our results suggested that there existed similar cell fitness-related enrichment and depletion of the gRNAs in the 12K TRAP-seq library.

To measure the SpCas9 editing outcome, we firstly filtered out indels commonly found in both WT and SpCas9 HEK293T cells (also see methods), which were introduced by oligo synthesis or PCRs. We next analyzed the editing frequency and indel profiles for all 12,000 TRAP sites (**Fig. 1c, Table S2**, also see CRISPR Atlas below). Although the SpCas9 expression was Dox inducible, significant editing efficiencies were detected for all gRNAs in the HEK293T-SpCas9 cells at 2, 8 and 10 days after transduction in Dox-free medium (**Fig. S10**), suggesting a substantial leakiness of SpCas9 expression. As expected, significantly higher editing efficiencies were achieved for all 12,000 gRNAs in Dox-addition HEK293T-SpCas9 cells at 8 and 10 days after transduction (**Fig. 1d**). These results support the notion that SpCas9 expression level and cultivation time affect gene editing efficiency [36]. We and others had demonstrated that the indel outcomes introduced by SpCas9 comprises mainly small deletions and insertions [37-39]. The distribution of indel profiles (deletion of 1-30 bp and insertion of 1-10 bp) of the 12K TRAP sites were thus assessed in the transduced HEK293T-SpCas9 cells. Two days after transduction, deletion or insertion of 1 bp were the two most frequent indel types in cells (**Fig. 1e, S11**). Following increased editing time (Dox-free groups, **Fig. S10**) and SpCas9 expression (Dox-induced groups, **Fig. 1e**), the frequency of other indel types rose significantly and 1 bp insertion was the most dominant indel type which is in agreement with previous findings (**Fig. 1e, S12**) [37]. With the indel outcome, we were capable of analyzing the mutation consequence of all indels on protein translation. More than 70% of the total indels led to out-of-frame genotypes (**Fig. S13**). In conclusion, we demonstrated that TRAP-seq is a simple method for massively parallel quantification of gRNA editing outcomes in cells.

### Characterization of nucleotide features affecting SpCas9 efficiency and indel outcomes

Development of more accurate rules for *in silico* CRISPR design relies heavily on datasets of gRNA activity and indel from a large number of gRNAs. The rich gRNA activity and indels profile data generated by TRAP-seq above were valuable for further improving the performance of CRISPR design. We sought to investigate if the gRNA activity and indel outcomes measured by the TRAP-seq can mirror previous findings about the effects of nucleotide features on CRISPR activity. Nucleotide features such as secondary structure [24] and GC content [40] of the guide sequences affect CRISPR editing efficiency. We analyzed the correlation between gRNA activity and GC content and secondary structure (deltaG energy) in the gRNA spacer. Our TRAP-seq results further confirmed that the gRNA spacer GC content (**Fig. 2a, S14**) and secondary structure (**Fig. 2b, S15**) affected SpCas9 gene editing efficiency in cells. The optimal GC content and deltaG energy is [50-70%] GC and [-5; -1] KJ/mol, respectively. Consistent with the previous finding [41], our TRAP-seq results revealed that the SpCas9 disfavors motifs of “TT” and “GCC” at the N17-N20 region (**Fig. 2c, S16**). Recent reports have discovered that indel profiles for a given gRNA is predictable [27, 28]. The SpCas9 predominantly generates blunt-end double-stranded DNA breaks (DSB) between the N17 and N18 nucleotide preceding the protospacer adjacent motif (PAM), which are most frequently repaired by the NHEJ and MMEJ pathways in mammalian cells [42]. We compared the indel profiles of approximately 12,000 gRNAs revealed by TRAP-seq to the predicted indel profiles by inDelphi, a machine learning program for SpCas9 indel prediction [27]. Our results show that the overall indel profiles (in the Dox-free or Dox-induced cells at day 8 and 10) are highly correlated with the predicted ones by inDelphi (**median r = 0.51-0.65, Fig. 2d and Fig. S17**). Of note, the overall correlation between the TRAP indels in transduced cell at day 2 and indel profiles predicted by inDelphi was much lower (**median r = 0.31, Fig. S17**), suggesting that the indel outcome also depends on the experimental conditions (e.g. Cas9 expression level, editing time etc.).

**Figure 2.**
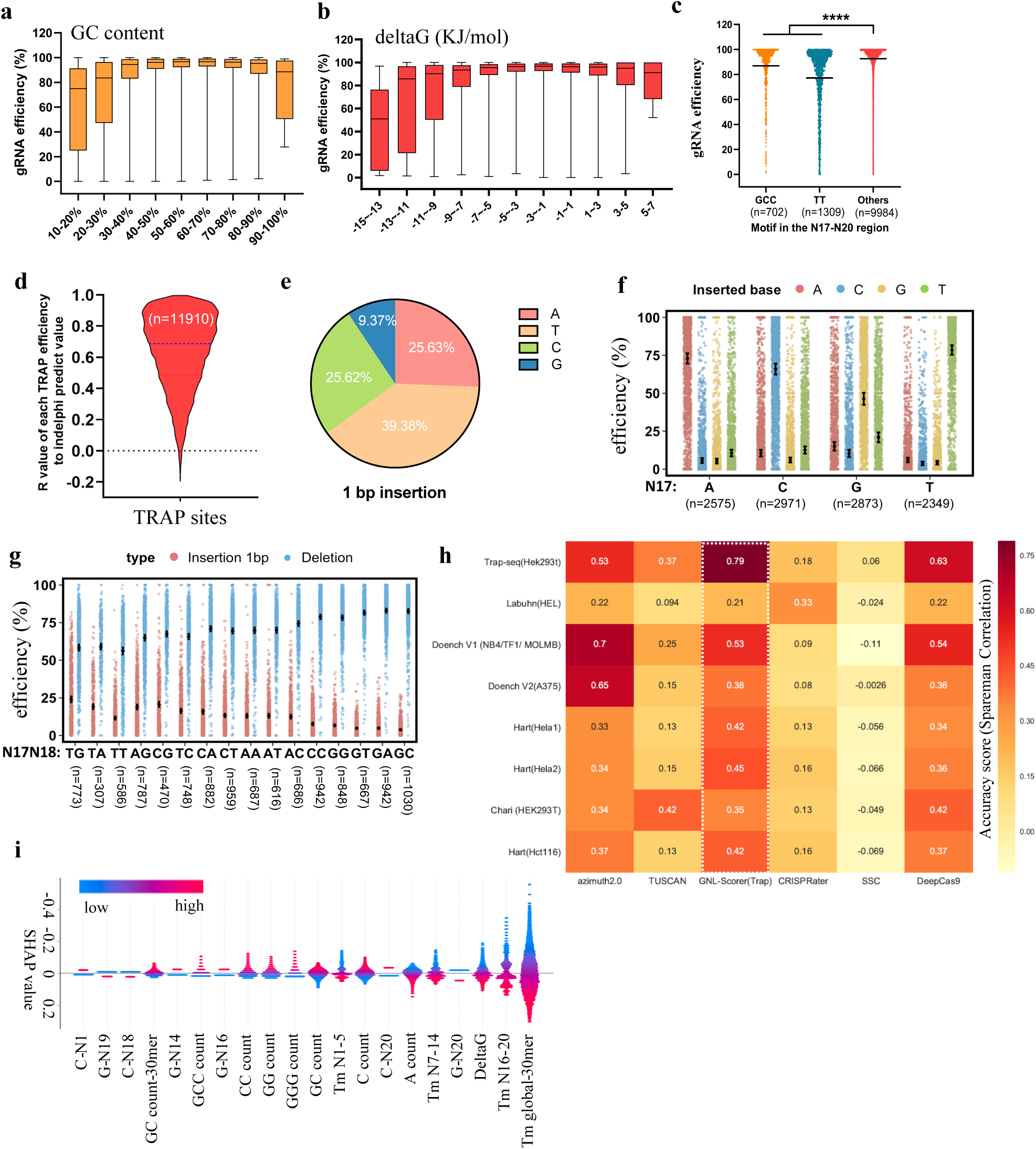
Characterization of features affecting SpCas9 efficiency and indel outcomes in cells. **a.** Box plot between GC content (with an interval of 10%) and SpCas9 gRNA efficiency measured in Dox-induced cells from Day 10. Results for other groups are shown in Fig. S14. **b.** Box plot between deltaG energy (with an interval of 2) and SpCas9 gRNA efficiency measured in Dox-induced cells from Day 10. Results for other groups are shown in Fig. S15. **c.** Comparison of SpCas9 efficiency between gRNAs harboring the GCC or TT motif in N17-N20 seed region and gRNAs without these two motifs. Data are shown for Dox-induced cells from Day 10. Results for other groups are shown in Fig. S16. “****”, p value less than 0.0001. **d.** Correlation between TRAP-seq indels and indels predicted by inDelphi from 11,910 sites. Data are shown for Dox-induced cells from Day 10. Results for other groups are shown in Fig. S17. **e.** Pie chart of the proportion of 1bp insertion among four bases (A, T, C, G) **f.** Correlation between the inserted 1 base and the nucleotide at N17 position. Data are shown for Dox-induced cells from Day 10. Results for other groups are shown in Fig. S18. **g.** Effects of N17N18 dinucleotide motifs on the indel frequency of 1bp insertion and deletions (1-30 bp). The gRNAs are divided into 16 groups based on the N17N18 motifs. For each gRNA, the total indel frequencies of 1-30bp deletions and 1bp insertion were analyzed. “n” indicates the number of gRNAs included for each group. **h.** Comparison of SpCas9 gRNA efficacy predictions in a regression schema for various datasets and prediction models. **i.** Top 20 features that weighted the most for the GNL machine learning model. Results are shown as the SHAP (SHapley Additive exPlanations) values. The 30mere comprises 4bp upstream, 20bp protospacer, 3 bp PAM, and 3 bp downstream sequences. Machine learning was based on gRNA efficiency data from Dox-induced cells at Day 10.

Among all indels, the 1bp insertion between N17 and N18 was the most abundant type (**Fig. 1e**). Previous studies had discovered that 1bp insertion is not random [27]. We therefor asked whether the nucleotides of 1bp insertion among our 12,000 TRAP sites followed the same principle. First, we quantified the frequency of inserted adenine (A), thymine (T), cytosine (C), guanine (G) among all 1bp insertions. The results show that the 1bp insertion favor T and disfavor G (**Fig. 2e**). For a given TRAP site, however, there is a strong preference of one nucleotide type (also seen **the CRISPR Atlas resource)**. Next, we divided all 12,000 TRAP sites into four groups based on the N17 or N18 nucleotide and quantified the frequency of inserted bases among all 1bp insertions. Our results show that the N17 nucleotide strongly defines the inserted base (**Fig. 2f, S18**), but less extensively affected by the N18 nucleotide (**Fig. S19**). Lastly, we divided all the 12,000 TRAP sites into 16 groups based on N17N18 sequence motifs and compared the indel frequency of 1bp insertion versus deletions. Despite a constantly higher frequency of deletions over insertions, motifs of GN (N = A, T, C, or G) and MC (M = A or C) at N17N18 favor deletions over 1bp insertion, as compared to TN and MG motifs (**Fig. 2g, S20**). Taken together, we show that high throughput TRAP-seq enables the identification and validation of features affecting SpCas9 editing efficiency and indel outcomes. The results corroborate previous findings that SpCas9 editing outcomes are predictable in cells.

### An improved machine learning model to predict SpCas9 efficiency

To further streamline the prediction of SpCas9 efficiency and the identification of nucleotide features important for gRNA activity, we randomly selected 80% of the 12K TRAP-seq gRNA efficiency and trained the GNL-Scorer [43] - a machine learning algorism that we previously developed based on the Bayesian Ridge regression (BRR) model and 2485 features. Our results showed that the GNL-scorer trained with the TRAP-seq dataset (GNL-Scorer (Trap)) gave an accuracy prediction score of over 70% (**Fig. S21**). To benchmark the performance of the TRAP-seq dataset and the GNL prediction algorithm, we compared our dataset and GNL-Scorer (Trap) with seven previously published datasets (from HEL, NB4, TF1, MOLMB, A375, Hela, HEK293T, HCT116 cells) and five prediction tools (DeepCas9, Azimuth-2.0, TUSCAN, CRISPRater, SSC). Our results showed that the GNL-Scorer (Trap) achieved the best accuracy score in five datasets (second best for the remaining 3 datasets) and have the best generalized prediction outcome across all test datasets (**Fig. 2h**). Using the SHapley Additive exPlanations (SHAP) algorithm for explaining the feature output, our results further revealed features (such as melting temperature, GC content delta G energy, sequences motifs etc.) that are important for the performance of our prediction model and SpCas9 efficiency (**Fig. 2i)**. Our results taken together suggest the TRAP-seq dataset based GNL-Scorer performs generally well for gRNA knockout efficiency prediction that the 12K SpCas9 gRNA efficiency dataset revealed by TRAP-seq enable better understanding of features affecting gRNA efficiency in cells, improve the design of gRNAs of high knockout efficiency. This improved GNL-scorer algorithm for predicting SpCas9 efficiency has been deposited to and available at public domain GitHub.

### Quantification of CRISPR-mediated adenine base editing at 12,000 sites by TRAP-seq

Unlike SpCas9 gene editing, we still lack large-scale data of CRISPR adenine base editing (ABE) efficiency. Such valuable data would enable us to develop better *in silico* ABE gRNA design tools. Since the TRAP site could confidentially recapitulate the ABE editing outcome of the corresponding endogenous site (**Fig. S2**), we sought to investigate the ABE editing outcomes in all 12,000 TRAP sites using the 12K TRAP-seq library. Although all the 12,000 gRNAs were not specifically designed for ABE editing, we reasoned that this “randomly” selected gRNA library would enable us to unbiasedly identify rules affecting ABE editing efficiency. To do this, we firstly transduced HEK293T-ABE cells with the 12K TRAP-seq lentivirus (MOI=0.3), and performed targeted amplicon sequencing of the TRAP DNAs from cells at 2, 7, and 11 days after transduction and cultured in Dox-free or Dox-addition medium (**Fig. S22**). For additional controls, we also performed similar experiments with wild-type HEK293T cells, with constant transduction coverage (4,690 cells per TRAP) and sequencing depth (approximately 1,000 reads per TRAP) (**Fig. S23**).

Next, we quantified the edited adenine events for each 37bp TRAP site in the 12K TRAP-seq library (**Fig. 3a, Table S3**). Our results showed that substantial editing events (within an editing window from N3 to N11) appeared 7 days after transduction in the HEK293T-ABE with Dox induction. The editing efficiency increased 4-5 folds when extending the cultivation time to 11 days (**Fig. 3b, S24**). Most importantly, the ABE editing window remained constant between N3 and N11 (**Fig. 3c, S25**), supporting the notion that the ABE base editor is highly conserved with respect to its editing region [19]. Our TRAP-seq results also validated that the highest ABE editing efficiency was observed for adenines located at N5, N6 and N7 (**Fig. 3d**). Lastly, quantification of all 42,790 edited adenine sites revealed that the ABE base editor conservatively generated A-to-G substitution (96.12%), and a small proportion of A-to-C (2.9%) and A-to-T (0.98%) substitutions (**Fig. 3e**). Although additional experiments will be required to test the TRAP-seq library in more cell lines, these initial studies suggest that the TRAP-seq is a highly valuable method for massively parallel quantification of CRISPR adenine base editing efficiency in cells.

**Figure 3.**
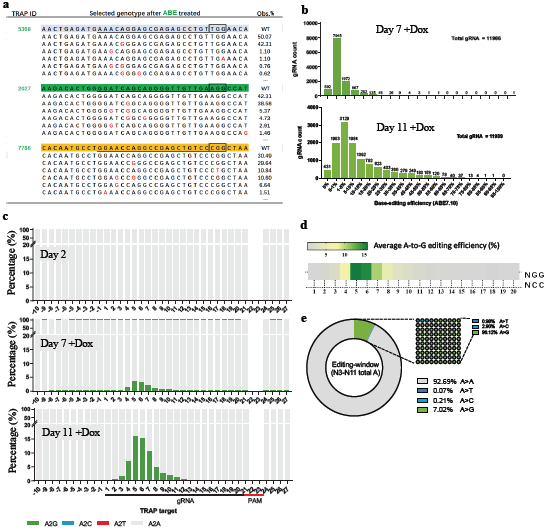
Quantification of ABE efficiency at 12,000 sites by TRAP-seq. **a.** Representation of top 5 adenine editing outcomes for three TRAP sites. Full ABE results for all 12,000 sites can be found at the CRISPR atlas resource. **b.** Quantification of overall ABE efficiency for all gRNAs from Dox-induced HEK293T-ABE cells 7- and 11-days post transduction. Other groups are presented in Fig. S23. **c.** Quantification of overall percentage of A-to-G, A-to-T, A-to-C, and A-to-A (unedited) events across the 37bp region of all TRAP sites. Results are from Dox-induced HEK293T-ABE cells at 2, 7- and 11-days post transduction. **d.** Heatmap quantification of the overall A-to-G editing efficiency within the 20nt protospacer region. **e.** Summary of substitution of Adenines within the ABE editing window N3-N11.

### Characterization of nucleotide features affecting ABE efficiency

We also sought to characterize features that affect ABE efficiency. To enable comparisons, we first selected two groups of gRNAs based on ABE efficiency: (1) high efficiency ABE gRNAs (n = 2,331, at least one edited adenine site had an efficiency over 20% with the protospacer region N1-N20) and (2) low efficiency ABE gRNAs (n = 2,589, the efficiency of any edited adenine site within the protospacer N1-N20 lower than 1%). Next, we compared the base percentage between the low and high efficiency gRNAs across the 37bp TRAP region for the low and high efficiency ABE gRNAs. Not surprisingly, high efficiency gRNAs showed overrepresentation of Adenine within the editing window N3-N7 (**Fig. S26**). Interestingly, high efficiency gRNAs favored Guanine over Thymine in the seed region (N17 to N20), the distal protospacer region (N-1 to N4) and the N21 base of the PAM. The presence of Cytosine in N20 was disfavored for high efficiency gRNAs.

We next sought to investigate the effect of intrinsic nucleotide preference on base editing efficiency. To identify the intrinsic nucleotide preference for ABE7.10, we focused on the ABE efficiency at N5, N6 and N7, which were the three highest ABE sites (**Fig. 3d**). Using a similar strategy to enable comparison, we first selected two groups of gRNAs: N5-N7 high efficiency (n = 2427, at least one edited adenine site had an efficiency over 20% within the core editing window N5-N7) and N5-N7 low efficiency (n = 2211, the efficiency of any edited adenine site within core editing window N5-N7 was lower than 1%). Next, we analyzed the preference of flanking bases (A, T, C, G) at N5-N7 for the low and high efficiency gRNAs. Our results revealed that the high efficiency ABE gRNAs favored upstream keto (K) bases (G, T) and downstream pyrimidine (Y) bases (T, C) (**Fig. 4a**). The presence of flanking adenine was strikingly overrepresented in the low efficiency ABE gRNAs (Fig. **4a**). To validate this observation, we analyze the correlation between A-to-G editing efficiency of all 12,000 TRAP sites in N5-N7 and their flanking bases. Consistent with the previous finding, the average ABE editing efficiency is higher if the edited adenine is flanked by Keto bases upstream and pyrimidine bases downstream (**Fig. 4b**), as compared to edited sites flanked by amino bases (A, C) upstream and purine bases (A, G) downstream. The overall ABE efficiency is much lower (approx. 2 to 7 folds) if the flanked base is adenine. Based on these observations, we further compared the frequency of tri-nucleotide flank-A motifs between the low and high ABE efficiency gRNAs. Using a cutoff of two folds, we categorized seven (B**A**C, K**A**T, T**A**R) and five (A**A**D, S**A**A) motifs (bold **A** refers to the deaminated adenine) as active and repressive flank-A motifs, respectively (**Fig. 4c**). To further validate that, we assessed A-to-G editing efficiency between all active or repressive flank-A motifs within our 12K TRAP sites. Our results showed that the A-to-G editing efficiency is significantly higher (p < 0.0001, fold change = 4) in the active motifs as compared to the repressive ones (**Fig. 4d, S27**).

Since the ABE editor shares principles of gRNA-guided DNA binding with SpCas9 nickase, we reasoned that many of the features (such as GC content, gRNA secondary structure (deltaG energy), N17-N20 motifs) that were known to influence SpCas9 editing efficiency should also affect ABE efficiency. To address that, we performed Pearson correlation analysis between the ABE efficiency and GC content and the deltaG energy of the guide sequence. Our results demonstrated that both GC content (**Fig. 4e, S28**) and secondary structure affect ABE efficiency (**Fig. 4f, S29**). We also investigated and validated that the presence of TT and GCC motifs at the N17-N20 seed region negatively affects ABE efficiency (**Fig. 4e, S30**).

**Figure 4.**
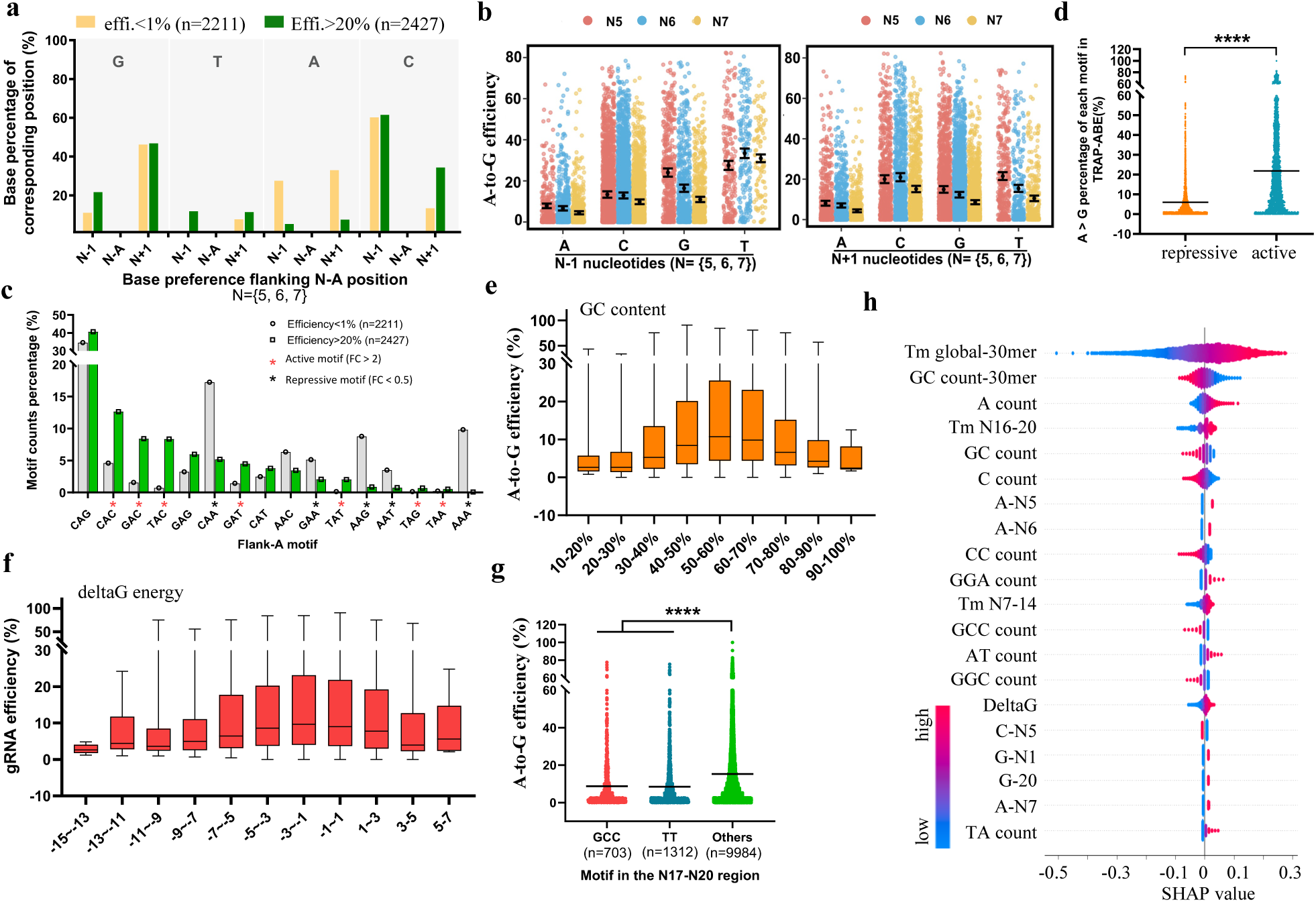
Characterization of nucleotide features affecting ABE efficiency. **a.** Proportion of frank-A (A bases located at the core editing window N5-N7) bases (A, T, C, G) between high (edited A efficiency > 20% for at least one A base within N5-N7) and low (edited A efficiency < 1% for any A base within N5-N7) efficiency ABE gRNAs. Results are based on the data from the Dox-induced HEK293T-ABE cells from day 11. **b.** A-to-G editing efficiency for A bases located at N5-N7, grouped based on the flanking bases. Results are based on the data from the Dox-induced HEK293T-ABE cells from day 11. **c.** Comparison of the frequency of N**A**N (N = A, T, C, G) trinucleotide motifs between high and low efficiency ABE gRNAs (bold A referred to the deaminated bases within the N5-N7 core editing window). Red and black asterisks indicate the active and repressive motifs based on a cutoff of two-fold difference. **d.** Scatter plot of edited A efficiency between sites within the active and repressive motifs. The number of “n” indicates number of sites. “****”, p value less than 0.0001. **e.** Box plot of overall A-to-G editing efficiency for all TRAP gRNAs according to the gRNA spacer GC content (with 10% interval). Results are based on the data from the Dox-induced HEK293T-ABE cells from day 11. Complementing results for other groups can be found in Fig. S26. **f.** Box plot of overall A-to-G editing efficiency for all TRAP gRNAs according to the gRNA spacer deltaG energy (with an interval of 2). Results are based on the data from the Dox-induced HEK293T-ABE cells from day 11. Complementing results for other groups can be found in Fig. S27. **g.** Dot plot of overall ABE efficiency between gRNAs have the GCC or TT motifs within the seed region N17-20 and gRNAs without these two motifs. “n” indicates the number of gRNAs within each group. Results were based on the data from the Dox-induced HEK293T-ABE cells from day 11. Complementing results for other groups can be found in Fig. S28. “****”, p value less than 0.0001. **h.** Top 20 features that weighted the most based on the GNL machine learning model that affect the overall ABE efficiency in cells. Results are shown as the SHAP (SHapley Additive exPlanations) values. The 30mere comprises 4bp upstream, 20bp protospacer, 3 bp PAM, and 3 bp downstream sequences. Complementary SHAP results for each edited Adenine site within the N3-N11 window were shown in Fig. S30. Results are based on the ABE editing data from the Dox-induced HEK293T-ABE cells from day 11.

### An improved machine learning model to predict ABE efficiency

We sought to apply our GNL-scorer machine learning model [43] to systematically identify features of importance for ABE efficiency (GNL-scorer_ABE) and ABE efficiency prediction rules, as well as developed a new machine learning-based tool for predicting ABE efficiency. We randomly selected 80% (the remaining 20% used for model evaluation) of the ABE efficiency data and trained the Bayesian Ridge regression (BRR)-based GNL-scorer model with 2485 features (**Fig. S31a**). The ABE efficiency prediction was performed for both A-to-G edited site efficiency in N3-N11 and the overall ABE efficiency. Our results showed that the accuracy of predicting ABE is above 60% for the core ABE editing window (**Fig. S31b**), and the accuracy of predicting the over ABE efficiency is approximately 70% (**Fig. S21**). The SHAP algorithm-based feature outputs further demonstrated that our machine learning results consistently revealed that features such as the global melting temperature, GC content, deltaG energy, and nucleotide compositions (such as the presence of Adenine in N5-N7, Guanine in N20) greatly affect the ABE efficiency (**Fig. 4f and S32**). Collectively, we demonstrate that the rich ABE editing efficiency data revealed by TRAP-seq enable us to systematically define factors influencing ABE efficiency and improve ABE gRNA design for future studies.

### Quantification of CBE-mediated recoding efficiency at 11,979 sites by TRAP-seq

After demonstrating that the TRAP-seq method is versatile for massively parallel quantification of SpCas9 and ABE efficiency, we sought to test the performance of TRAP-seq for high throughput quantification of CBE efficiency. As mentioned earlier, all the gRNA spacers of our 12K TRAP-seq library were retrieved from the iSTOP database. This allows us to address the STOP recoding efficiency of all 11,979 gRNA and 3,832 genes by CBE in cells (**Fig. 5a**). First, based the optimized and constant conditions of lentivirus library transduction, we performed 10 parallel TRAP-seq-library based CRISPR CBE editing experiments in the Dox-inducible HEK293T-CBE cells (**Fig. S33**). Constant sequencing depths (1,000X coverage) and TRAP representation (**r = 0.96-0.97**) were achieved for Dox-free and Dox-addition HEK293T-CBE cells 2 and 11 days after transduction by targeted amplicon sequencing of the TRAP region (**Fig. S34**). Next, to assess CBE editing, we quantified the efficiency of C-to-T edit, as well as C-to-G and C-to-A edits, of all C bases within the 37bp of all 11,979 TRAP sites (**Fig. 5b, Table S4**, full results were shown in the CRISPR Atlas database). As seen from the overall (**Fig. 5c, S35**) and C-to-T site (**Fig. 5d**) CBE efficiency plots, quantification of the average efficiency of all edited Cs for the 11979 gRNAs revealed that there was an even distribution of gRNAs editing efficiency from 5% to 75%. There were 371 gRNAs with very low CBE efficiency (0-5%) (**Fig. 5c**). The C-to-T CBE efficiency was primarily detected in the Dox-induced HEK293T-CBE cells at 11 days after transduction, indicating there was very minor leakiness of CBE editor expression (**Fig. 5c, S35**). Of particular note, compared to ABE, the editing window of CBE was broader (N1 to N16) but the highest cytosine editing efficiency was similarly found at N6 (**Fig. 5d, S36a**). Quantification of all edited Cs revealed that the majority (93.18%) were C-to-T edits, however, low frequency of unspecific C-to-G (4.54%) and C-to-A (2.27%) edits were also observed for the CBE (BE4-gam) base editor (**Fig. S36b**).

**Figure 5.**
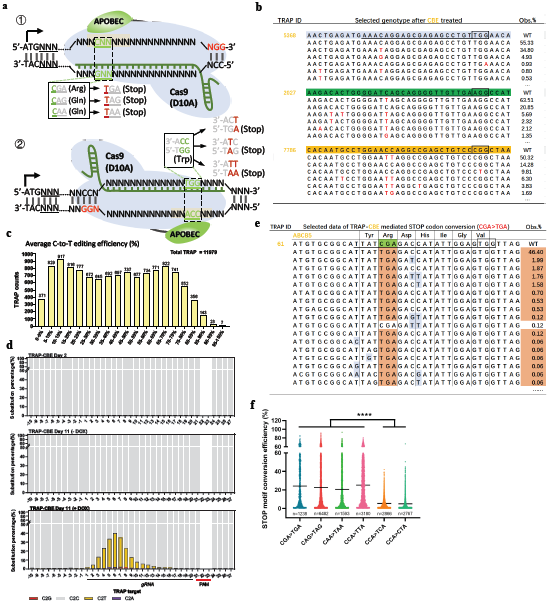
Quantification of CBE-mediated recoding efficiency at 12,000 sites by TRAP-seq. **a.** Schematic illustration of the 6 stop-codon recoding schemes by CBE. Cas9 (D10A) is the Cas9 nickase used in the testing CBE editor. The stop-codon recoding scheme is draw based on sense and anti-sense strands. “ATG”, translation start site; “NGG”, PAM of SpCas9. **b.** Representation of top 5 cytosine base editing outcomes for three TRAP sites. Full CBE frequency results can be found in the CRISPR Atlas database. **c.** Quantification of overall CBE efficiency for all gRNAs in Dox-induced HEK293T-CBE cells at 11 days. Other groups are presented in Fig. S33. **d.** Quantification of overall percentage of C-to-T, C-to-G, C-to-A, and C-to-C (unedited) events across the 37bp region of all TRAP sites. Results are based on HEK293T-CBE cells from 2 days post transduction, and transduced HEK293T- CBE cells from 11 days cultured in Dox-free or Dox-addition medium. **e.** Representation of CBE-mediated recoding efficiency at one TRAP site measured by TRAP-seq. Complementary results referred to Fig. S35 and CRISPR atlas database. **f.** Comparison of C-to-T efficiency between sites within the 6 stop-codon recoding types. “n” indicates the number of sites within each group. “****”, p value less than 0.0001.

We next investigated the stop-codon recoding efficiency by CBE in cells. With the current setups in HEK293T-CBE cells, the median STOP efficiency of all 11,979 gRNA was approximately 22% (**Fig. 5e, S37, S38a**). A total of 3,481 genes (90%) were successfully knocked out by stop-codon recoding with CBE (**Fig. 38b**). We also sought to analyze the efficiency of the 6 types of recoding into stop codons (**Fig. 5a**). As seen in **Fig. 5f**, the recoding efficiencies of C**C**A-to-CTA, **C**CA-to-TCA and **CC**A-to-TTA were significantly higher (**p value < 0.0001**) than **C**GA-to-TGA, **C**AG-to-TAG and **C**AA-to-TAA (bold **C** refers to the deaminated cytosines). Taken together, we here demonstrated that the TRAP-seq technology enables massively parallel quantification of CBE-mediated recoding capacity in cells.

### Characterization of nucleotide features affecting CBE efficiency

The large-scale CBE efficiency data from 56,887 edited Cs and 11,979 gRNAs enable us to characterize the features that affect CBE efficiency in cells. To simplify the analysis and characterization, we first selected two opposing types of CBE gRNAs based on their efficiencies: low efficiency (n = 1,844, the efficiency of any edited C within the protospacer is < 5%) and high efficiency (n = 1,731, At least one edited C efficiency within the protospacer is > 60%). Next, we compared the base preference across the 37bp TRAP regions between the two types of gRNAs. Our results clearly show that the high efficiency CBE gRNAs favor the presence of Cytosine (N4-N8), but disfavor Guanine (N3-N7) and Adenine (N5-N6) within the core editing window (**Fig. S39**). The presence of Thymine within the seed region (N8-N20) of protospacers was underrepresented in the high efficiency CBE gRNAs, which was in contrast to the N3-N6 region. Similar to our findings in ABE, high efficiency CBE gRNAs favor the presence of Guanine at proximal PAM region (N19-N21) and disfavored Cytosine at N20.

Since the core editing window of CBE is N4-N8 (**Fig. S36**), we focused on the deaminated Cytosines within N4-N8 when further analyzing the effect of flanking bases on CBE efficiency. To enable comparison, we selected C-to-T edited sites of low (< 5%, N = 4,898) and high (> 50%, N = 5,058) efficiency within the N4-N8 region. Our results show that the presence of Thymine upstream is strikingly overrepresented in the highly efficient C-to-T editing, whereas the presence of Guanine upstream is only present in the low efficiency CBE sites (**Fig. 6a**). In addition, the highly efficient CBE sites are less frequently flanked by Adenine downstream and more frequently flanked by Cytosine, as compared to low efficiency CBE sites (**Fig. 6a**). To validate this observation, we calculated the C-to-T editing efficiency at N4-N8 for all 12,000 TRAP sites. Consistent with previous observations, the overall efficiency of CBE sites flanked by Thymine upstream is approximately two-fold higher than with other flanking bases. Of note, sites flanked by Guanine upstream (as well as downstream, but to less extent) show much lower CBE efficiency (fold changes = 2 - 12 folds) (**Fig. 6b**). This provides highly valuable knowledge for designing CBE gRNAs with better editing outcome.

**Figure 6.**
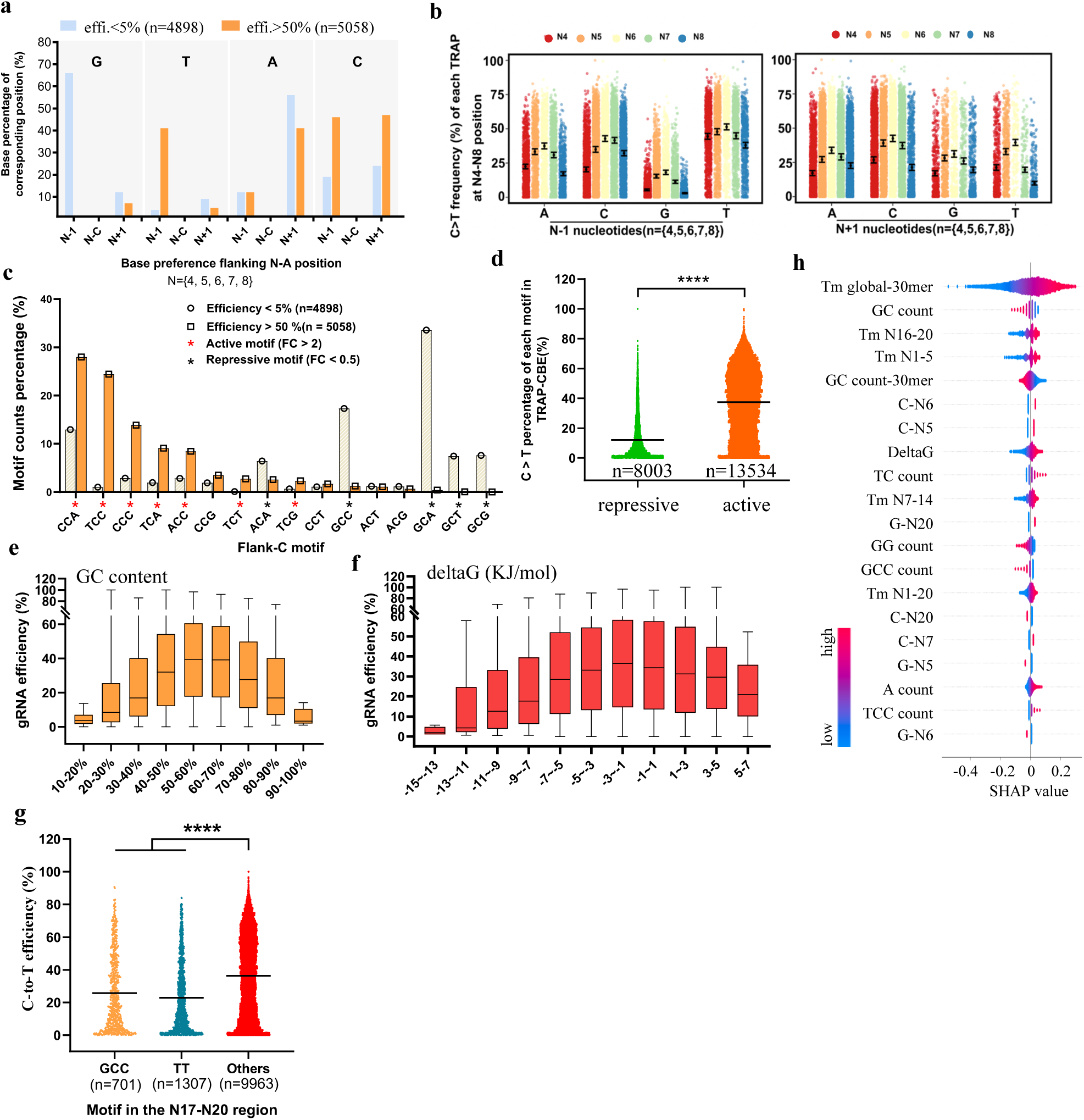
Characterization of nucleotide features affecting CBE efficiency in cells by TRAP-seq. **a.** Proportion of frank-C (N4-N8) bases (A, T, C, G) between high (at least one edited C base within N4-N8 was >50% edited) and low (edited C efficiency for any C base within N4-N8 was less than 1%) efficiency CBE gRNAs. Results are based on the data from the Dox-induced HEK293T-CBE cells from day 11. **b.** C-to-T editing efficiency for C bases located at N4-N8, grouped based on the flanking bases. Results are based on the data from the Dox-induced HEK293T-CBE cells from day 11. **c.** Comparison of the presence of the 16 N**C**N trinucleotide motifs between high and low efficiency CBE gRNAs (bold C referred to the deaminated cytosine within the N4-N8 core editing window). Red and black asterisks indicate the active and repressive motifs based on a cutoff of two-fold difference. **d.** Scatter plot of edited C efficiency between sites within the active and repressive motifs. The number of “n” indicates number of sites. “****”, p value less than 0.0001. **e.** Box plot of overall C-to-T editing efficiency for all TRAP gRNAs according to the gRNA spacer GC content (with 10% interval). Results are based on the data from the Dox-induced HEK293T-CBE cells from day 11. Complementing results for other groups can be found in Fig. S36. **f.** Box plot of overall C-to-T editing efficiency for all TRAP gRNAs according to the gRNA spacer deltaG energy (with an interval of 2). Results are based on the data from the Dox-induced HEK293T-CBE cells from day 11. Complementing results for other groups can be found in Fig. S38. **g.** Scatter plot of overall CBE efficiency between gRNAs have the GCC or TT motifs within the seed region N17-20 and gRNAs without these two motifs. “n” indicates the number of gRNAs within each group. Results were based on the data from the Dox-induced HEK293T-CBE cells from day 11. Complementing results for other groups can be found in Fig. S41. **h.** Top 20 features that weighted the most based on the GNL machine learning model, which affect the overall CBE efficiency in cells. Results are shown as the SHAP (SHapley Additive exPlanations) values. The 30mere comprises 4bp upstream, 20bp protospacer, 3 bp PAM, and 3 bp downstream sequences. Complementary SHAP results for each edited cytosine site within the protospacer N1-N20 were shown in Fig. S42. Results are based on the CBE editing data from the Dox-induced HEK293T-CBE cells from day 11.

Since the flanking bases played an important role on CBE efficiency, we reasoned that there exists a preference of tri-nucleotide flank-C motifs (N**C**N; N=A, T, C, G; bold C refers to the deaminated cytosine located at N4, N5, N6, N7 or N8) for active or repressive CBE. To identify these motifs, we compared the frequency of all 16 N**C**N motifs between the low (N=4898, efficiency < 5%) and high (N=5058, efficiency > 50%) efficiency cytosine editing sites in N4-N8. Based on a two-fold difference, we identified seven (T**C**N, C**C**M and A**C**C; N=A, T, C, G; M = A, C) and five (A**C**A and G**C**N) as active and repressive motifs, respectively (**Fig. 6c**). To validate this, we further compared the C-to-T editing efficiency between the active and repressive motifs for all 21537 edited C sites in our 12K TRAP-seq library. The C-to-T editing efficiency of the active motifs were significantly higher (fold change = 3, p < 0.0001) than those within the repressive motifs (**Fig. 6d, S40**). We also sought to investigate if the CBE efficiency shared features with SpCas9 editing efficiency using our CBE 12K TRAP-seq data. Thus, we analyzed the correlation between CBE efficiency and the GC content, the gRNA spacer secondary structure, as well as the proximal PAM motifs. Our results show that, similar to SpCas9 and ABE, the CBE efficiency is affected by the gRNA spacer GC content (**Fig. 6e, S41**) and secondary structure (**Fig. 6f, S42**). The CBE efficiency of gRNAs is significantly (**p < 0.0001**) lower with TT or GGC motifs at the proximal (17-N20) PAM region (**Fig. 6g, S43**).

### An improved machine learning model to predict CBE efficiency

We further took advantage of our GNL-scorer machine learning model to development a prediction tool and systematically evaluate the effect of 2485 features on CBE efficiency. Based on randomly selecting 80% of 12K gRNAs CBE efficiency for model training and 20% for model evaluation, our results showed that the accuracy prediction score of CBE efficiency by the GNL machine learning model reaches approximately 80% (**Fig. S21**). Apart from predicting the overall CBE efficiency, our machine learning-based prediction tool provides highly precise predicting outcome of the site-specific CBE efficiency within the editing window (**Fig. S44**). Consistently, our machine learning results further showed that features such as melting temperatures, GC content, deltaG energy, nucleotide composition (e.g. the presence of cytosine at N5-N7, nucleotide counts, motifs) greatly affect CBE efficiency (**Fig. 6h and Fig S45**). Finally, the machine learning-based CBE efficiency prediction algorism (GNL-scorer_CBE) has also been deposited to the GitHub to facilitate the design of CBE gRNAs of high efficiency. Taken together, we hereby demonstrate that high-throughput quantification of CBE efficiency by TRAP-seq enables the better understanding and design of highly efficient CBE gRNAs in cells.

### The CRISPR Atlas

As part of this work, a human CRISPR atlas database (http://www.crispratlas.com/crispr) has been launched to present and integrate all the SpCas9, ABE and CBE efficiency and editing outcomes for 12,000 gRNAs in HEK293T cells. This CRISPR atlas is presented with a gene-centered and gRNA-centered summary of the overall efficiency of gRNAs and editing outcomes. For SpCas9, total efficiency, indel (1-30bp deletions, 1-10bp insertions) profiles were presented for each gRNA. For ABE and CBE, the overall gRNA efficiency and graphical presentation of base substitution efficiency across the 37bp TRAP region were shown for all 12,000 TRAP sites. We believe that the CRISPR atlas database generated by this study will complement the existing CRISPR resources in gRNA design [44], efficiency prediction [45, 46] and indel prediction [27], thus streamlining the application of CRISPR in functional studies.

## DISCUSSION

In conclusion, the work described here demonstrates the broad applicability of the TRAP-seq system for massively parallel quantification of SpCas9, ABE and CBE efficiency in human cells. Recent studies published by other groups have demonstrated that the surrogate target sites can well mimic the SpCas9 indel outcomes at the corresponding endogenous sites, and thus predict the SpCas9 indel profile for a given gRNA in cells [26, 28, 40]. Consistent with that, we demonstrate corroborating findings with our TRAP-seq method and further expand the data of SpCas9 knockout efficiency and indel profiles with 12,000 gRNAs. This will aid the improvement of CRISPR gRNA design for gene knockout purposes with machine learning models [40, 45]. With such a large amount of SpCas9 efficiency data, it is possible to systematically identify both previously known as well as novel features that affect CRISPR gene editing efficiency. Importantly, according to our knowledge, this is the first time that both ABE and CBE efficiencies are measured at such a large scale in cells. Based on the 12K ABE and CBE TRAP-seq data, our analyses identify several novel features (such as the preference of flanking bases, active/repressive tri-nucleotide motifs) that strongly influence ABE and CBE efficiency in cells, respectively. We believe that incorporating the nucleotide features of importance for ABE and CBE efficiency from this study will improve the performance of in silico base editing designers such as BE-Designer [47] and Beditor [48]. However, we acknowledge that there might be a difference in the DNA repair machinery between different cell types and organisms, which will potentially affect the SpCas9, ABE and CBE efficiency and outcome. Additional experiments will be required to test the SpCas9, ABE and CBE efficiency with TRAP-seq in more cell lines in the future.

The concept of using surrogate target sites to capture the gene editing outcomes is highly attractive. We and other groups have generated dual-fluorescence-based surrogate systems for rapid evaluation of ZFNs, TALENs and CRISPR-Cas9 activity in cells [29, 49, 50]. The DSBs generated by CRISPR-Cas9 were predominantly repaired by the NHEJ and MMEJ pathways, which will lead to the introduction of small indels at the DBS site. However, large deletions or chromosomal rearrangements have also been reported in CRISPR editing as outcomes of repaired mediated by e.g. HDR or SSA in cells [51, 52]. The TRAP-seq system developed in this study is based on a 37bp surrogate target site. Thus, SpCas9 editing outcomes such as large deletions or chromosomal arrangements will not be captured by our method. However, for ABE and CBE, the editing outcomes would not be affected by such a size-related problem.

Earlier, we have discovered that chromatin accessibility at the editing sites affect CRISPR gene editing efficiency [24]. Since the TRAP-seq library were randomly inserted in the genome of the targeted cells, the chromatin accessibility state of the surrogate site might be different from the endogenous target site. It would be interesting to apply the TRAP-seq system to systematically analyze the epigenetic factors (e.g. DNA methylation, chromatic accessibility) on ABE, CBE efficiency in future studies.

In this study, we demonstrate the TRAP-seq system with applications in massively parallel quantification of editing efficiency for SpCas9 and base editors (ABE and CBE) derived from the SpCas9. The current CRISPR-based gene editing toolbox has been greatly expanded with the engineered SpCas9 (e.g. xCas9, eSpCas9, SpCas9-HF1), the SpCas9 orthologs (e.g. SaCas9, StCas9, NmCas9) and other Cas proteins (e.g. Cas12a) [53-55]. However, features affecting the editing efficiency and indel outcomes are still rarely explored for most of these Cas proteins, which will limit the applications of this great toolbox. We believe that the TRAP-seq will become an important technology for the whole CRISPR gene editing society to better understand how CRISPR gene editing works in cells. The CRISPR atlas database generated by this study will become a CRISPR-centered portal, in which we provide experimentally validated gRNAs for CRISPR gene editing. Taken together, the TRAP-seq technology, the SpCas9/ABE/CBE efficiency of 12,000 gRNAs, and the CRISPR atlas database will enable us to better functionally understand how CRISPR works in cells and improve CRISPR in both research, therapeutic and drug discovery applications.

## MATERIALs and METHODS

### Vector construction

The empty pLenti-TRAP-seq vector backbone (shown in **Fig. S1**) was generated by a serial of cloning. Briefly, we replaced the SpCas9 open reading frame (ORF) in pLentiCRISPRv2-puro (Addgene plasmid # 98290) plasmid with an enhance green fluorescence protein (EGFP) ORF based on *XbaI* and *BmHI* digestion and T4 ligation. Next, we replaced the gRNA cassette in the EGFP-inserted pLentiCRISPRv2-puro with a synthetic Golden-Gate Assembly cassette, and hence generated a lentivirus-based vector (hereafter referred as pLenti-TRAP-seq) allowing the insertion of TRAP DNA to the Golden-Gate cloning site by GGA. The full sequencing of the pLenti-TRAP-seq vector can be downloaded from our CRISPR atlas database (www.crispratlas.com/crispr). Original plasmid stock can be acquired from the corresponding authors’ lab.

The doxycycline inducible SpCas9, ABE, and CBE vectors were generated by subcloning. Briefly, based on a PiggyBac transposon system (full GenBank vector sequences can be found at the CRISPR atlas website), which consists of an all-in-one expression system: (1) An expression cassette of a TRE promote-driven protein expression cassette with multiple cloning sites (MCS). (2) An expression cassette of a consecutive promoter-driven Tetracycline-Controlled Transcriptional Activation and hygromycin. The coding sequences of SpCas9 (Addgene plasmid ID # 41815), ABE 7.10 (Addgene plasmid ID # 102919), and CBE (Addgene plasmid ID # 100806) were PCR amplified and inserted to the MCS of the PiggyBac transposon system. All vectors were validated by Sanger sequencing.

### TRAP 12K oligos design and microarray synthesis

A typical TRAP oligo consists of the BsmBI recognition site “**cgtctc**” with 4 bp specific nucleotides “acca” upstream, following the GGA cloning linker “aCACC”, one bp “g” for initiating transcription, then the 20 bp gRNA sequences of “gN20”, 82bp gRNA scaffold sequence, 37 bp surrogate target sequences (10bp upstream sequences, 23 bp protospacer and PAM sequences, 4 bp downstream sequence), the downstream linker “GTTTg” and another BsmBI binding site and its downstream flanking sequences “acgg”. An example of the typical TRAP oligo sequence was shown below: “acca**cgtctc**aCACCgGTCCCCTCCACCCCACAGTGGTTTTAGAGCTAGAAATAGCAAGTTA AAATAAGGCTAGTCCGTTATCAACTTGAAAAAGTGGCACCGAGTCGGTGCTTTTTT<u>ACT</u> <u>TTTATCTGTCCCCTCCACCCCACAGTGGGGCCAC</u>GTTTg**gagacg**acgg”. Sequences in the black frame is the 20 bp gRNA spacer. The underline sequence is the 37 bp surrogate target site, which is termed as “TRAP target” shortly in this study.

For the 12K TRAP oligo design, we used bioinformatic tools to automatically generate the 12K TRAP oligo pools. Briefly, 1) we selected approximately 7,000 genes from the a drugable gene database (http://dgidb.org); 2) Discard all the exons which the DNA length was less than 23 bp with filtering; 3) Select the first three coding exons of each gene. If the exons number is less than 3, keep all the exons; 4) Extract all the possible gRNA sequences (including the PAM sequence “NGG”) in these filtered exons sequence, analyzes and predictd the off-target sites of each gRNA using FlashFry version 1.80 (https://github.com/mckennalab/FlashFry), discarded gRNAs with potential off-target of 0-3 bp mismatches in human genome; 5) Rank each gRNA based on the number of off-target site in an ascending order; 6) Map and extract the 10 bp upstream and 4 bp downstream flanking sequence of each selected gRNA, construct the TRAP target sequence as 10 bp upstream + 23 bp gRNA (include PAM) + 4 bp downstream = 37 bp; 7) Filter out TRAPs with BsmBI recognition site, because of GGA cloning; 8) Compared all the selected gRNAs with the database of CRISPR-iSTOP [56]; 9) Construct the full length sequence of each TRAP, which is 170 bp; In total, the first 12K TRAP-seq oligos contain 3832 genes and 12000 TRAPs were contained in the final TRAP-library. The 12K oligo pools was synthesized in Genscript® (Nanjing, China).

### TRAP-12K plasmid library preparation

First, the TRAP 12K oligos were cleaved and harvested from the microarray and diluted to 1 ng/*μ*L. Next, we performed PCR amplifications using the primers: TRAP-oligo (BsmBI GGA)-F: 5’-TACAGCTaccacgtctcaCACC-3’; TRAP-oligo (BsmBI GGA)-R: 5’-AGCACAAccgtcgtctccAAAC-3’.

The PCR reaction was carried out using PrimeSTAR HS DNA Polymerase (Takara, Japan) following the manufacturer’s instruction. Briefly, each PCR reaction contained 1 *μ*L oligo template, 0.2 *μ*L PrimeSTAR polymerase, 1.6 *μ*L dNTP mixture, 4 *μ*L PrimeSTAR buffer, 1 *μ*L forward primer (10 uM) and 1 *μ*L reverse primer (10 uM) and ddH2O to a final volume of 20 *μ*L.

The thermocycle program was 98°C 2min, (98°C/10s,55°C*/10s,72°C/30s) with 21 cycles, then 72°C for 7min and 4°C hold. To avoid amplification bias of oligos introduced by PCR, we conducted gradient thermocycles and performed PCR products gray-intensity analysis to determine the optimal PCR cycles of 21. The best thermocycles should be in the middle of an amplification curve. In this study, the *PCR cycles* was 21 for oligos amplification. But for PCR amplification of TRAP from cells integrated with TRAP lentivirus, the *PCR cycle* was 25. The final TRAP PCR product length was 184 bp. We performed **72** × parallel PCR reactions for 12K oligos amplification, then these PCR products were pooled and gel purified by 2% agarose gel. 1 *μ*g purified PCR product were quantified with PCR-free next generation sequencing (MGI Tech).

The PCR products of TRAP oligos were then used for Golden Gate Assembly (GGA) to generate the TRAP 12K plasmids library. For each GGA reaction, the reaction mixture contained 100 ng pLenti-TRAP-seq vector, 10 ng purified 12K TRAP oligos-PCR products, 1 *μ*L T4 ligase (NEB), 2 *μ*L T4 ligase buffer (NEB), 1 *μ*L BsmBI restriction enzyme (ThermoFisher Scientific, FastDigestion) and ddH2O to a final volume of 20 *μ*L. The GGA reactions were performed at 37°C 5 min and 22°C 10 min for 10 cycles, then 37°C 30 min and 75°C 15 min. **36** × parallel GGA reactions were performed and the ligation products were pooled into one tube.

Transformation was then carried out using home-made chemically competent DH5a cells. For each reaction, 10 *μ*L GGA ligation product was transformed in to 50 *μ*L competent cells and all the transformed cells were spread on one LB plate (15 cm dish in diameter) with Xgal, IPTG and Amp selection. High ligation efficiency was determined by the presence of very few blue colonies (also see **Fig. S4**). To ensure that there is sufficient coverage of each TRAP in the 12K TRAP-seq library, **42** × parallel transformations were performed and all the bacterial colonies were scraped off and pooled together for plasmids midi-prep. For NGS-based quality quantification of TRAP coverage, midi-prep plasmids were used as DNA templates for TRAP PCR amplifications, followed by gel purification and NGS sequencing.

### TRAP-12K lentivirus packaging

HEK293T cells were used for lentivirus package. All the cells were cultured in Dulbecco’s modified Eagle’s medium (DMEM) (LONZA) supplemented with 10 % fetal bovine serum (FBS) (Gibco), 1% GlutaMAX (Gibco), and penicillin/streptomycin (100 units penicillin and 0.1 mg streptomycin/mL) (The culture medium was named as D10 shortly) in a 37 °C incubator with 5% CO2 atmosphere and maximum humidity. Cells were passaged every 2-3 days when the confluence was approximately 80-90%.

For lentivirus packaging: **Day 1**: Wild-type HEK293T cells were seeded to a 10 cm culture dish, 4 × 10^6^ cells per dish (10 dishes in total); **Day 2**: Transfection. Briefly, we refreshed the medium with 7 mL fresh culture medium to 1 hour before transfection (be gently, as the HEK293T cells are easy to be detached from the bottom of dish); Next, we performed transfection with the PEI 40000 transfection method. For 10 cm dish transfection, the DNA/PEI mixture contains 13 *μ*g pLenti-TRAPseq 12K vectors, 3 *μ*g pRSV-REV, 3.75 *μ*g pMD.2G, 13 *μ*g pMDGP-Lg/p-RRE, 100 *μ*L PEI 40000 solution (1 *μ*g/ *μ*L in sterilized ddH2O) and supplemented by serum-free opti MEM without phenol red (Invitrogen) to a final volume of 1 mL. The transfection mixture was pipetted up and down several times gently, then kept at room temperature (RT) for 20 min, then added into cells in a dropwise manner and mix by swirling gently. **Day 3**: Changed to fresh medium; **Day 4**: Harvest and filter all the culture medium of the 10 cm dish through a 0.45 *μ*m filter, pool the filtered media into one bottle. Each 10 cm dish generated approximately 7∼8 mL lentivirus crude. Add polybrene solution (Sigma-Aldrich) in to the crude virus to a final concentration of 8 *μ*g/mL. Aliquot the crude virus into 15 mL tubes (5 mL/tube) and store in -80 °C freezer.

### Lentivirus titer quantification by flow cytometry (FCM)

As the pLenti-TRAP-seq vector expresses a EGFP gene, the functional titer of our lentivirus prep was assayed by FCM as described previously [57]. Briefly, 1) **DAY 1**: split and seed HEK293T cells to 24-well plate, 5 × 10^4^ cells per well. Generally, 18 wells were used to perform the titter detection, a gradient volume of the crude lentivirus was added into the cells and each volume was tested by replicates. In this experiment, the crude virus gradients were 2.5 *μ*L, 5 *μ*L, 10 *μ*L, 20 *μ*L, 40 *μ*L, 80 *μ*L, 160 *μ*L and 320 *μ*L for each well. Another 2 wells of cells were used for cell counting before transduction; 2) **DAY 2**: Conduct lentivirus transduction when cells reach up to 60∼80% confluence. Before transduction, detach the last two wells of cells using 0.05% EDTA-Trypsin to determine the total number of cells in one well (N_(initial)_). Then change the remaining wells with fresh culture medium containing 8 *μ*g/mL polybrene, then add the gradient volume of crude virus into each well and swirling gently to mix; 3) **DAY 3**: Change to fresh medium without polybrene; 4) **DAY 4**: Harvest all the cells and wash them twice in PBS. Fix the cells in 4% formalin solution at RT for 20 min, then spin down the cell pellet at 2,000 rpm for 5 min. Discard the supernatant and re-suspend the cell pellet carefully in 600 *μ*L PBS, and conduct FCM analysis immediately. FCM was performed using a BD LSRFortessa™cell analyzer with at least 30,000 events collected for each sample in replicates.

The FCM output data was analyzed by the software Flowjo vX.0.7. Percentage of GFP-positive cells was calculated as: 𝒴% = N _(GFP-positive cells)_ / N _(total cells)_ × 100 %. Calculate the GFP percentage of all samples. For accurate titter determination, there should be a linear relationship between the GFP positive percentages and crude volume. The titter (Transducing Units (TU/mL) calculation according to this formula: TU/mL = (N_(initial)_ × 𝒴% × 1000)/ V. V represents the crude volume (*μ*L) used for initial transduction.

### Generation of Doxycycline-inducible spCas9/ABE7.10/CBE stable cell lines

TRE-spCas9, TRE-ABE7.10 and TRE-CBE stable cells were generated by PiggyBac transposon systems. For stable cell lines establishment, HEK293T cells were transfected with pPB-TRE-spCas9-Hygromycin (or pPB-ABE7.10-hygromycin, pPB-CBE-hygromycin) vector and pCMV-hybase with a 9:1 ratio. Briefly, 1 × 10^5^ HEK293T cells were seeded in 24-well plate and transfections were conducted 24 h later using lipofectamine 2000 reagent following the manufacturer’s instruction. Briefly, 450 ng pPB-TRE-spCas9-Hygromycin (or pPB-ABE7.10-hygromycin, pPB-CBE-hygromycin) vectors and 50 ng pCMV-hybase were mixed in 25 *μ*L optiMEM (tube A), then 1.5 *μ*L lipofectamine 2000 reagent was added in another 25 *μ*L optiMEM and mix gently (tube B). Incubate tube A and B at RT for 5 min, then add solution A into B gently and allow the mixture incubating at RT for 15 min. Add the AB mixture into cells evenly in a dropwise manner. Cells transfected with pUC19 were acted as negative control. Culture medium was changed to selection medium with 50 *μ*g/mL hygromycin 48h after transfection. Completion of selection took approximately 5-7 days until the negative cells were all dead in the un-transfected cells. The cells were allowed to grow in 50 *μ*g/mL hygromycin containing D10 medium for 3-5 days for further expansion. PCR-based genotyping were carry out using the primers: ***spCas9-iden-F***: gacacctacgatgatgatctcg; ***spCas9-iden***-***R***: tggtgctcatcatagcgcttga; ***ABE7.10-iden-F***: 5’-cagtactcgtgctcaacaatcg-3’; ***ABE7.10-iden-R***: 5’-ggcgttgcgaacaccgaataca-3’; ***BE4-iden-F***: 5’-ttcttcgatccgagagagctcc-3’; ***BE4-iden-R***: 5’-ctgcaccttgtgttcggacag-3’.

For functional tests of the spCas9, ABE7.10 and CBE4 expression cells, individual TRAP constructs packaged in lentivirus particles were transduced into the cells. Transduced cells were harvested for indel analysis 6 days after transduction. Indel analysis were carried out for both the TRAP site and the endogenous genome target sites.

### 12K TRAP-seq library lentivirus transduction

HEK293T-SpCas9, -ABE7.10 and -CBE4 cells were cultured in D10 medium with 50 *μ*g/mL hygromycin throughout the whole experiment. For 12K TRAP-seq library transduction, we followed the procedures showed in **Fig. S7, S21, S31**. Briefly, 1) Day -1: 2.5 × 10^6^ cells per 10 cm dish were seeded, and 12 dishes in total. For each group, one dish was used for cell number determination before transduction and one dish for drug-resistance (puromycin) test control and the remaining 10 dishes were used for the 12K TRAP-seq lentivirus library transduction; 2) Day 0: We determined the approximate cell number per dish by cell countering. This was used to determine the volume of crude lentivirus used for transduction using a multiplicity of infection (MOI) of 0.3. The low MOI (0.3) ensures that most infected cells receive only 1 copy of the lentivirus construct with high probability [34]. The calculation formula is: V = N × 0.3 / TU. V = volume of lentivirus crude used for infection (mL); N = cell number in the dish before infection; TU = the titter of lentivirus crude (IFU/mL). In this study, take the TRE-ABE7.10 group for instance, there were 1.875 × 10 ^7^ cells in one dish, the TRAP 12K lentivirus crude titter = 3.8 × 10 ^6^ IFU/mL. Thus V = 1.875 × 10 ^7^ × 0.3 / 3.8 × 10 ^6^ = 1.48 mL. The 12K TRAP-seq transduction coverage per dish is 1.875 × 10 ^7^ × 0.3 / 12000 = 469 x. As we performed 10 replicates for each group, the overall coverage would reach to about 4690 x. In this study, V_spCas9_ = 1.26 mL, V_ABE7.10_ = 1.48 mL, V_wt_= 1.37 mL for each dish. For transduction, we added aforementioned volume of crude virus to each group in a dropwise manner and mix by swirling gently. The infected cells were cultured in a 37 °C incubator; 3) Day 1: 24 hours after transduction, split the transduced cells of each dish to 3 dishes equally; 4) Day 2: For the 3 dishes of split (30 dishes in total, 3 divided into sub-groups), sub-group 1 (10 dishes) were harvested and labeled as the Day 2 after 12K TRAP-seq transduction. All cells from this sub-group were pooled into one tube and stored in -20 °C freezer for genomic DNA extraction; The sub-group 2 (10 dishes) was changed to fresh D10 medium contains 50 *μ*g/mL hygromycin + 1 *μ*g/mL puromycin (Dox-free group); The sub-group 3 (10 dishes) was changed to D10 medium contains 50 *μ*g/mL hygromycin + 1 *μ*g/mL puromycin + 1 *μ*g /mL doxycycline (Dox-induction group). For the WT HEK293T cells (Group 3) screening, hygromycin but not puromycin should be excluded from the culture medium; 5) The transduced cells were spitted every 2∼3 days when cell confluence reaches up to 90%. At the indicated time points in Fig. S7, 21, 31, cells were harvested and stored in -20 °C freezer for further genomic DNA extraction.

### PCR amplicons of TRAPs from cells

Genomic DNA was extracted using the phenol-chloroform method. The genomic DNA were digested with RNase A (OMEGA) to remove RNA contamination (In this study, 50 *μ*g RNase A worked well to digest the RNA contamination in 100∼200 *μ*g genomic DNA after incubating in 37 °C for 30 min). Then the genomic DNA was purified and subjected to PCR for amplification of the TRAP DNA. The PCR primers were: **TRAP-NGS-F1**: 5’-GGACTATCATATGCTTACCGTA-3’ and **TRAP-NGS-R1**: 5’-ACTCCTTTCAAGACCTAGCTAG-3’. The PCR product length is 252 bp. In this study, 5 *μ*g genomic DNA was used as temperate in one PCR reaction which contained approximately 7.6 × 10^5^ copies of TRAP construct (assuming 1 × 10^6^ cells contain 6.6 *μ*g genomic DNA), which covered about 63 × coverage of the 12K TRAP-seq library. In total, 32 × parallel PCR reactions were performed to achieve approximately 2,016 × coverage of each TRAP construct. For each PCR reaction, briefly, 50 *μ*L PCR reaction system consists of 5 *μ*g genomic DNA, 0.5 *μ*L PrimeSTAR polymerase, 4 *μ*L dNTP mixture, 10 *μ*L PrimeSTAR buffer, 2.5 *μ*L forward primer (10 uM) and 2.5 *μ*L reverse primer (10 uM) and supplemented with ddH2O to a final volume of 50 *μ*L. The thermocycle program was 98°C 2min, (98°C for 10s,55°C for 10s,72°C for 30s) with 25 cycles, then 72°C for 7min and 4°C hold. Then purify all the PCR products by 2% gel, pool the products together and conduct deep amplicon sequencing.

### Deep amplicon sequencing

MGISEQ-500 (MGI of BGI in China) was used to perform the amplicons deep sequencing following the standard operation protocol. First, PCR-free library was prepared using MGIeasy FS PCR-free DNA library Prep kit following the manufacturer’s instruction. Briefly, measure the concentration of purified PCR products using Qubit 4™ fluorometer (Invitrogen) and dilute the concentration of each sample to 10 ng/ *μ*L. 10 *μ*L diluted PCR product was mixed with an A-Tailing reaction which contained A-Tailing enzyme and buffer, incubated at 37°C for 30 minutes then 65°C for 15 min to inactive the enzyme. Then the A-Tailed sample was mixed with PCR Free index adapters (MGI.), T4 DNA Ligase and T4 ligase buffer to add index adapter at both 3’ and 5’ ends of PCR products. The reaction was incubated at 23°C for 30 min and then purified with XP beads. Then denature the PCR products to be single-strand DNA (ssDNA) by incubating at 95 °C for 3 min and keep on 4 °C for the subsequent step. Transform the ssDNA to be circles using cyclase (MGI) at 37 °C for 30 min and then digested to remove linear DNA using Exo enzyme at 37 °C for 30 min. Purify the products again by XP beads and assay the concentration of library by Qubit 4™ fluorometer. The amplicons libraries were subjected to deep sequencing on the MGISEQ-2000 platform. In this study, for each lane 4 samples (6 ng each) were pooled together for deep sequencing. To avoid sequencing bias induced by base unbalance of TRAP sequence, 12 ng whole-genome DNA library (balance library) was mixed with the 4 PCR samples in a final concentration of 1.5 ng/ *μ*L and sequenced in one lane. All the samples were subjected to pair-ended 150 bp deep-sequencing on MGISEQ-500 platform.

### Data analysis

In order to evaluate the sequencing quality of amplicons and filter the low-quality sequencing data, the default parameters of Fastqc-0.11.3 (http://www.bioinformatics.babraham.ac.uk/projects/fastqc/) and fastp-0.19.6 (https://github.com/OpenGene/fastp) were used to carry out the filtration procedure and generate the clean dataset of each sample. The clean sequencing segments of pair-ended TRAP segments were merged using FLASh-1.2.1 (http://ccb.jhu.edu/software/FLASH/index.shtml) to obtain full-length TRAP constructs. The expression characteristics of all the sequences were analyzed by python 3.6, and most of the BsmBI linker fragments changed in orientation (GTTTGGAG-> GTTTGAAT). Therefore, in order to obtain the amplified fragment reads of each TRAP reference sequence, the TRAP sequence BsmBI Linker was removed from the reference sequence. The BWA-MEM algorithm of bwa(http://bio-bwa.sourceforge.net/) was used for local alignment, and the reads of all samples were divided into 12,000 independent libraries. Due to the existence of sequencing or synthesis introduced errors, each library was then filtered. In order to simplify the filtering process, the filtration strategy varies from TRAP-ABE7.10, TRAP-CBE4.0-gam to TRAP-SpCas9. For ABE7.10 and CBE4.0-gam, they mainly cause single-base variation, rarely introduce insertion and deletion, the trap sequence length remains unchanged before and after editing. Therefore, we filter the sequence of each library by locking the intermediate 37bp sequence starting with gRNA + scaffold fragment and ending with GTTT. While TRAP-SpCas9 mainly cause insertion and deletion, the length of trap sequence change around 37bp. Therefore, we adopt three steps to filter the sequence of each library. The first step is to obtain the sequence containing gRNA + scaffold fragment as dataset1. The second step is to obtain the sequence containing GTTTGAAT in dataset1 as dataset2. The third step is to extract the intermediate trap sequence from dataset2, which removed the length limit. In order to eliminate the interference of background noise before analyzing editing efficiency, all mutations or indels found in WT HEK293T cells group were removed from the Dox group in advance. For the TRAP-ABE7.10 and TRAP-CBE4.0-gam, the total editing efficiency for each trap is calculated according to the following formula:

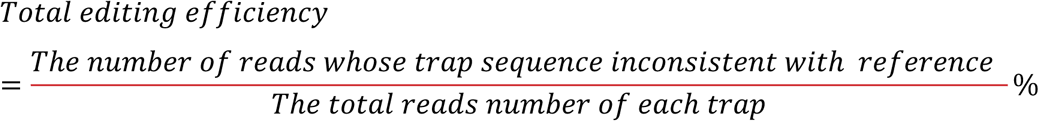

and the substitution percentage in the 37bp editing window is calculated according to the following formula:

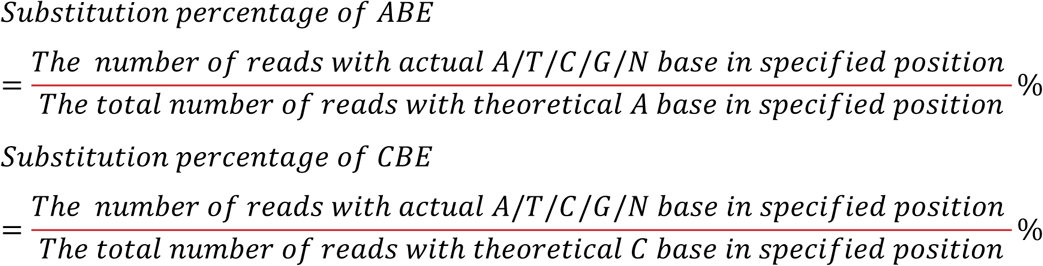

For the TRAP-SpCas9 system, the total editing efficiency for each trap is calculated according to the following formula:

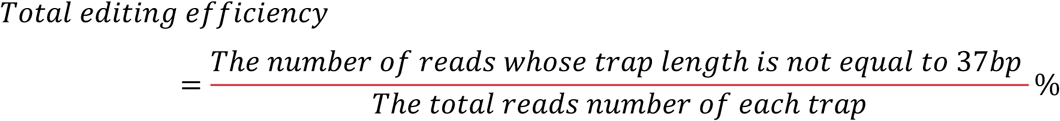

and the average fraction of indels from 30bp deletion to 10bp insertion is calculated according to the following formula:

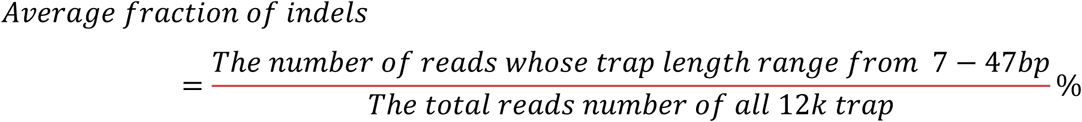

Example of selecting low and high efficency gRNA for ABE and CBE. At least one site in this sequence has more than 20% efficient for each TRAP, it is considered that the whole sequence has more than 20% analytical value. Firstly, for the range of N1-N20, divide all TRAP library into a group with an efficiency of greater than 20% and others with an efficiency of 1%, compare the base distribution of the two groups. Then for the range of N5-N7, divide all TRAP library into the above two groups, calculating the base mutation preference of *N-1* & *N + 1* sites, motif preference and the editing efficiency of N17-N20 sites including GCC, TT and other motif, respectively. For the CBE system, the statistical method is basically the same as that of ABE. Due to the high editing efficiency of CBE, in order to correct statistical deviations, CBE divides the efficiency greater than 50% and less than 5% into two groups when calculating the N1-N20 base preference and base mutation preference of *N-1 & N+1* site from N4-N8. For the editing window is wider, the efficiency greater than 50% and less than 5% is divided into two groups during the motif preference in N4-N8 window and the editing efficiency of N17-N20 sites including GCC, TT and other motif. Python-3.6 and R scripts were used for efficiency and motif analysis of all the TRAP samples. All visualizations use GraphPad Prism8.2 and R package ggplot2.

### GNL machine learning featurization

The feature set applied in our model construction contains 2485 features, which includes following five categories. (*i*) 604 features of “one-hot” encoding of the nucleotide. There are two subsets in this category: position-dependent and position-independent. And each category applies to the one nucleotide and pairwise nucleotide. Such as “_nuc_pd_Order2” consisted of e.g. AA_1/AT_1/AG_1, and “_nuc_pd_Order1” consisted of e.g. A_1/T_1/G_1/C_1. (*ii*) 3 GC features, which consists of GC count, GC count < 10, GC count > 10. (iii) 16 features of the two nucleotides flanking the NGG PAM in the 5’ and 3’. (iv) Five thermodynamic features. We calculated five thermodynamic features ausing the “Tm_staluc function” in Biopython package. All these features above were derived from the 30mer of target sequences. (v) 1856 features of three nucleotides with “position-dependent” and “position-independent”, such as ACG/AGG and ACG_1/ACT_2. Note that, all these features were encoded by 30mer context sequence.(vi) Free energy (DeltaG), which was calculated by the local version of “mfold” (http://unafold.rna.albany.edu/?q=mfold/download-mfold). We used the binary programme “quikfold” in the same bin file of “mfold” to calculate multiple input sequences in the same time. All the parameters were set as default, to make sure the outputs of each 20mer sequences be the same as the webpage.

### Model training: Comparison with other machine learning models

We trained model to predict the cleavage efficiency of each site in the editing windows among ABE and CBE editing system. For the whole sequence sites of gRNA, three type of editing system was uniformly trained by the same BRR model. To select the optimized predictive model, we initially compared the predictive performance among eight models, they are, Bayesian Ridge regression (BRR), gradient boosted regression tree (GBRT), decision tree (DT), L1-regression (L1-reg), L2-regression (L2-reg), linear regression (LR), neural network (NN), random forest (RF). All these algorithms were trained under the same features vector spaces. The mean performance of each algorithm was conducted by 10-fold cross validation. Because all of them can be used as regression, the performance was evaluated by SCC (Spearman Correlation coefficient). The model with best performance, highest SCC value and lowest S.D., was selected. Finally, BRR was outperform than other counterparts. All the models applied for training using the scikit-learn package in python. During training, optimal hyper-parameter was chose using the inner 10-fold cross validation, using the grid search. After that, α and λ were both set as 1.e-6.

### Bayesian Ridge Regression

Bayesian Ridge regression is changed from Bayesian linear regression by adding the prior of coefficient “ω” as spherical Gaussian and the priors over lambda are chosen to be gamma distributions, which is similar to the classical Ridge regression [58]. Bayesian linear regression is briefly shown as (1), and the coefficients of w is hypothesized as the spherical distribution to find a maximum posteriori estimation of ω as (2) shows.

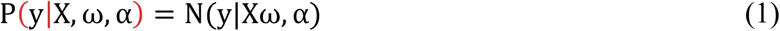

Where α is treated as a random variable that is to be estimated from the data as gamma distribution.

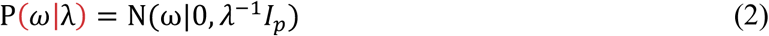

Where λ is also treated as a random variable that is to be estimated from the data, and also be hypothesized as gamma distribution.

### Model explanation

In addition to train the model with high performance, we also interested in the model importance for our final model. We used **SHAP (SHapley Additive exPlanations) algorithm [59], which** is a unified approach to explain the output of any machine learning model. Importantly, we excluded the necessary site when training the site model for each editing system. E.g. We drop A1 when training the N1 site of ABE system. So, the importance of the left features can be ranked by the SHAP value, and the top 20 important features of each model were shown for each editor.

## DATA AVAILABILITY

NGS data: CNBG accession number: TBD

Code for machine learning: https://github.com/TerminatorJ/CRISPR-TRAP-seq.git

CRISPR atlas: www.crispratlas.com/crispr

## ACKNOWLEDGEMENT

This project is supported by the Sanming Project of Medicine in Shenzhen (SZSM201612074), Qingdao-Europe Advanced Institute for Life Sciences Grant, Guangdong Provincial Key Laboratory of Genome Read and Write (No. 2017B030301011) and Guangdong Provincial Academician Workstation of BGI Synthetic Genomics(No. 2017B090904014). Y.L. is supported by BGI-Research and Brainstem Center of Excellence (Danish Innovation Fund, BrainStem). L.L. is supported by the Lundbeck Foundation (R219–2016-1375) and the DFF Sapere Aude Starting grant (8048-00072A). G.C. is supported by National Human Genome Research Institute of the National Institutes of Health [RM1HG008525]. We thank the China National GeneBank for the support of executing the project under the framework of Genome Read and Write.

## AUTHOR CONTRIBUTION

L.L. and Y.L. conceived the idea. L.B, G.C, L.L and Y.L oversaw the whole study. X.X, K.Q, L.L. and Y.L. and designed the study. X.X, K.Q, X.L, and X.P performed most of the experiments and analyses. All authors have contributed to the execution of the experiments and studies. L.L. and Y.L. drafted the manuscript. All authors discussed the results and contributed to the final manuscript.

