## supplemental files for "Massively parallel quantification of CRISPR editing in cells by TRAP-seq enables better design of Cas9, ABE, CBE gRNAs of high efficiency and accuracy"

**This document includes**

**1. Supplementary Figure Legends**

**2. Supplementary Figures**

#### SUPPLEMENTARY FIGURE LEGENDS

**Figure S1. Schematic illustration of the TRAP-seq vector's core gene elements**
Golden-Gate Assembly is based on the BsmBI restriction enzyme. Insertion of each TRAP DNA into the GGA site will replace the lacZ expression cassette by 139bp TRAP DNA.

**Figure S2. Validation of SpCas9, ABE and CEB efficiency and indel outcome in three TRAP and their corresponding endogenous genome sites**
(a, b) Comparison of indel profiles (a) and efficiency (b) between the TRAP and endogenous genome site *AAVS1*.
(c, d) Comparison of ABE profiles (c) and efficiency (d) between the TRAP and endogenous genome site *INHBC*.
(e, f) Comparison of CBE profiles (e) and efficiency (f) between the TRAP and endogenous genome site *TYMP*.

**Figure S3. Optimization of protocols for amplifying the 12K TRAP-seq library by PCRs**
**a.** Procedure of PCR amplification, QC by next generation sequencing, Golden Gate Assembly and vector prep and QCs.
**b.** Gel electrophoresis results for optimizing the PCR cycle numbers for TRAP DNA amplification.
**c.** Quantification of gel intensity in S3b by Image J.

**Figure S4. Highly efficient cloning by Golden-Gate Assembly**
Representative images of the LB plates with the majority of colonies as while ones, which indicate that the LacZ cassette in the empty pLenti-TRAP-seq vector has been replaced. Only very few blue colonies appeared in the whole plate, suggesting high cloning efficiency.

**Figure S5. Measurement of lentivirus titer by flowcytometry**
Two independent lentivirus preps were generated for the 12K TRAP-seq library and the functional transduction unit was measured using fluorescence activated cell sorting. The 12K TRAP-seq lentivirus prep 1 was selected and used for the whole study. Value were given as mean and S.D. n = 2.

**Figure S6. Evaluation of sequencing coverage of the 12K TRAP DNAs in the synthetic oligo pools, GGA plasmids, and WT HEK293T cells transduced with the 12K TRAP-seq lentivirus library.**

**a.** Ranked dot plot the sequencing depth (read counts) of each TRAP DNA.

**b**. Pearson correlation of the TRAP coverages among oligos, plasmids, and transduced cells.

**Figure S7. Schematic illustration of massively parallel quantification of 12,000 gRNAs efficiency in the HEK293T-SpCas9 cells.**

MOI, multiplexity of infection; The day of transduction was defined as Day 1. As the HEK293T-SpCas9 cells grew very rapidly, transduced cells were passaged every second day. Days of harvesting the cells for DNA extraction were indicated.

**Figure S8. Enrichment of transduced cells by puromycin selection**

Representative images of transduced cells captured under normal light filter and filter for the EGFP fluorescent. Cells were from the through groups as indicated in Fig. S7. Magnification, 40X.

**Figure S9. Evaluation 12K TRAP DNA coverage and representation in the transduced cells by targeted amplicon sequencing**

**a.** Quantification of TRAP coverage by violin plot. Values were based on log 1P (read+1) of each TRAP DNA across all groups

**b.** Pearson correlation analysis for each trap between WT cells and SpCas9 expressing cells (with or without Dox induction)

**Figure S10. Distribution of gRNAs according to SpCas9 efficiencies**

**a.** Histogram plot overview of the 11,936 gRNAs efficiency detected by targeted amplicon sequencing of the transduced HEK293T-SpCas9 cells from Day 8, cultured in Dox-free medium.

**b.** Histogram plot overview of the 11,960 gRNAs efficiency detected by targeted amplicon sequencing of the transduced HEK293T-SpCas9 cells from Day 10, cultured in Dox-free medium. Corresponding results for Dox-induction groups were shown in Fig. 1d.

**Figure S11. Distribution of indel profiles introduced to all the TRAP sites by SpCas9**

**a.** Summary of indels (1-30bp deletion, 1-10bp insertion) for the 11,936 TRAP sites in the transduced HEK293T-SpCas9 cells from Day 8, cultured in Dox-free medium.

**b.** Summary of indels (1-30bp deletion, 1-10bp insertion) for the 11,960 TRAP sites in the transduced HEK293T-SpCas9 cells from Day 10, cultured in Dox-free medium.

Corresponding results for Dox-induction cells were shown in Fig.1e.

**Figure S12. Distribution of major indel types**

The proportion of indel types were classified to six major types: 1bp insertion, 2 bp insertion, long insertion (3-10 bp), small deletion (1-12 bp), medium deletion (13-21 bp) and large deletion (22-30 bp). Results were presented for the Dox-free groups. Others refers mainly WT reads, as well as a small proportion of reads containing indels larger than 30bp deletion or 10bp insertion.

**Figure S13. Analysis of SpCas9-introduced indels on protein translation for all 12,000 sites**

Indels (deletion/insertion) were analyzed based on they are of size 3n. For indels of size 3n, these indels were defined as in-frame indels. All the remain indels of 3n+1 and 3n+2 were defined as out-of-frame indels.

**Figure S14. The effect of gRNA spacer GC content on SpCas9 efficiency**

All the 12,000 gRNAs were separated into 9 groups based on the GC content (with an internal of 10%, e.g. 10-20% represents GC content >= 10% and <20%) of the 20nt spacer sequences. Box plots of the SpCas9 efficiency for all the corresponding gRNAs were presented for the indicated groups.

**Figure S15. The effect of gRNA spacer deltaG energy on SpCas9 efficiency**

All the 12,000 gRNAs were separated into 11 groups based on the deltaG energy (with an interval of 2; e.g. -15~-13 refers to deltaG energy >= -15 and < -13) of the 20nt spacer sequences. Box plots of the SpCas9 efficiency for all the corresponding gRNAs were presented for the indicated groups.

**Figure S16. The effect of GCC and TT motifs at N17-N20 of the gRNA Spacer on SpCas9 efficiency**

The approximately 12,000 gRNAs were divided into three groups base on whether there is a GCC or TT motif present at the seed N17-N20 region of the protospacer. The corresponding SpCas9 efficiency (indel percentage) of each gRNAs were presented with dot plot jittering. "***", p value less than 0.001. "****", p value less than 0.0001.

**Figure S17. Correlation between indels measured by TRAP-seq and indels predicted with inDelphi.**

For each gRNA, correlation was performed between the indels that we measured with TRAP-seq and the indels that we predicted using the machine learning program inDelphi. Results were presented with violin plot, with mean R value and 1 standard deviation given for each group. "n" refers to the number of gRNAs.

**Figure S18. Effect of N17 base on the indel outcome of 1bp insertion**

The gRNAs were divided into four groups based on the N17 base (A, T, C, G). Only gRNAs that contain 1bp insertion indel were included in this analysis. For each gRNA, the frequency of each inserted base (A, T, C, G) were presented for each group, with mean value indicated. Value for day 10 (Dox+) was shown in Fig. 2g.

**Figure S19. Effect of N18 base on the indel outcome of 1bp insertion**

The gRNAs were divided into four group based on the N18 base (A, T, C, G). Only gRNAs that contain 1bp insertion indel were included in this analysis. For each gRNA, the frequency of each inserted base (A, T, C, G) were presented for each group, with mean value indicated.

**Figure S20. Effect of N17N18 dinucleotide motifs on indel outcomes of 1bp insertion and deletions**

Effects of N17N18 dinucleotide motifs on the indel frequency of 1bp insertion and deletions (1-30 bp). The gRNAs were divided into 16 groups based on the N17N18 motifs. For each gRNA, the total indel frequencies of 1-30bp deletions and 1bp insertion were provided as dot plots. "n" indicates the number of gRNAs included for each group. Results for the Day10 (Dox+) group were presented in Fig. 2h.

**Figure S21. Spearman correlation analysis of the GNL machine learning tools for SpCas9, ABE, and CBE gRNA efficiency prediction**

A. Total number of gRNAs used of the GNL machine training (80%) and prediction (20%).

B. Spearman correlation-base analysis of the accuracy of GNL machine learning tools for SpCas9, ABE and CBE efficiency (random testing of 10 replicates).

**Figure S22. Schematic illustration of the ABE experiment**

The HEK293T-ABE cells stably express a doxycycline inducible ABE editor. MOI, multiplexity of infection. Days for passaging and harvesting the transduced cells for analysis were indicated. Detail description of the experimental procedure was shown in the methods. Transduced cells were harvested for analyses 2, 7 and 11 days after transduction.

**Figure S23. NGS coverage and representation of the 12K TRAP DNAs in the transduced HEK293T-ABE cells**

**a**. Violin plot of log1P (count +1) for each TRAP DNA for all ABE experimental groups

**b.** Pearson correlation analysis of TRAP representation between WT and HEK293T-ABE cells (with or without dox induction).

**Figure S24. Overall ABE efficiency in Dox-free transduce cells**

Histogram presentation of the overall ABE efficiency of all TRAP sites detected in the Dox-free HEK293T cells from 2, 7, and 11 days post transduction. Numbers indicate the number of gRNAs (TRAP) for each efficiency range (an interval of 5%). Results for the Dox-induction groups were shown in Fig. 3b.

**Figure S25. ABE editing profiles**

Summary of the ABE editing profiles for all adenine sites within the 37bp TRAP site. For each site, the average frequencies of A-to-G (A2G), A-to-T, A-to-C, and A-to-A (unedited) of all TRAP sites were presented in the percentile bar graph. Results were shown for the WT cells, and the Dox-free HEK293T-ABE cells from 7 and 11 days after transduction. Results for other groups were shown in Fig. 3c.

**Figure S26. Comparison of base frequency between low and high ABE efficiency TRAPs**

Two groups of TRAP sites were selected: Low efficiency (< 1%) and high efficiency (> 20%). Low efficiency refers to TRAP sites that the efficiency of any edited adenine within the N1-N20 is less than 1%. The high efficiency group refers to TRAP sites that at least one edited adenine within N1-

N20 has an efficiency higher than 20%. The base (A, T, C, G) frequency between the low and high efficiency TRAPs were presented. "n" indicates the number of TRAPs included.

**Figure S27. The effect of NAN trinucleotide motifs on ABE efficiency**

Scatter plot of edited A efficiency between sites within the active and repressive motifs. Active motifs include CAC, GAC, TAC, GAT, TAT, TAG, TAA. Repressive motifs include CAA, GAA, AAG, AAT, AAA. Bolded A refers to the deaminated adenine. The number of "n" indicates number of sites. "****", p value less than 0.0001. Results for the Day 11 (Dox+) HEK293T-ABE cells were shown in **Fig. 4d**.

**Figure S28. Effect of GC content on ABE efficiency.**

All the 12,000 gRNAs were separated into 9 groups based on the GC content (with an internal of 10%, 10-20% represents GC content >= 10% and <20%) of the 20nt spacer sequences. Box plots of the ABE efficiency (overall editing frequency) for all the corresponding gRNAs were presented for the indicated groups. Results for the Day 11 (Dox+) HEK293T-ABE cells were shown in **Fig. 4e**.

**Figure S29. The effect of gRNA spacer deltaG energy on ABE efficiency**

All the 12,000 gRNAs were separated into 11 groups based on the deltaG energy (with an interval of 2; e.g. -15~-13 refers to deltaG energy >= -15 and < -13) of the 20nt spacer sequences. Box plots of the ABE efficiency for all the corresponding gRNAs were presented for the indicated groups. Results for the Day 11 (Dox+) HEK293T-ABE cells were shown in **Fig. 4f**.

**Figure S30. The effect of GCC and TT motifs at N17-N20 on ABE efficiency**

The approximately 12,000 gRNAs were divided into three groups base on whether there is a GCC or TT motif present at the seed N17-N20 region. Scatter violin plots for the corresponding ABE efficiency of each gRNAs were presented. "**", p value less than 0.01. "****", p value less than 0.0001. Results for the Day 11 (Dox+) HEK293T-ABE cells were shown in **Fig. 4g**.

**Figure S31. Spearman evaluation of the GNL machine learning tool for prediction of ABE efficiency at each site of the editing window (N3-N11).**

A. Bar plot of the number of sites used for model training (80%) and evaluation (20%).

B. Spearman correlation-based evaluation of model accuracy for predicting the ABE base editing efficiency at each site within N3-N11 (values were shown as the mean and one SD, 10 random tests).

**Figure S32. Machine learning-based identification of features affecting ABE efficiency at each site (N3-N11)**

Efficiencies at each site within the ABE editing window (N3-N11) were used for machine learning trainings. Only the ABE efficiency from Day 11 (Dox+) HEK293T-ABE cells was use for this analysis. Top 20 features that weighted the most for the GNL machine learning model were provided for each site. Results were shown as the SHAP (SHapley Additive exPlanations) values. The 30mere comprises 4bp upstream, 20bp protospacer, 3 bp PAM, and 3 bp downstream sequences. Machine learning was based on gRNA efficiency data from Dox-induction cells at Day 10. For simplification, nucleotide features, such as “A”, “AA”, “GGC” refer to the total count of each motif with the TRAP region, “G_1” refers to “G” at N1 position. 5mer-start refers to the melting temperature of N1-N5. 5mer-end refers to the melting temperature of N16-N20. Tm global-spacer, refer to the melting temperature of the spacer N1-N20. 8mer-middel refers to the melting temperature of N7-N14.

**Figure S33. Schematic illustration of the CBE experiment**

The HEK293T-CBE cells stably express a doxycycline inducible CBE editor. MOI, multiplexity of infection. Days for passaging and harvesting the transduced cells for analysis were indicated. Detail description of the experimental procedure was shown in the methods. For CBE experiment, we did not analyze the intermediate time point.

**Figure S34. NGS coverage and representation of the 12K TRAP DNAs in the transduced HEK293T-CBE cells**

**a**. Violin plot of log1P (count +1) for each TRAP DNA for all CBE experimental groups

**b.** Pearson correlation analysis of TRAP representation between WT and HEK293T-CBE cells (with or without dox induction).

**Figure S35. Overall CBE efficiency in Dox-free transduced cells**

Histogram presentation of the overall CBE efficiency of all TRAP sites detected in the Dox-free HEK293T-CBE cells from 2, 7, and 11 days post transduction. Numbers indicate the number of gRNAs (TRAP) for each efficiency range (an interval of 5%). Results for the Dox-induction groups were shown in Fig. 5b.

**Figure 36 High throughput evaluation of recoding to stop codons by TRAP seq**

**b.** Summary of the median efficiency of stop-codon recoding by CBE of 11,875 TRAP sites, ranked by total efficiency.

**c.** Quantification of genes with the 12K TRAP-seq library, that were successfully recoded by CBE.

**Figure S37. CBE-mediated stop-codon recoding efficiency measured by TRAP-seq**

A representative of the six types of stop-codon recoding events by CBE, revealed by TRAP-seq. The frequency for each indel reads was presented. Indels that lead to the generation of stop codons were highlighted in orange text. TRAP (ID 61) was replotted in Figure 5e.

**Figure 38 CBE editing window and specificity**

**a.** Heatmap quantification of the overall C-to-T editing efficiency within the 20nt protospacer region.

**b.** Summary of substituted cytosines and specificity within the CBE editing window N1-N16.

**Figure S39. Comparison of base frequency between low and high CBE efficiency TRAPs**

Two groups of TRAP sites were selected: Low efficiency (< 5%) and high efficiency (> 50%). The low efficiency group refers to TRAP sites that the efficiency of any edited cytosine within the N1-N20 is less than 5%. The high efficiency group refers to TRAP sites at least one edited cytosine within N1-N20 has an efficiency higher than 50%. The base (A, T, C, G) frequency between the low and high efficiency TRAPs were presented. "n" indicates the number of TRAPs included.

**Figure S40. The effect of NAN trinucleotide motifs on ABE efficiency**

Dot plots of edited C efficiency between sites within the active and repressive motifs. Active motifs include C**C**A, T**C**C, C**C**C, T**C**A, A**C**C, T**C**T, T**C**G. Repressive motifs include A**C**A, G**C**C, G**C**A, G**C**T, G**C**G. Bolded C refers to the deaminated cytosine. The number of "n" indicates number of sites.

"****", p value less than 0.0001. Results for the Day 11 (Dox+) HEK293T-CBE cells were shown in **Fig. 6d**.

**Figure S41. Effect of GC content on CBE efficiency.**

All the 12,000 gRNAs were separated into 9 groups based on the GC content (with an internal of 10%, 10-20% represents GC content >= 10% and <20%) of the 20nt spacer sequences. The CBE efficiency (overall editing frequency) for all the corresponding gRNAs were presented for the indicated groups. Results for the Day 11 (Dox+) HEK293T-CBE cells were shown in **Fig. 6e**.

**Figure S42. The effect of gRNA spacer deltaG energy on CBE efficiency**

All the 12,000 gRNAs were separated into 11 groups based on the deltaG energy (with an interval of 2; e.g. -15~-13 refers to deltaG energy >= -15 and < -13) of the 20nt spacer sequences. The CBE efficiency for all the corresponding gRNAs were presented for the indicated groups. Results for the Day 11 (Dox+) HEK293T-CBE cells were shown in **Fig. 6f**.

**Figure S43. The effect of GCC and TT motifs at N17-N20 of the gRNA Spacer on CBE efficiency**

The approximately 12,000 gRNAs were divided into three groups base on whether there is a GCC or TT motif at the seed N17-N20 region. Dot plots for the corresponding CBE efficiency of each gRNAs were presented. "*", p value less than 0.05. "****", p value less than 0.0001. Results for the Day 11 (Dox+) HEK293T-CBE cells were shown in **Fig. 6g**.

**Figure S44. Spearman evaluation of the GNL machine learning tool for prediction of CBE efficiency at each site of gRNA spacer (N1-N20).**

A. Bar plot of the number of sites used for model training (80%) and evaluation (20%).

B. Spearman correlation-based evaluation of model accuracy for predicting the CBE base editing efficiency at each site within N1-N20 (values were shown as the mean and one SD, 10 random tests).

**Figure S45. Machine learning-based identification of features affecting CBE efficiency at each site (N1-N20)**

Efficiencies at each site within the spacer (N1-N20) were used for machine learning trainings. Only the CBE efficiency from Day 11 (Dox+) HEK293T-CBE cells was use for this analysis. Top 20 features that weighted the most for the GNL machine learning model were provided for each site.

Results were shown as the SHAP (SHapley Additive exPlanations) values. The 30mere comprises 4bp upstream, 20bp protospacer, 3 bp PAM, and 3 bp downstream sequences. Machine learning was based on gRNA efficiency data from Dox-induction cells at Day 10. For simplification, nucleotide features, such as “A”, “AA”, “GGC” refer to the total count of each motif with the TRAP region, “G_1” refers to “G” at N1 position. 5mer-start refers to the melting temperature of N1-N5. 5mer-end refers to the melting temperature of N16-N20. Tm global-spacer, refer to the melting temperature of the spacer N1-N20. 8mer-middel refers to the melting temperature of N7-N14.

S1

Illustration of the TRAP-seq vector and cloning

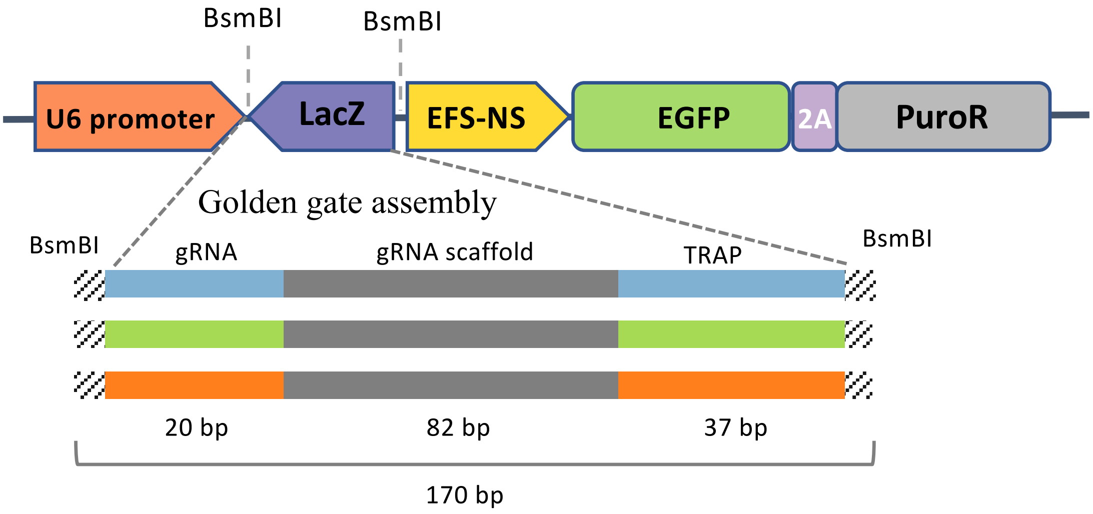

S2

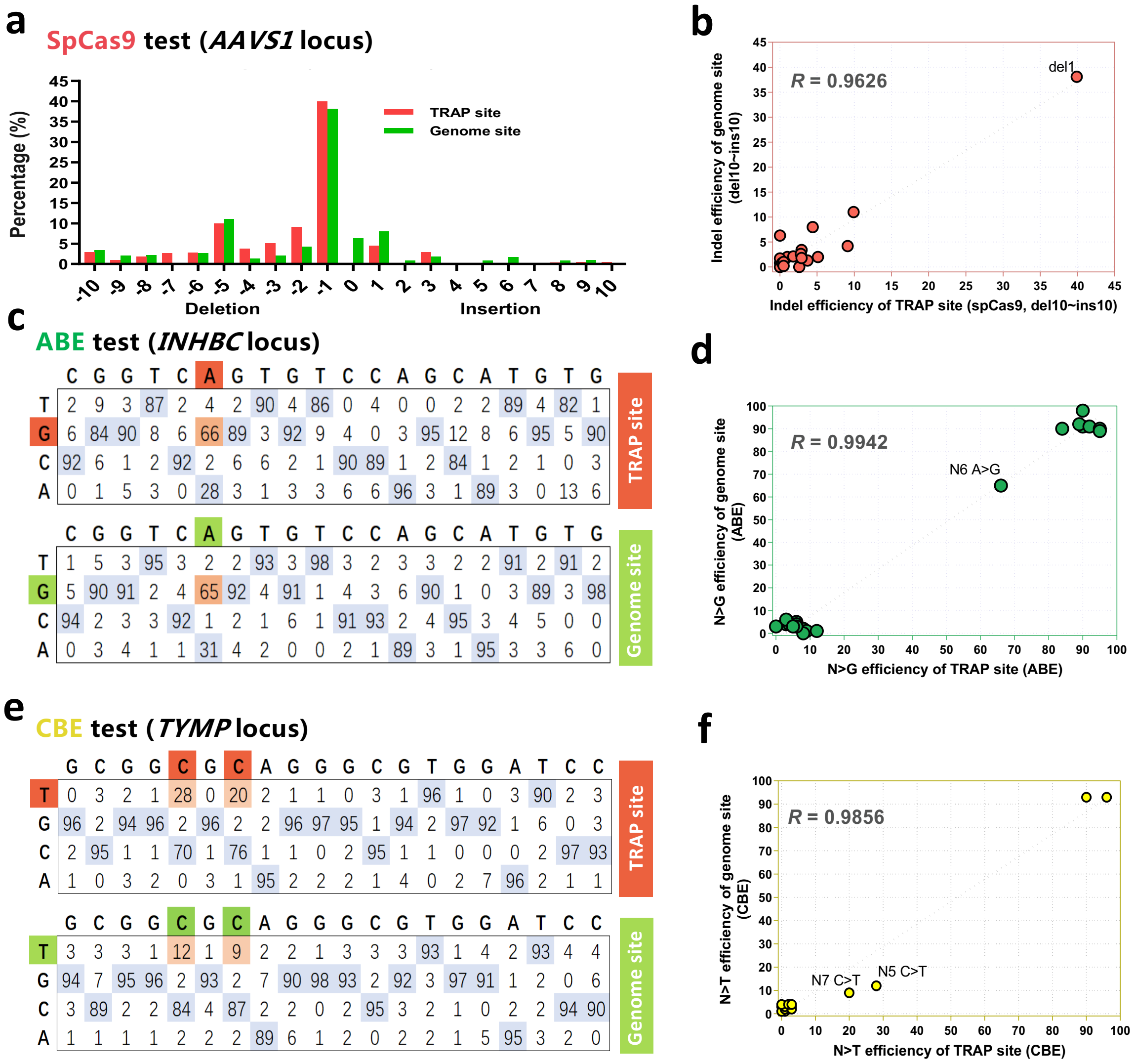
a
SpCas9 test (AAVS1 locus)
Percentage (%)
45 40 35 30 25 20 15 10 5 0
TRAP site
Genome site
-10 -9 -8 -7 -6 -5 -4 -3 -2 -1 0 1 2 3 4 5 6 7 8 9 10
Deletion
Insertion
b
R = 0.9626
del1
Indel efficiency of genome site (del10~ins10)
Indel efficiency of TRAP site (spCas9, del10~ins10)
c
ABE test (INHBC locus)
C G G T C A G T G T C C A G C A T G T G
T 2 9 3 87 2 4 2 90 4 86 0 4 0 0 2 2 89 4 82 1
G 6 84 90 8 6 66 89 3 92 9 4 0 3 95 12 8 6 95 5 90
C 92 6 1 2 92 2 6 6 2 1 90 89 1 2 84 1 2 1 0 3
A 0 1 5 3 0 28 3 1 3 3 6 6 96 3 1 89 3 0 13 6
TRAP site
C G G T C A G T G T C C A G C A T G T G
T 1 5 3 95 3 2 2 93 3 98 3 2 3 3 2 2 91 2 91 2
G 5 90 91 2 4 65 92 4 91 1 4 3 6 90 1 0 3 89 3 98
C 94 2 3 3 92 1 2 1 6 1 91 93 2 4 95 3 4 5 0 0
A 0 3 4 1 1 31 4 2 0 0 2 1 89 3 1 95 3 3 6 0
Genome site
d
R = 0.9942
N6 A>G
N>G efficiency of genome site (ABE)
N>G efficiency of TRAP site (ABE)
e
CBE test (TYMP locus)
G C G G C G C A G G G C G T G G A T C C
T 0 3 2 1 28 0 20 2 1 1 0 3 1 96 1 0 3 90 2 3
G 96 2 94 96 2 96 2 2 96 97 95 1 94 2 97 92 1 6 0 3
C 2 95 1 1 70 1 76 1 1 0 2 95 1 1 0 0 0 2 97 93
A 1 0 3 2 0 3 1 95 2 2 2 1 4 0 2 7 96 2 1 1
TRAP site
G C G G C G C A G G G C G T G G A T C C
T 3 3 3 1 12 1 9 2 2 1 3 3 3 93 1 4 2 93 4 4
G 94 7 95 96 2 93 2 7 90 98 93 2 92 3 97 91 1 2 0 6
C 3 89 2 2 84 4 87 2 2 0 2 95 3 2 1 0 2 2 94 90
A 1 1 1 1 2 2 2 89 6 1 2 0 2 2 1 5 95 3 2 0
Genome site
f
R = 0.9856
N7 C>T
N5 C>T
N>T efficiency of genome site (CBE)
N>T efficiency of TRAP site (CBE)

G

S3

### Optimization of protocols for amplifying the TRAP-12K oligo library by PCRs

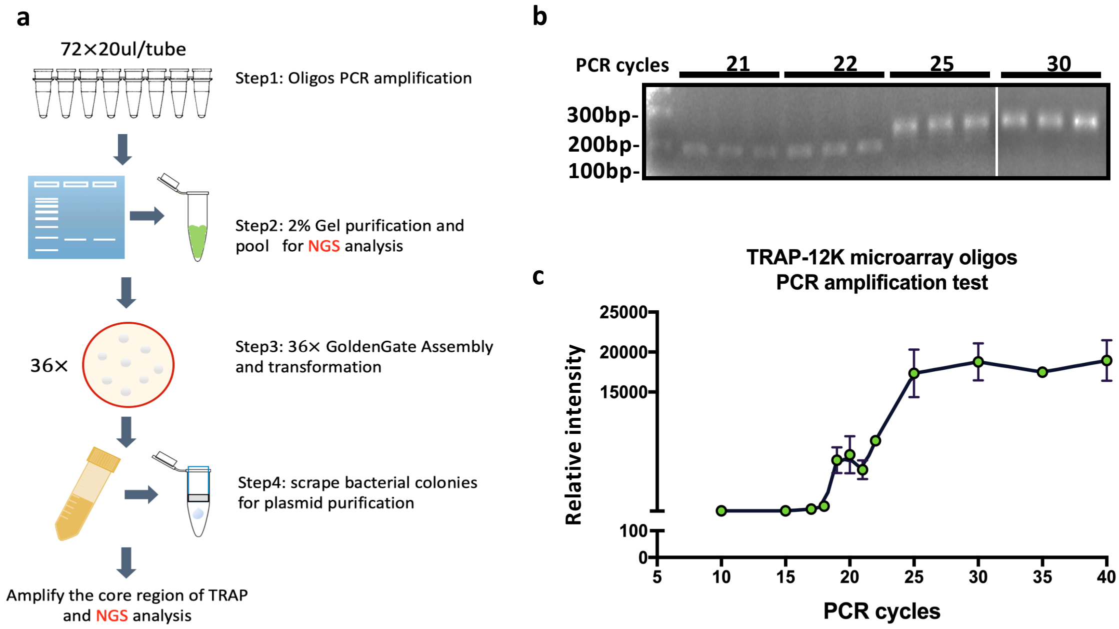

S4

Highly efficent cloning of the TRAP-12K oligos into the lenti-TRAP-seq vector by golden gate assembly

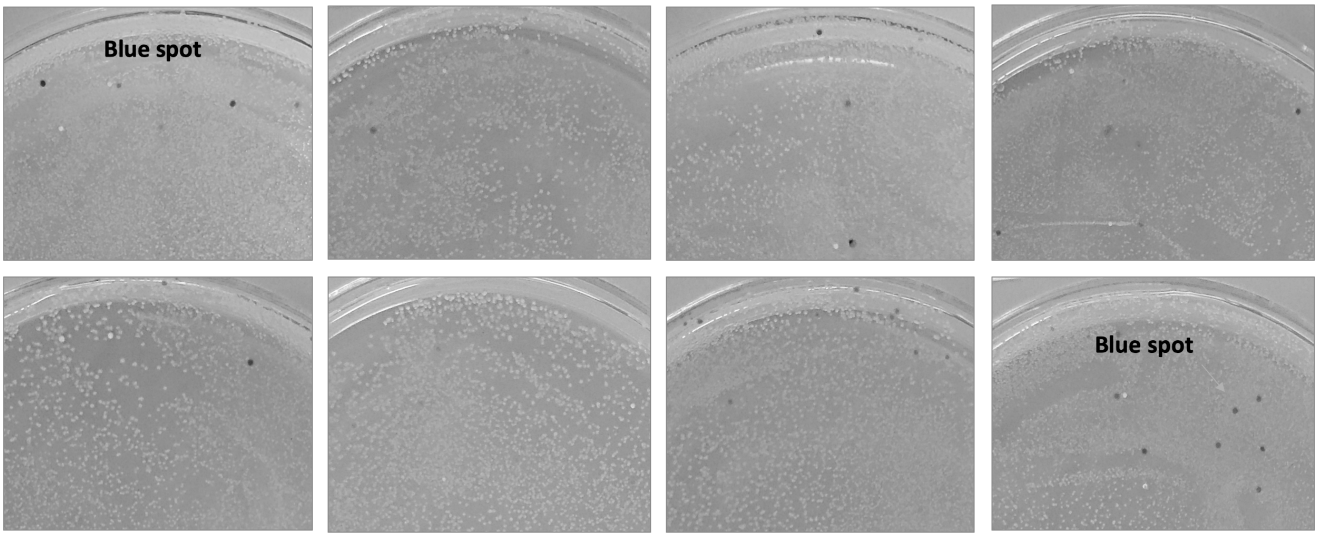

### Quantification of infection titer of the TRAP-12K lentivirus library

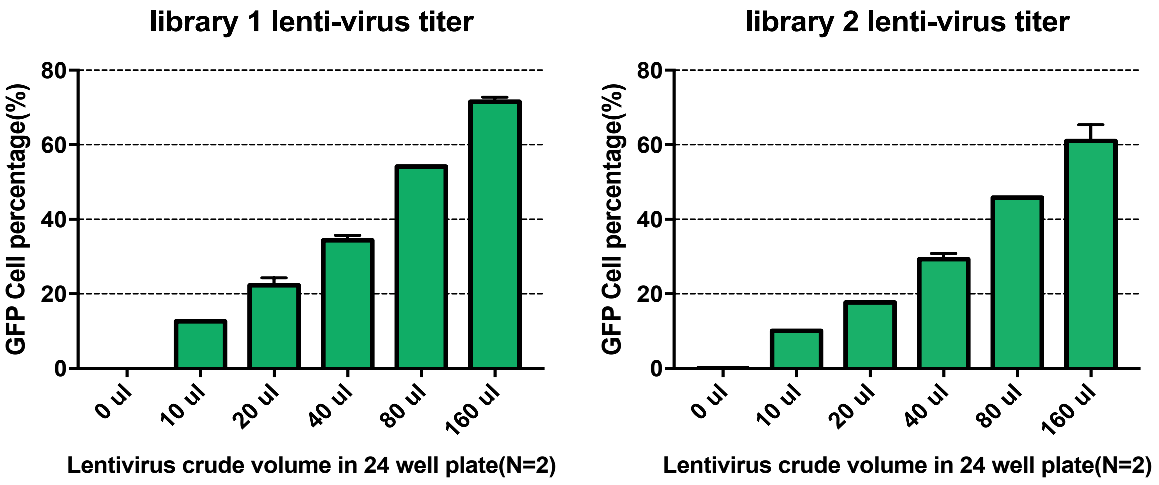

**TRAP-library prep 1titer= 3.8 x $10^6$ UI/mL**

**TRAP-library prep 2 titer= 2.5 x $10^6$ UI/mL**

QC of retaining the coverage of each TRAP in the TRAP-12K oligos,
GGC plasmids and transduce cells

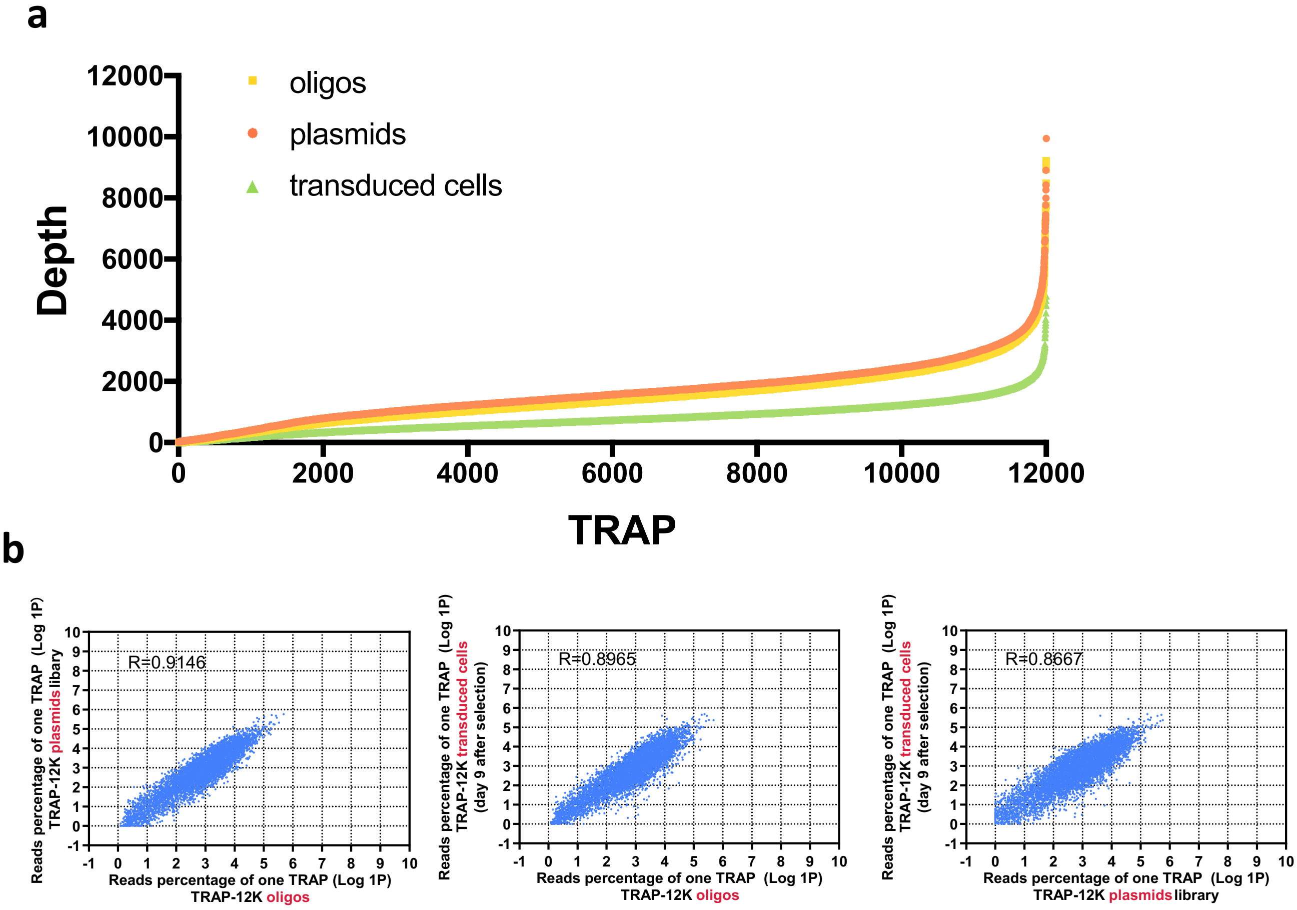

#### Schematic illustration of the experiment

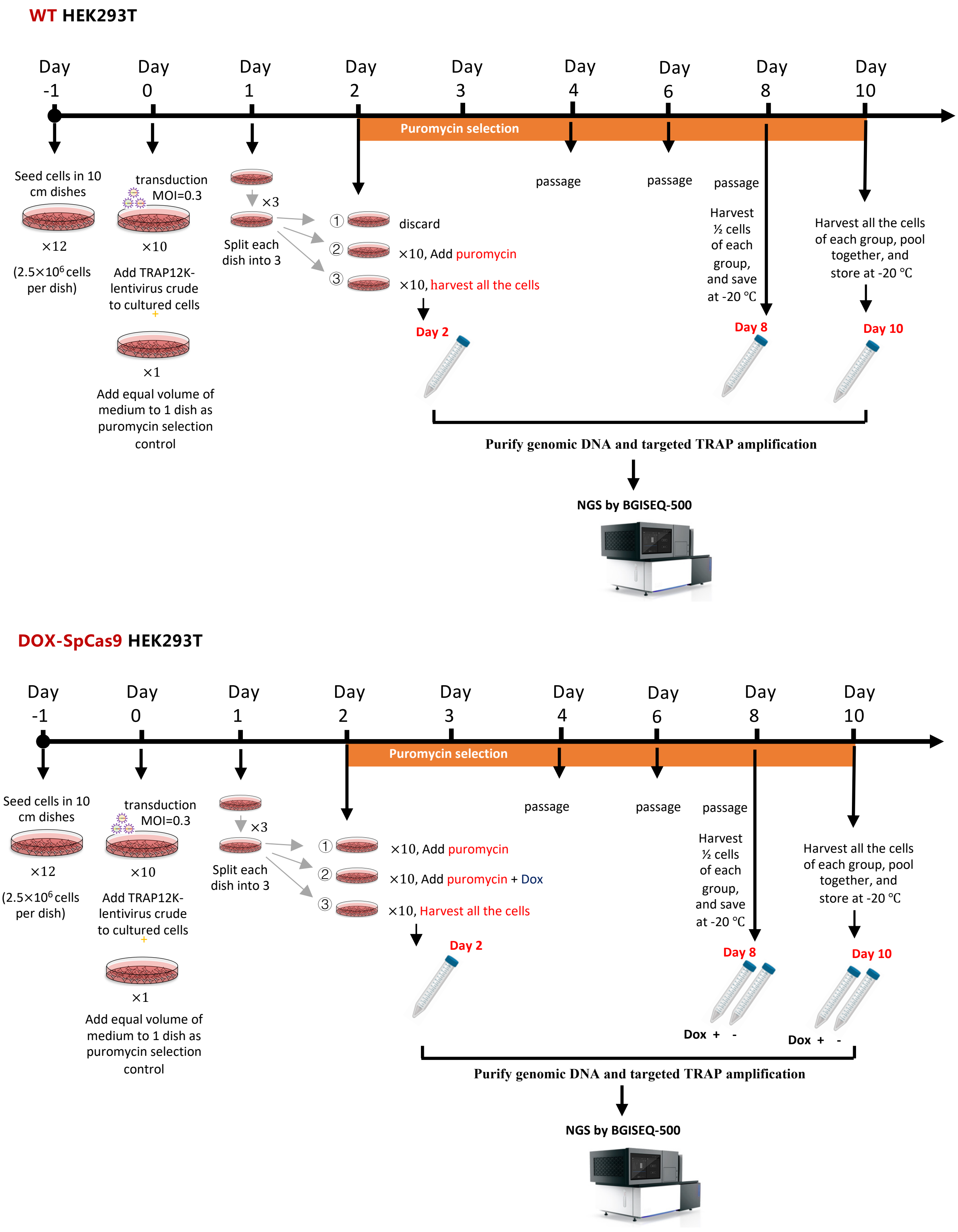

S8

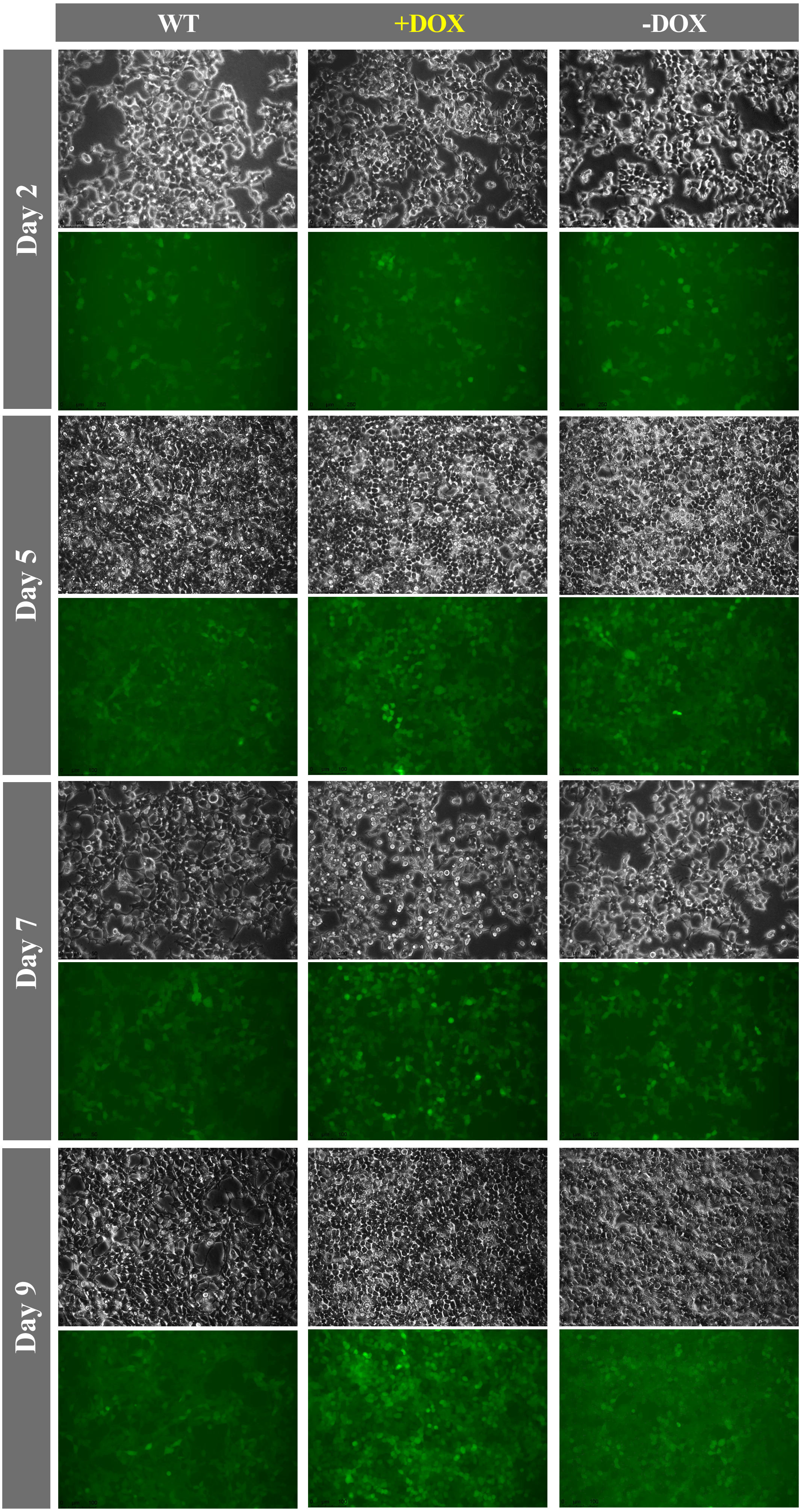

S9

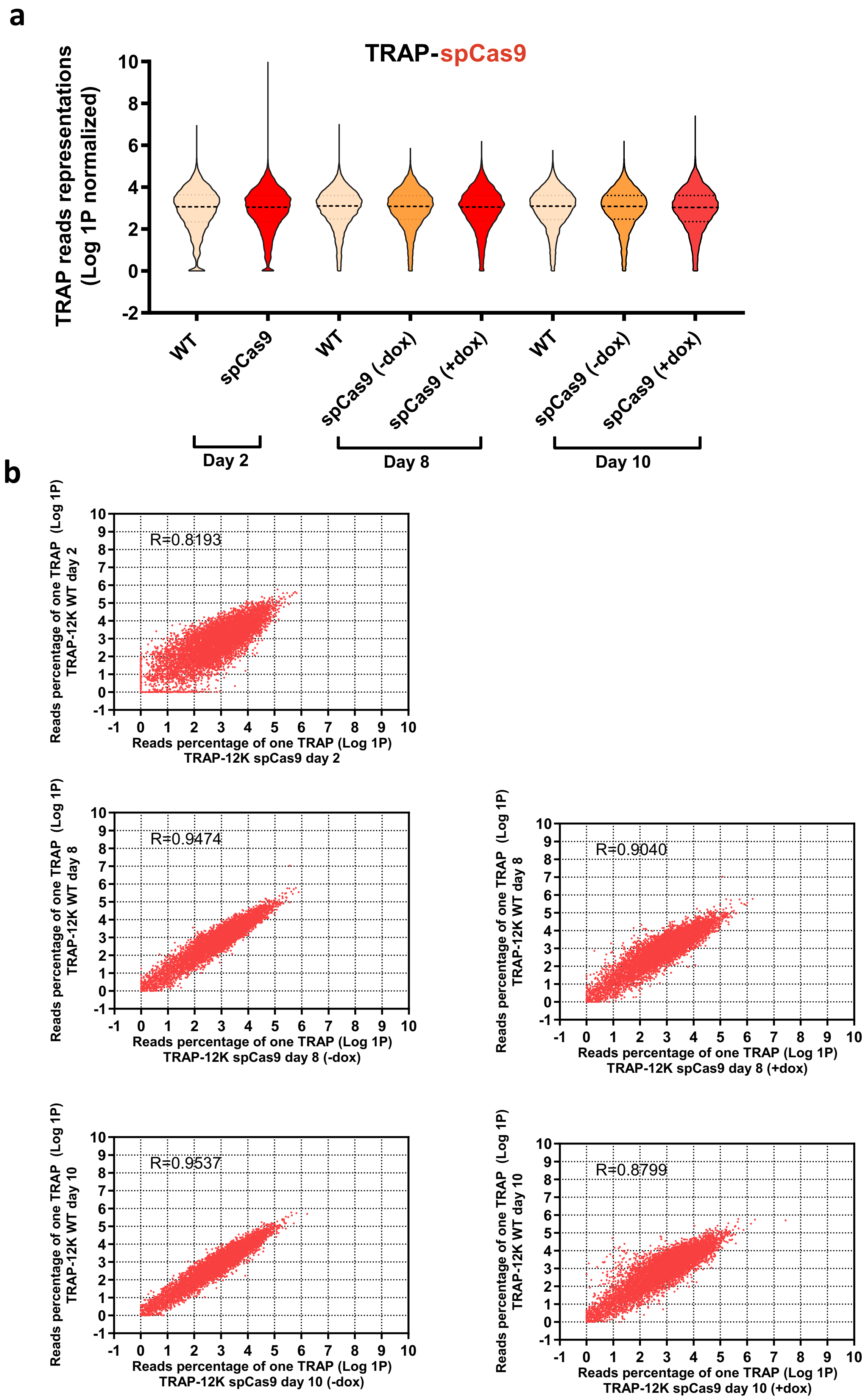
a
TRAP-spCas9
TRAP reads representations (Log 1P normalized)
WT
spCas9
WT
spCas9 (-dox)
spCas9 (+dox)
WT
spCas9 (-dox)
spCas9 (+dox)
Day 2
Day 8
Day 10
b
R=0.8193
Reads percentage of one TRAP (Log 1P) TRAP-12K WT day 2
Reads percentage of one TRAP (Log 1P) TRAP-12K spCas9 day 2
R=0.9474
Reads percentage of one TRAP (Log 1P) TRAP-12K WT day 8
Reads percentage of one TRAP (Log 1P) TRAP-12K spCas9 day 8 (-dox)
R=0.9040
Reads percentage of one TRAP (Log 1P) TRAP-12K WT day 8
Reads percentage of one TRAP (Log 1P) TRAP-12K spCas9 day 8 (+dox)
R=0.9537
Reads percentage of one TRAP (Log 1P) TRAP-12K WT day 10
Reads percentage of one TRAP (Log 1P) TRAP-12K spCas9 day 10 (-dox)
R=0.8799
Reads percentage of one TRAP (Log 1P) TRAP-12K WT day 10
Reads percentage of one TRAP (Log 1P) TRAP-12K spCas9 day 10 (+dox)

**S10**

**a**

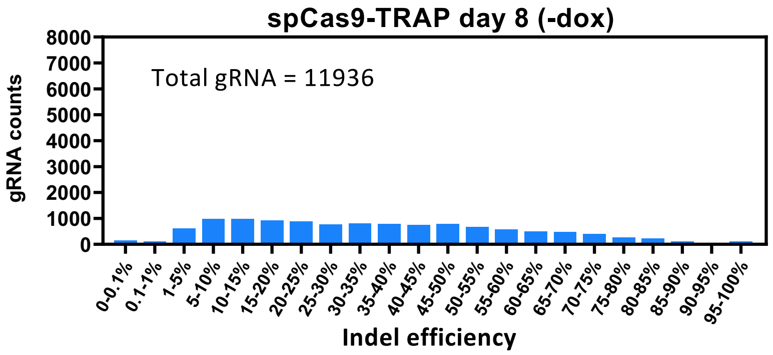
spCas9-TRAP day 8 (-dox)
Total gRNA = 11936
gRNA counts
8000
7000
6000
5000
4000
3000
2000
1000
0
0-0.1%
0.1-1%
1-5%
5-10%
10-15%
15-20%
20-25%
25-30%
30-35%
35-40%
40-45%
45-50%
50-55%
55-60%
60-65%
65-70%
70-75%
75-80%
80-85%
85-90%
90-95%
95-100%
Indel efficiency

**b**

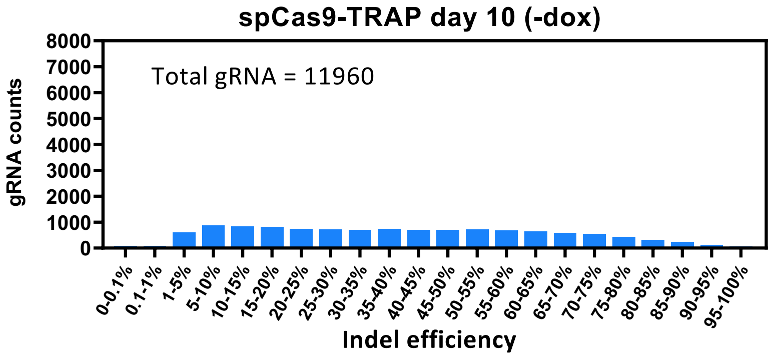
spCas9-TRAP day 10 (-dox)
Total gRNA = 11960
gRNA counts
8000
7000
6000
5000
4000
3000
2000
1000
0
0-0.1%
0.1-1%
1-5%
5-10%
10-15%
15-20%
20-25%
25-30%
30-35%
35-40%
40-45%
45-50%
50-55%
55-60%
60-65%
65-70%
70-75%
75-80%
80-85%
85-90%
90-95%
95-100%
Indel efficiency

**S11**

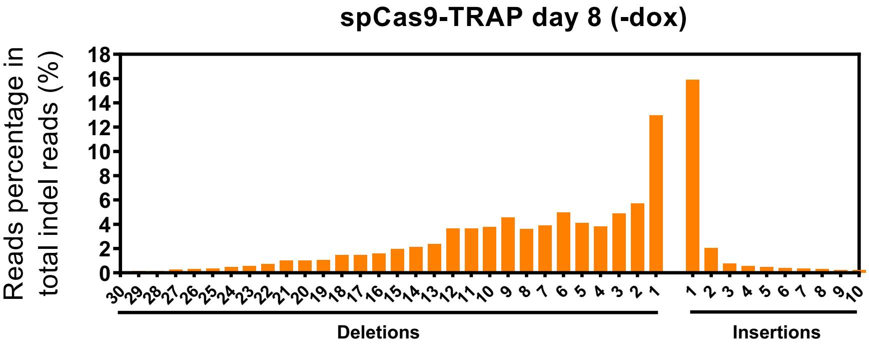
spCas9-TRAP day 8 (-dox)
Reads percentage in total indel reads (%)
0
2
4
6
8
10
12
14
16
18
30 29 28 27 26 25 24 23 22 21 20 19 18 17 16 15 14 13 12 11 10 9 8 7 6 5 4 3 2 1
1 2 3 4 5 6 7 8 9 10
Deletions
Insertions

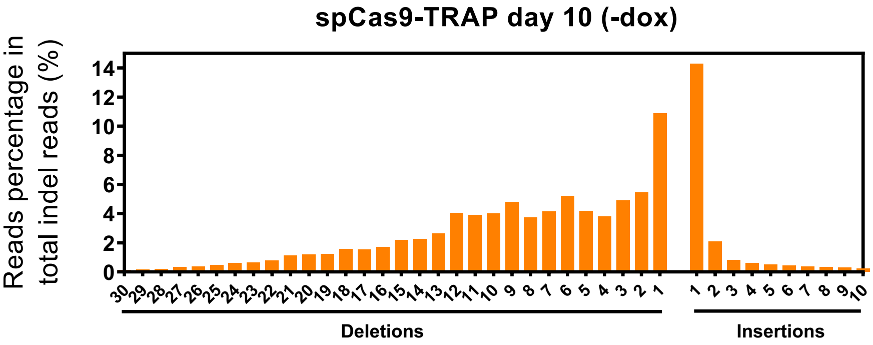
spCas9-TRAP day 10 (-dox)
Reads percentage in total indel reads (%)
0
2
4
6
8
10
12
14
30 29 28 27 26 25 24 23 22 21 20 19 18 17 16 15 14 13 12 11 10 9 8 7 6 5 4 3 2 1
1 2 3 4 5 6 7 8 9 10
Deletions
Insertions

**S12**

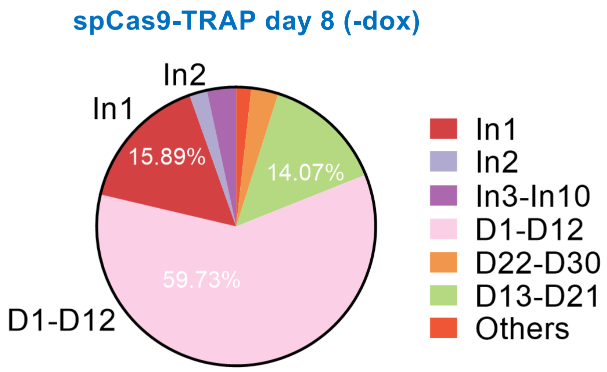
spCas9-TRAP day 8 (-dox)
In2
In1
15.89%
14.07%
59.73%
D1-D12
In1
In2
In3-In10
D1-D12
D22-D30
D13-D21
Others

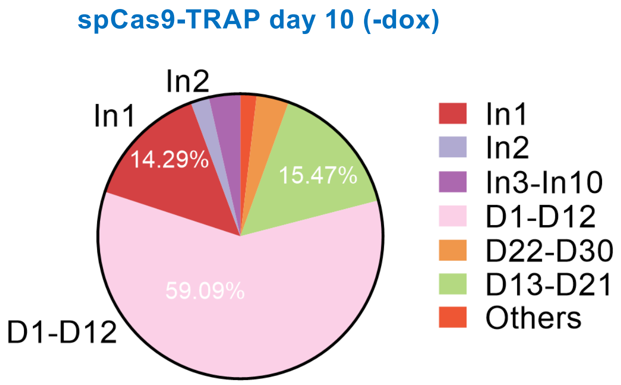
spCas9-TRAP day 10 (-dox)
In2
In1
14.29%
15.47%
59.09%
D1-D12
In1
In2
In3-In10
D1-D12
D22-D30
D13-D21
Others

**S13**

### Proportion of in-frame and out of frame indels

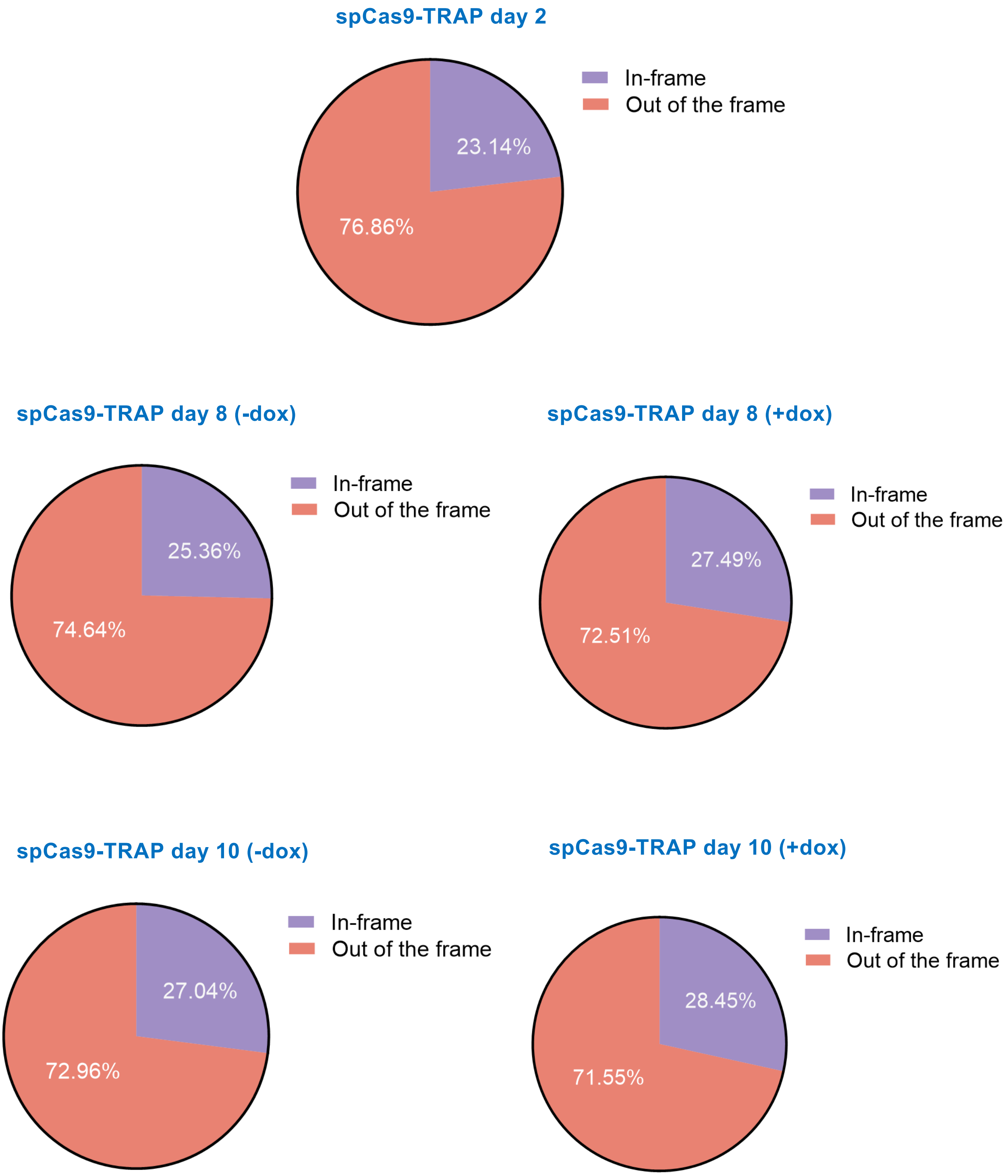

**S14**

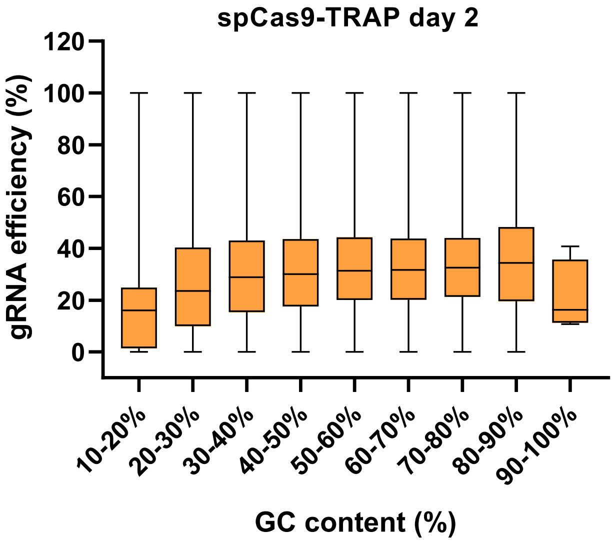
spCas9-TRAP day 2
gRNA efficiency (%)
120
100
80
60
40
20
0
10-20%
20-30%
30-40%
40-50%
50-60%
60-70%
70-80%
80-90%
90-100%
GC content (%)

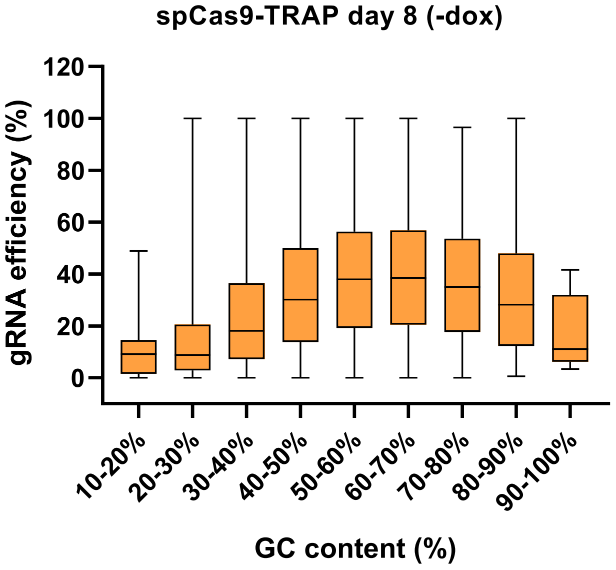
spCas9-TRAP day 8 (-dox)
gRNA efficiency (%)
120
100
80
60
40
20
0
10-20%
20-30%
30-40%
40-50%
50-60%
60-70%
70-80%
80-90%
90-100%
GC content (%)

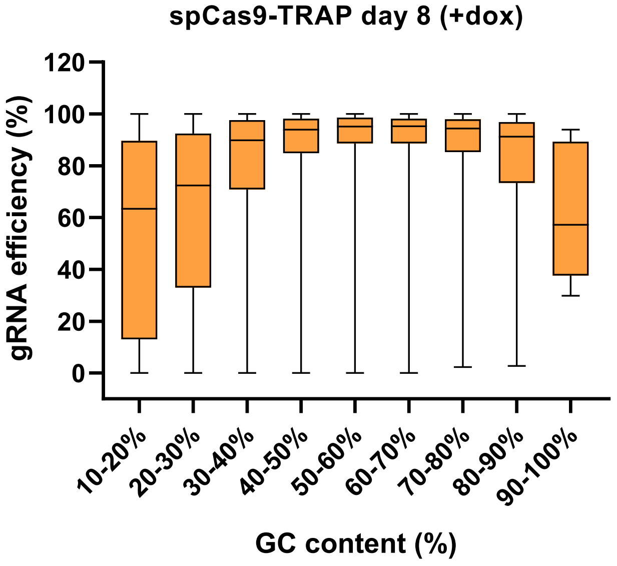
spCas9-TRAP day 8 (+dox)
gRNA efficiency (%)
120
100
80
60
40
20
0
10-20%
20-30%
30-40%
40-50%
50-60%
60-70%
70-80%
80-90%
90-100%
GC content (%)

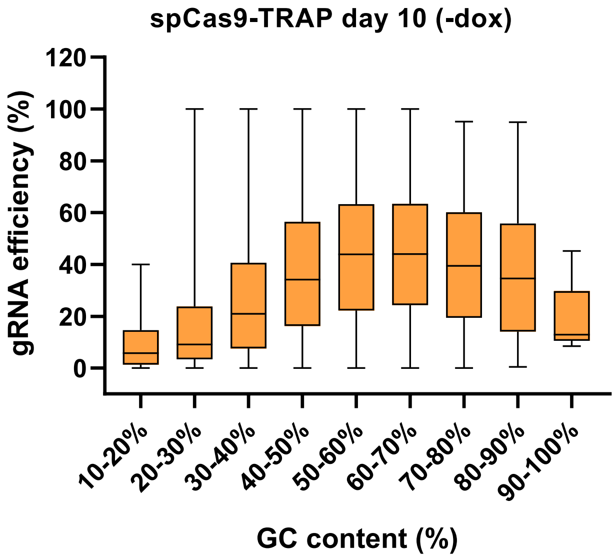
spCas9-TRAP day 10 (-dox)
gRNA efficiency (%)
120
100
80
60
40
20
0
10-20%
20-30%
30-40%
40-50%
50-60%
60-70%
70-80%
80-90%
90-100%
GC content (%)

**S15**

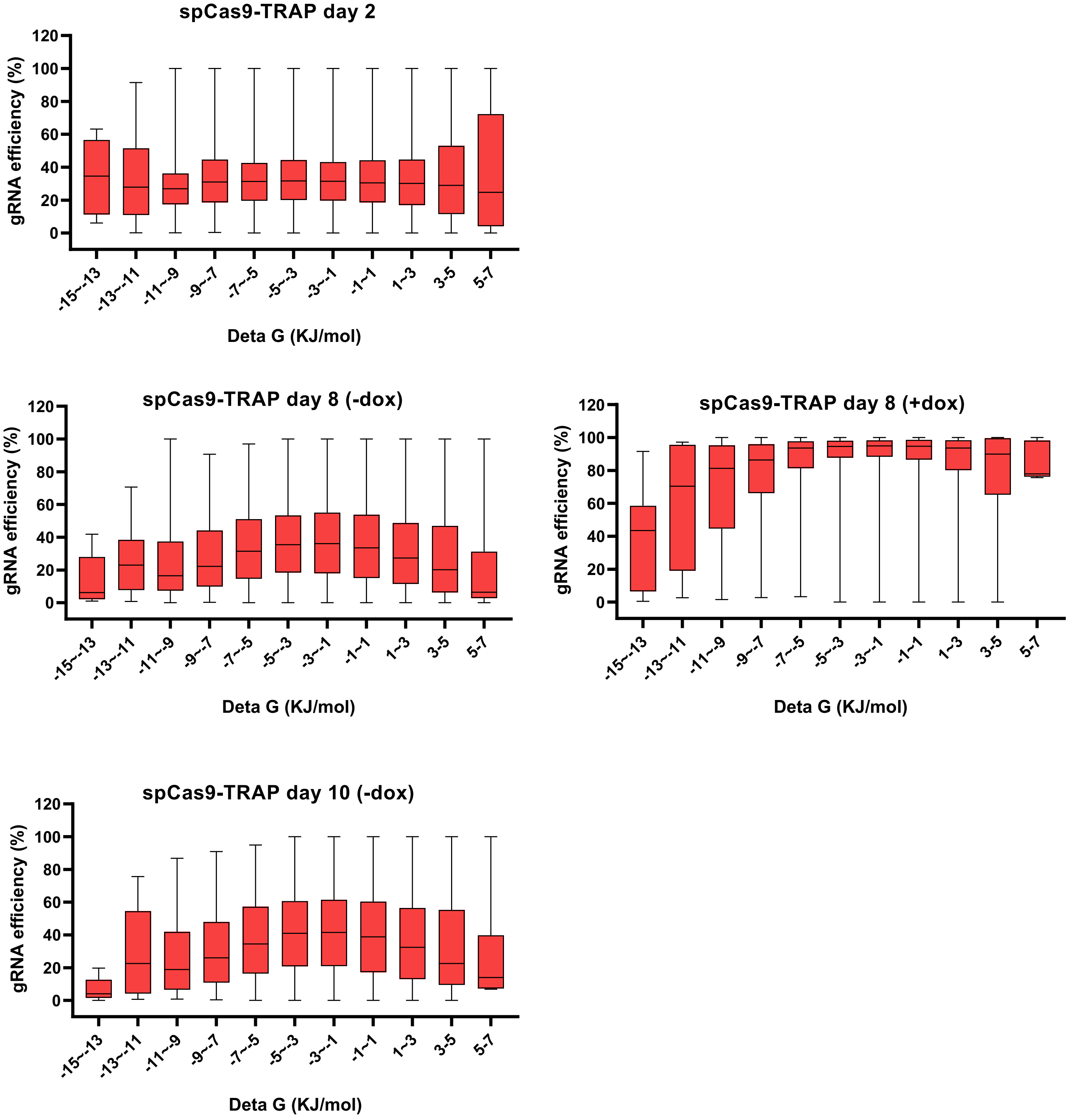
spCas9-TRAP day 2
gRNA efficiency (%)
0
20
40
60
80
100
120
-15~-13
-13~-11
-11~-9
-9~-7
-7~-5
-5~-3
-3~-1
-1~1
1~3
3-5
5-7
Deta G (KJ/mol)
spCas9-TRAP day 8 (-dox)
gRNA efficiency (%)
Deta G (KJ/mol)
spCas9-TRAP day 8 (+dox)
gRNA efficiency (%)
Deta G (KJ/mol)
spCas9-TRAP day 10 (-dox)
gRNA efficiency (%)
Deta G (KJ/mol)

**S16**

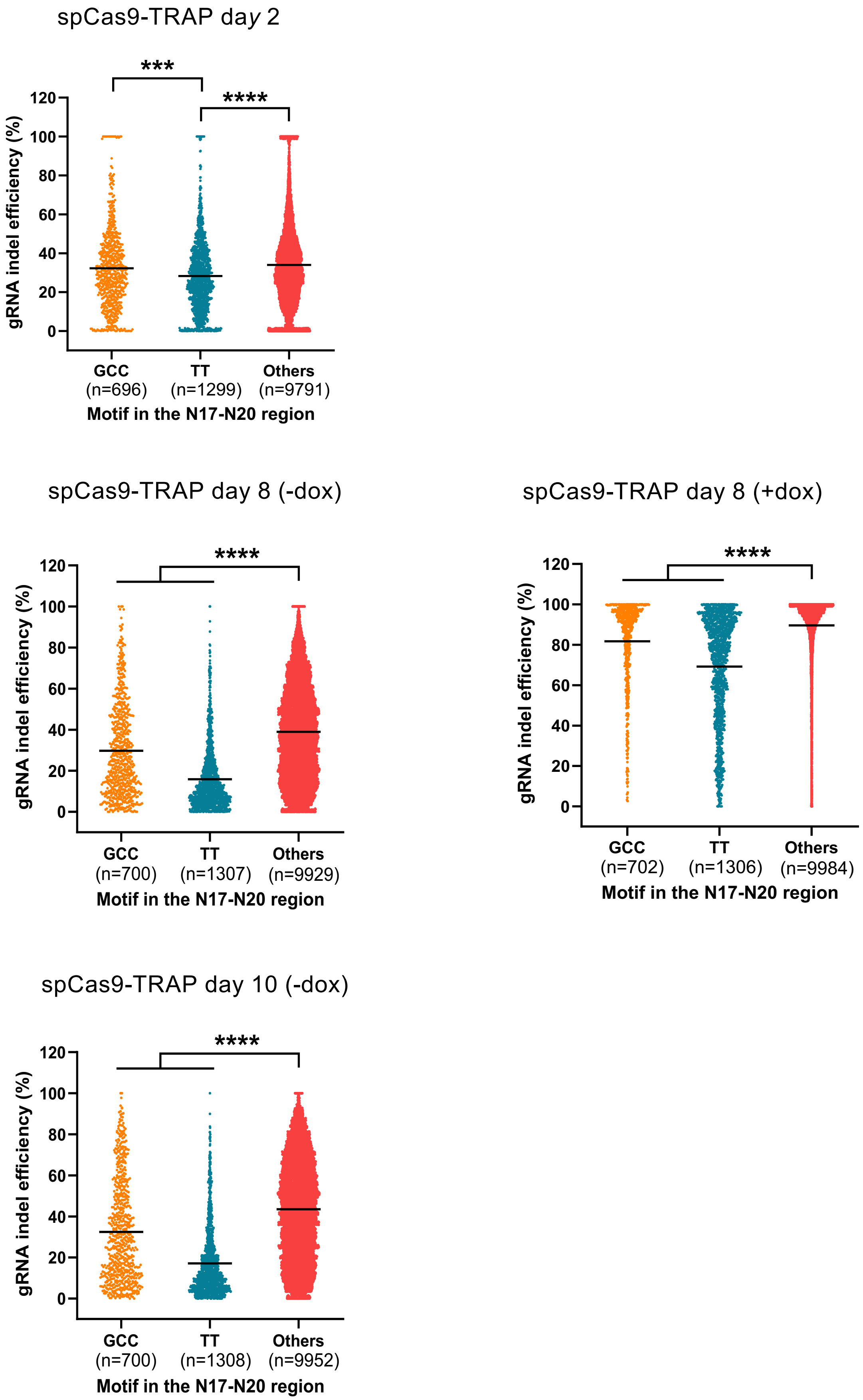
spCas9-TRAP day 2
***
****
gRNA indel efficiency (%)
120
100
80
60
40
20
0
GCC
(n=696)
TT
(n=1299)
Others
(n=9791)
Motif in the N17-N20 region
spCas9-TRAP day 8 (-dox)
****
gRNA indel efficiency (%)
120
100
80
60
40
20
0
GCC
(n=700)
TT
(n=1307)
Others
(n=9929)
Motif in the N17-N20 region
spCas9-TRAP day 8 (+dox)
****
gRNA indel efficiency (%)
120
100
80
60
40
20
0
GCC
(n=702)
TT
(n=1306)
Others
(n=9984)
Motif in the N17-N20 region
spCas9-TRAP day 10 (-dox)
****
gRNA indel efficiency (%)
120
100
80
60
40
20
0
GCC
(n=700)
TT
(n=1308)
Others
(n=9952)
Motif in the N17-N20 region

**S17**

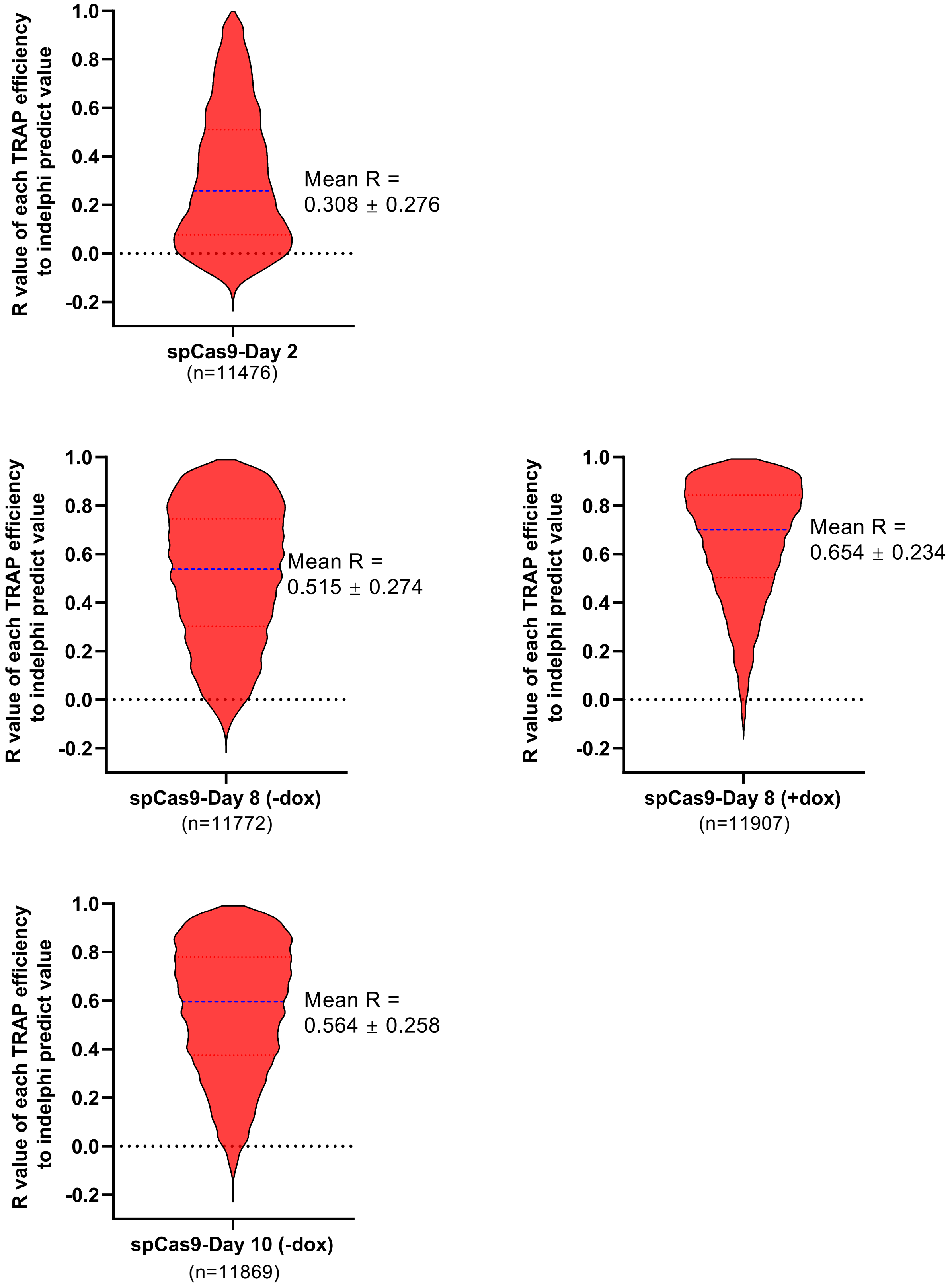
R value of each TRAP efficiency to indelphi predict value
1.0
0.8
0.6
0.4
0.2
0.0
-0.2
Mean R =
0.308 ± 0.276
spCas9-Day 2
(n=11476)
R value of each TRAP efficiency to indelphi predict value
Mean R =
0.515 ± 0.274
spCas9-Day 8 (-dox)
(n=11772)
R value of each TRAP efficiency to indelphi predict value
Mean R =
0.654 ± 0.234
spCas9-Day 8 (+dox)
(n=11907)
R value of each TRAP efficiency to indelphi predict value
Mean R =
0.564 ± 0.258
spCas9-Day 10 (-dox)
(n=11869)

# S18

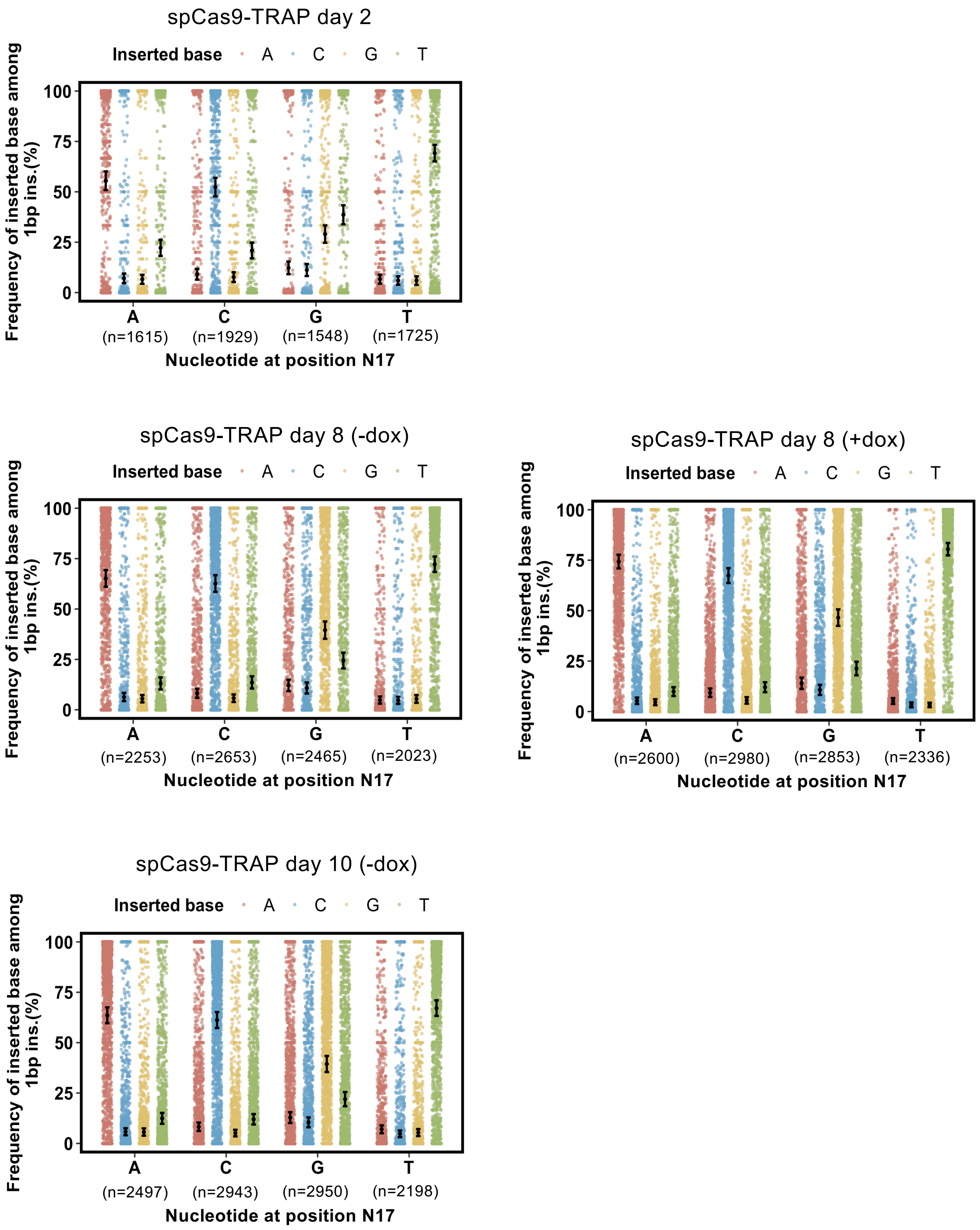
spCas9-TRAP day 2
Inserted base A C G T
Frequency of inserted base among 1bp ins.(%)
100
75
50
25
0
A (n=1615)
C (n=1929)
G (n=1548)
T (n=1725)
Nucleotide at position N17
spCas9-TRAP day 8 (-dox)
Inserted base A C G T
Frequency of inserted base among 1bp ins.(%)
A (n=2253)
C (n=2653)
G (n=2465)
T (n=2023)
Nucleotide at position N17
spCas9-TRAP day 8 (+dox)
Inserted base A C G T
Frequency of inserted base among 1bp ins.(%)
A (n=2600)
C (n=2980)
G (n=2853)
T (n=2336)
Nucleotide at position N17
spCas9-TRAP day 10 (-dox)
Inserted base A C G T
Frequency of inserted base among 1bp ins.(%)
A (n=2497)
C (n=2943)
G (n=2950)
T (n=2198)
Nucleotide at position N17

**S19**

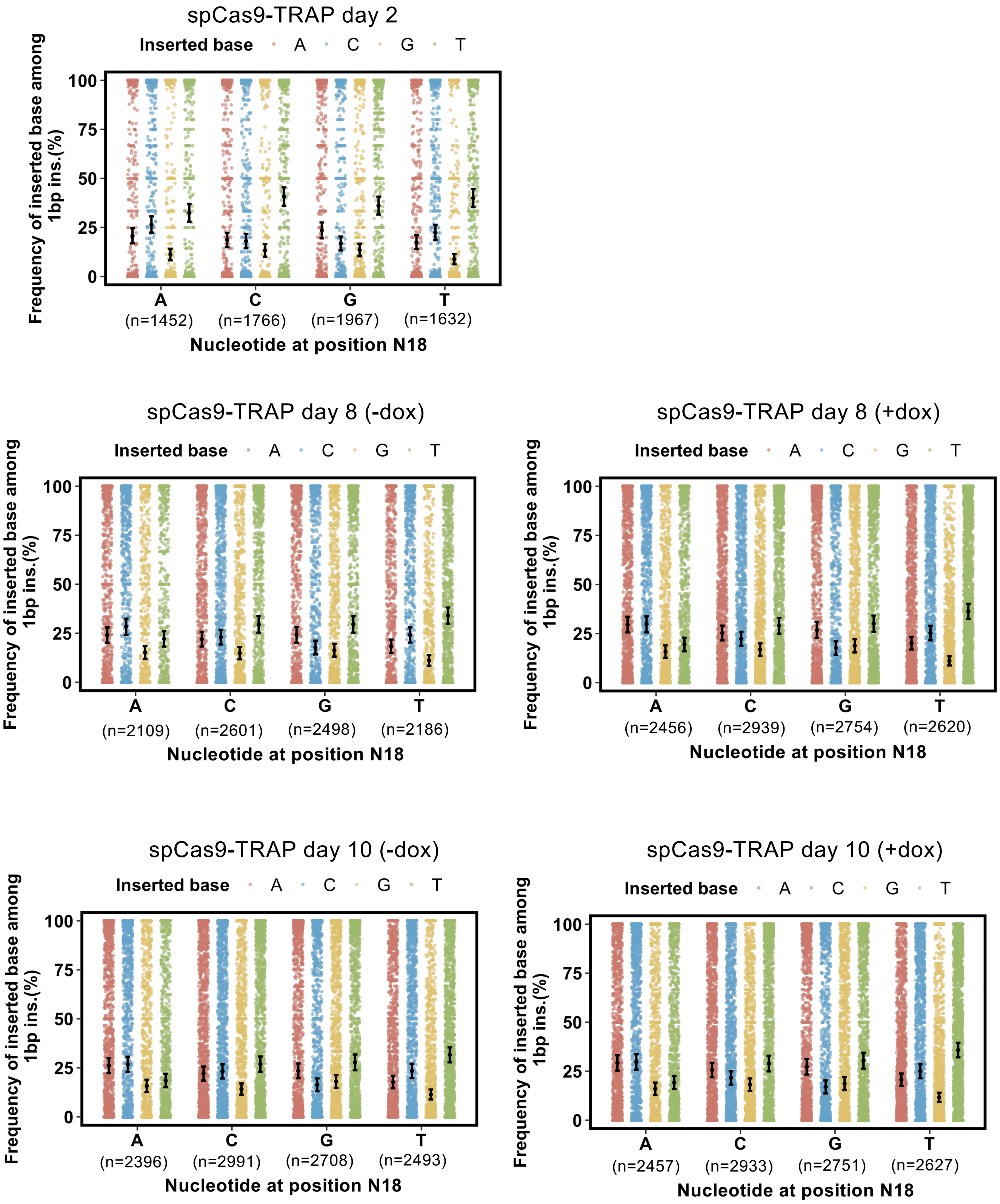
spCas9-TRAP day 2
Inserted base A C G T
Frequency of inserted base among 1bp ins.(%)
100
75
50
25
0
A (n=1452)
C (n=1766)
G (n=1967)
T (n=1632)
Nucleotide at position N18
spCas9-TRAP day 8 (-dox)
Inserted base A C G T
Frequency of inserted base among 1bp ins.(%)
A (n=2109)
C (n=2601)
G (n=2498)
T (n=2186)
Nucleotide at position N18
spCas9-TRAP day 8 (+dox)
Inserted base A C G T
Frequency of inserted base among 1bp ins.(%)
A (n=2456)
C (n=2939)
G (n=2754)
T (n=2620)
Nucleotide at position N18
spCas9-TRAP day 10 (-dox)
Inserted base A C G T
Frequency of inserted base among 1bp ins.(%)
A (n=2396)
C (n=2991)
G (n=2708)
T (n=2493)
Nucleotide at position N18
spCas9-TRAP day 10 (+dox)
Inserted base A C G T
Frequency of inserted base among 1bp ins.(%)
A (n=2457)
C (n=2933)
G (n=2751)
T (n=2627)
Nucleotide at position N18

# S20

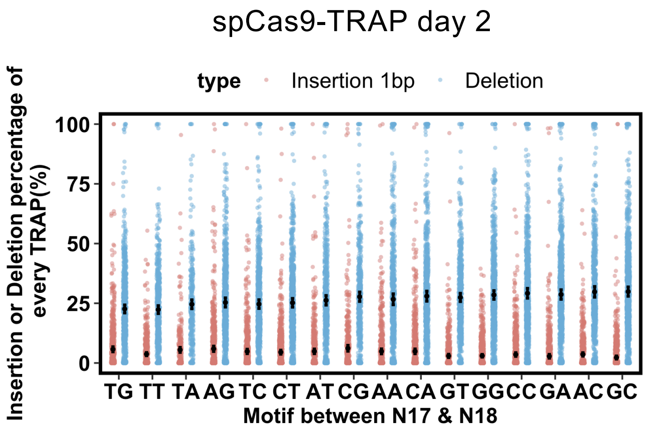
spCas9-TRAP day 2
type Insertion 1bp Deletion
Insertion or Deletion percentage of every TRAP(%)
100
75
50
25
0
TG TT TA AG TC CT AT CG AA CA GT GG CC GA AC GC
Motif between N17 & N18

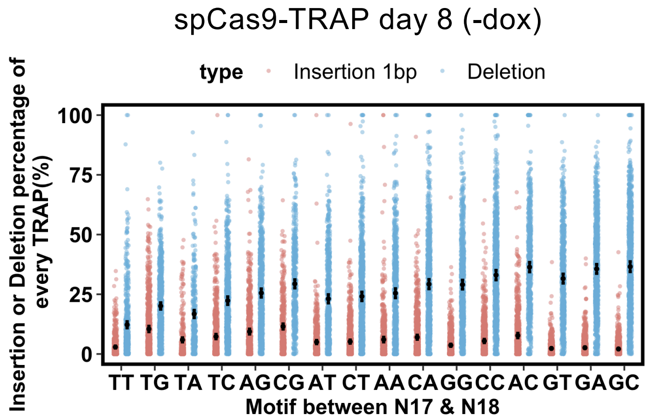
spCas9-TRAP day 8 (-dox)
type Insertion 1bp Deletion
Insertion or Deletion percentage of every TRAP(%)
100
75
50
25
0
TT TG TA TC AG CG AT CT AA CA GG CC AC GT GA GC
Motif between N17 & N18

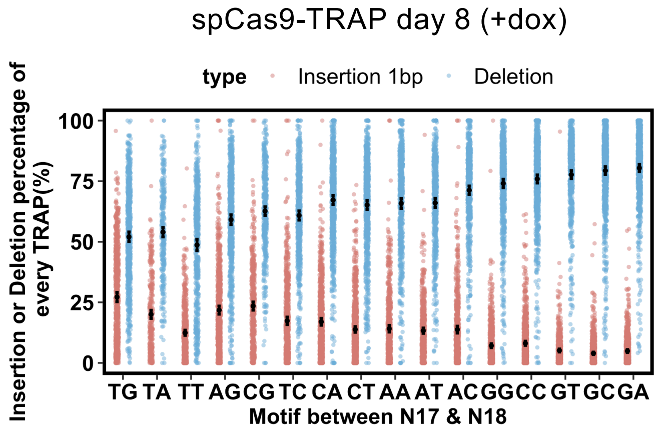
spCas9-TRAP day 8 (+dox)
type Insertion 1bp Deletion
Insertion or Deletion percentage of every TRAP(%)
100
75
50
25
0
TG TA TT AG CG TC CA CT AA AT AC GG CC GT GC GA
Motif between N17 & N18

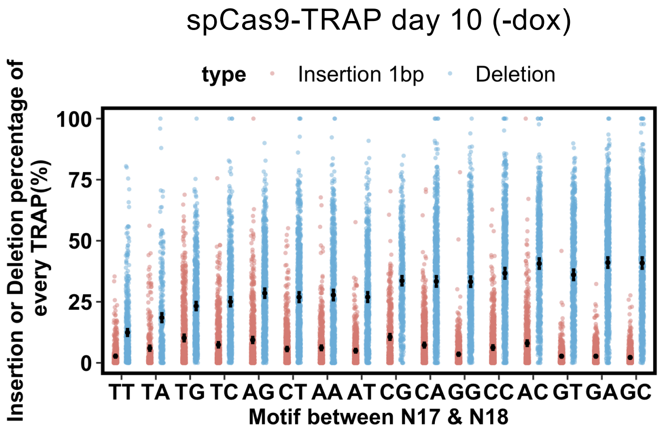
spCas9-TRAP day 10 (-dox)
type Insertion 1bp Deletion
Insertion or Deletion percentage of every TRAP(%)
100
75
50
25
0
TT TA TG TC AG CT AA AT CG CA GG CC AC GT GA GC
Motif between N17 & N18

### Fig S21

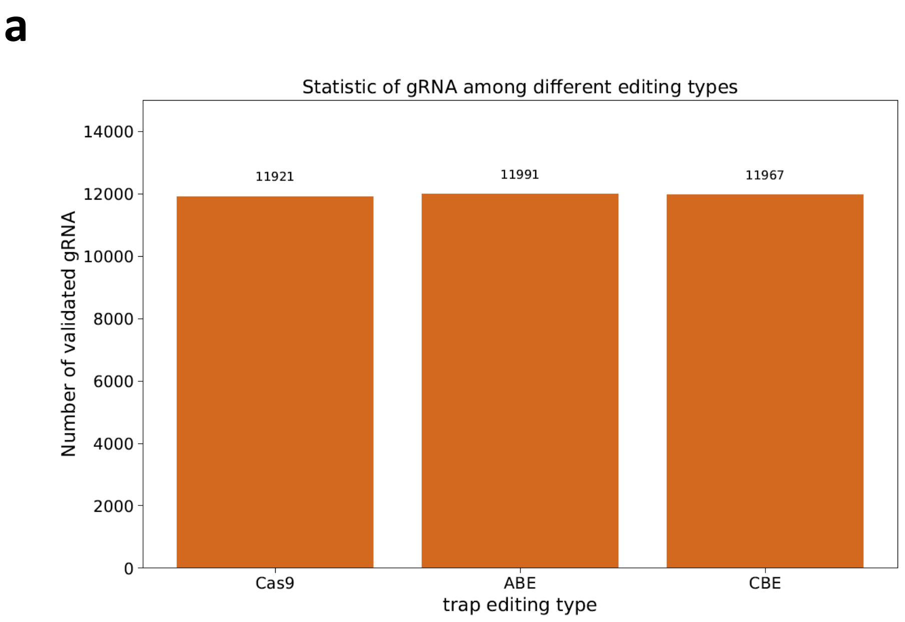
a
Statistic of gRNA among different editing types
Number of validated gRNA
14000
12000
10000
8000
6000
4000
2000
0
11921
11991
11967
Cas9
ABE
CBE
trap editing type

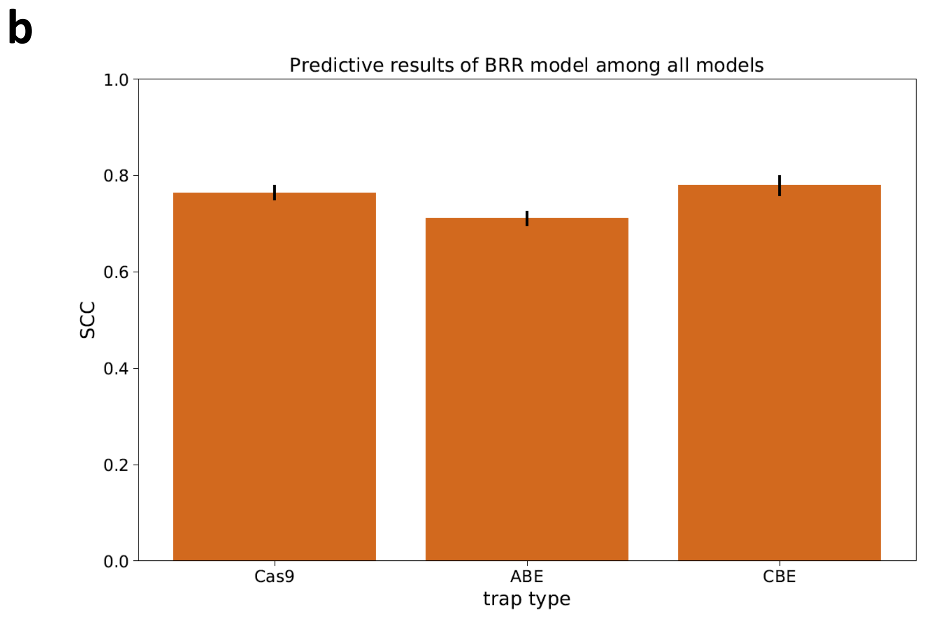
b
Predictive results of BRR model among all models
SCC
1.0
0.8
0.6
0.4
0.2
0.0
Cas9
ABE
CBE
trap type

#### Schematic illustration of the experiment

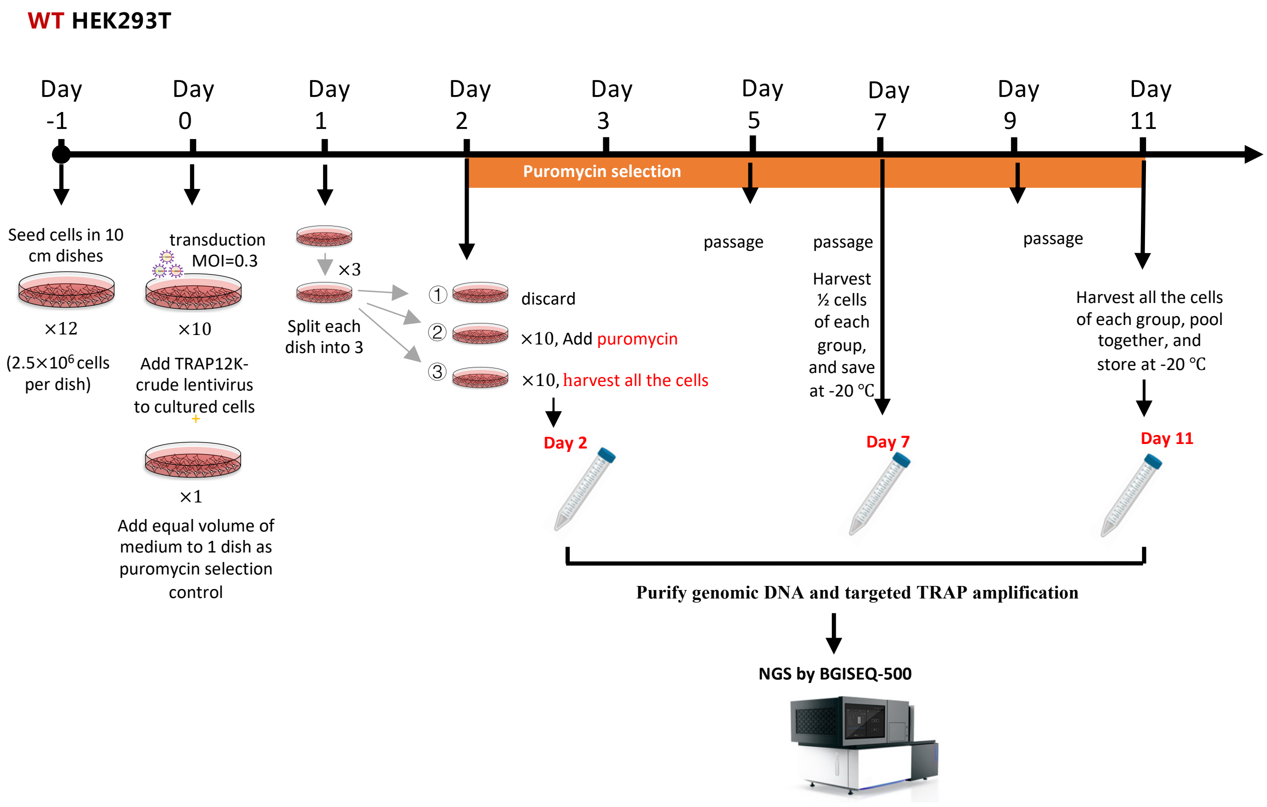

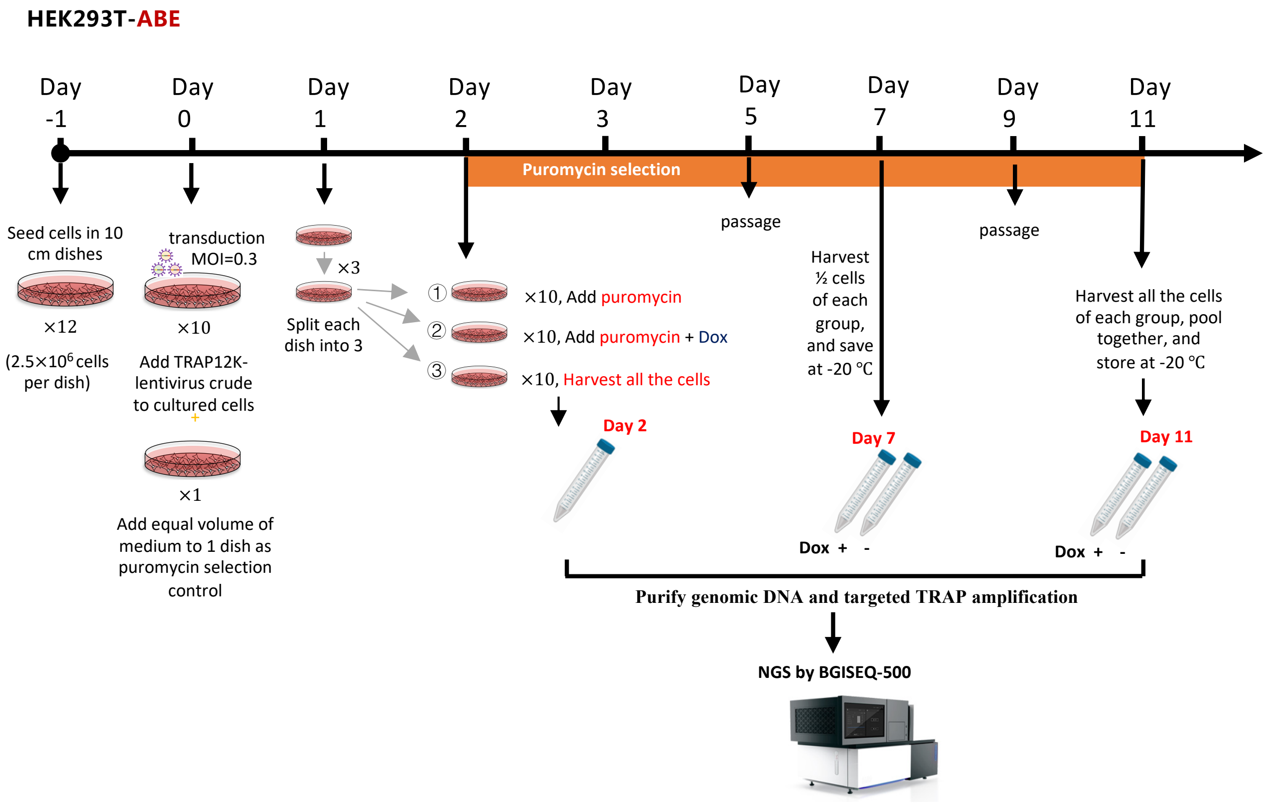

S23

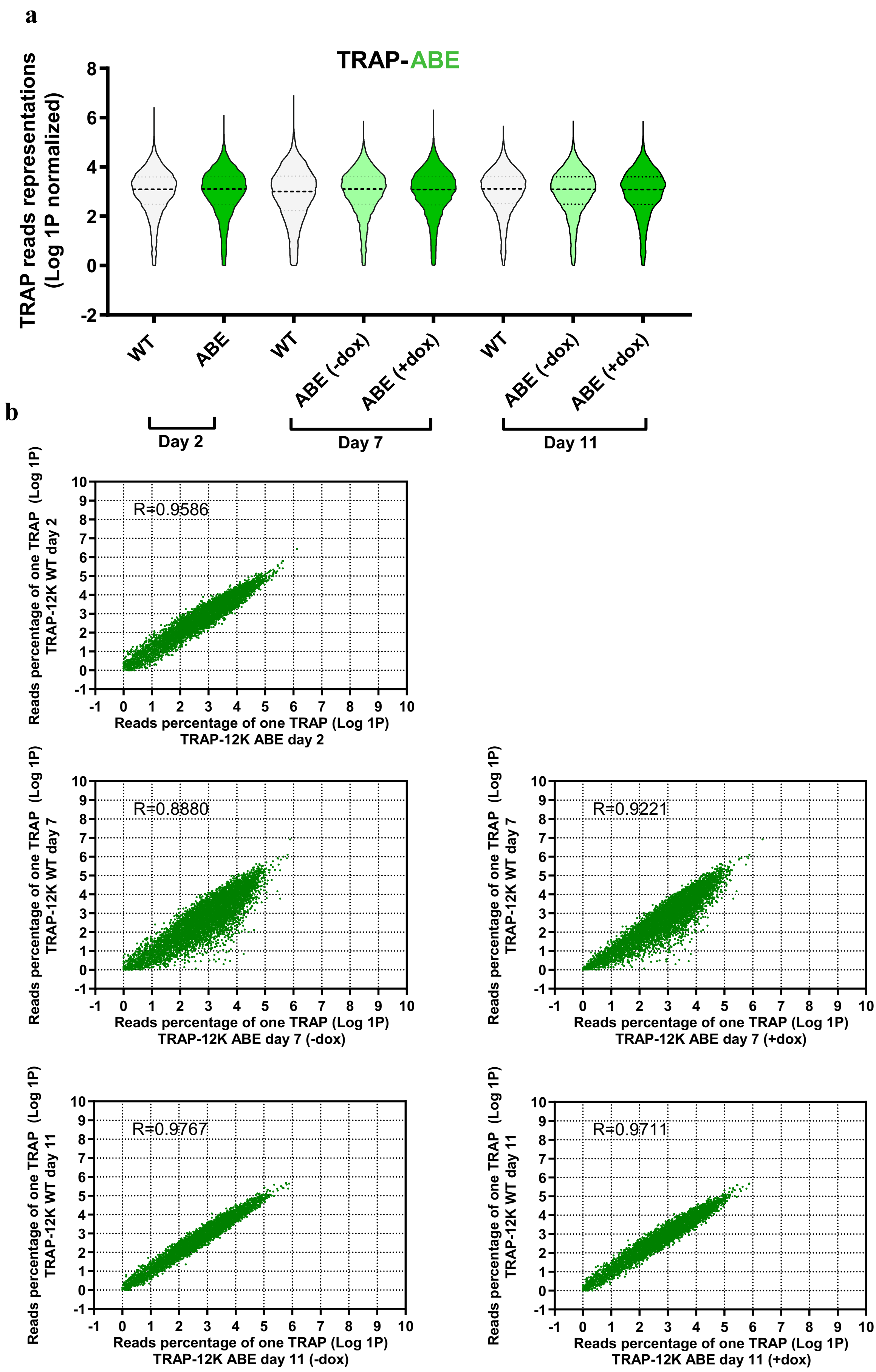
a
TRAP-ABE
TRAP reads representations (Log 1P normalized)
8
6
4
2
0
-2
WT
ABE
WT
ABE (-dox)
ABE (+dox)
WT
ABE (-dox)
ABE (+dox)
Day 2
Day 7
Day 11
b
R=0.9586
Reads percentage of one TRAP (Log 1P) TRAP-12K WT day 2
Reads percentage of one TRAP (Log 1P) TRAP-12K ABE day 2
R=0.8880
Reads percentage of one TRAP (Log 1P) TRAP-12K WT day 7
Reads percentage of one TRAP (Log 1P) TRAP-12K ABE day 7 (-dox)
R=0.9221
Reads percentage of one TRAP (Log 1P) TRAP-12K WT day 7
Reads percentage of one TRAP (Log 1P) TRAP-12K ABE day 7 (+dox)
R=0.9767
Reads percentage of one TRAP (Log 1P) TRAP-12K WT day 11
Reads percentage of one TRAP (Log 1P) TRAP-12K ABE day 11 (-dox)
R=0.9711
Reads percentage of one TRAP (Log 1P) TRAP-12K WT day 11
Reads percentage of one TRAP (Log 1P) TRAP-12K ABE day 11 (+dox)

**S24**

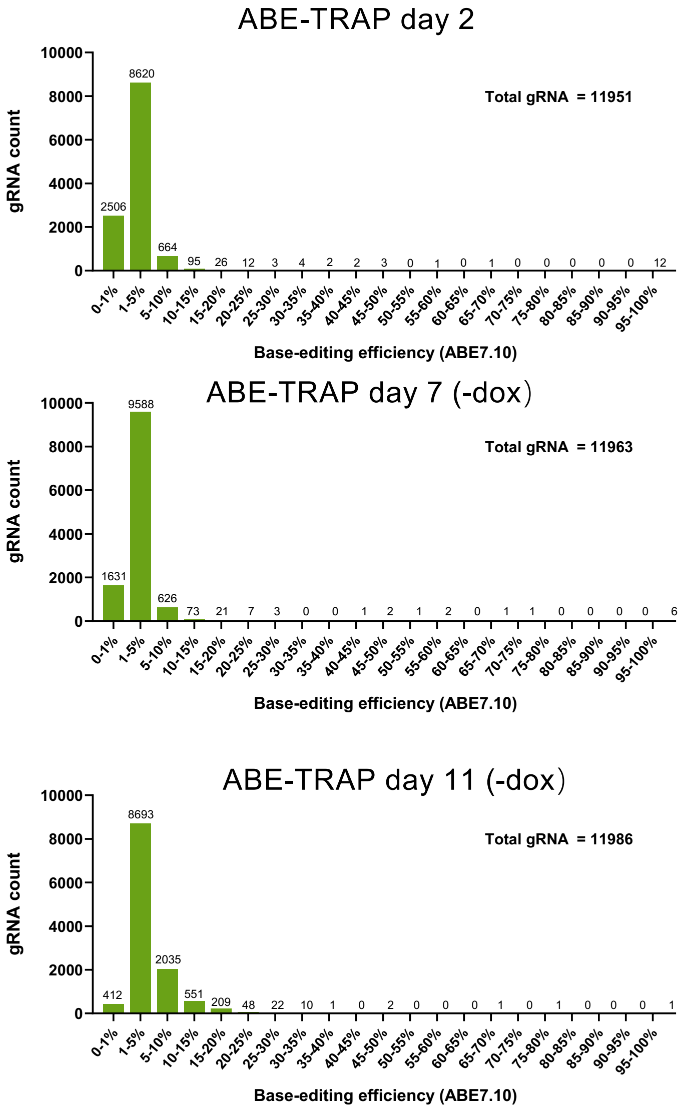
ABE-TRAP day 2
Total gRNA = 11951
gRNA count
10000
8000
6000
4000
2000
0
2506
8620
664
95
26
12
3
4
2
2
3
0
1
0
1
0
0
0
0
0
12
0-1%
1-5%
5-10%
10-15%
15-20%
20-25%
25-30%
30-35%
35-40%
40-45%
45-50%
50-55%
55-60%
60-65%
65-70%
70-75%
75-80%
80-85%
85-90%
90-95%
95-100%
Base-editing efficiency (ABE7.10)
ABE-TRAP day 7 (-dox)
Total gRNA = 11963
1631
9588
626
73
21
7
3
0
0
1
2
1
2
0
1
1
0
0
0
0
6
ABE-TRAP day 11 (-dox)
Total gRNA = 11986
412
8693
2035
551
209
48
22
10
1
0
2
0
0
0
1
0
1
0
0
0
1

**S25**

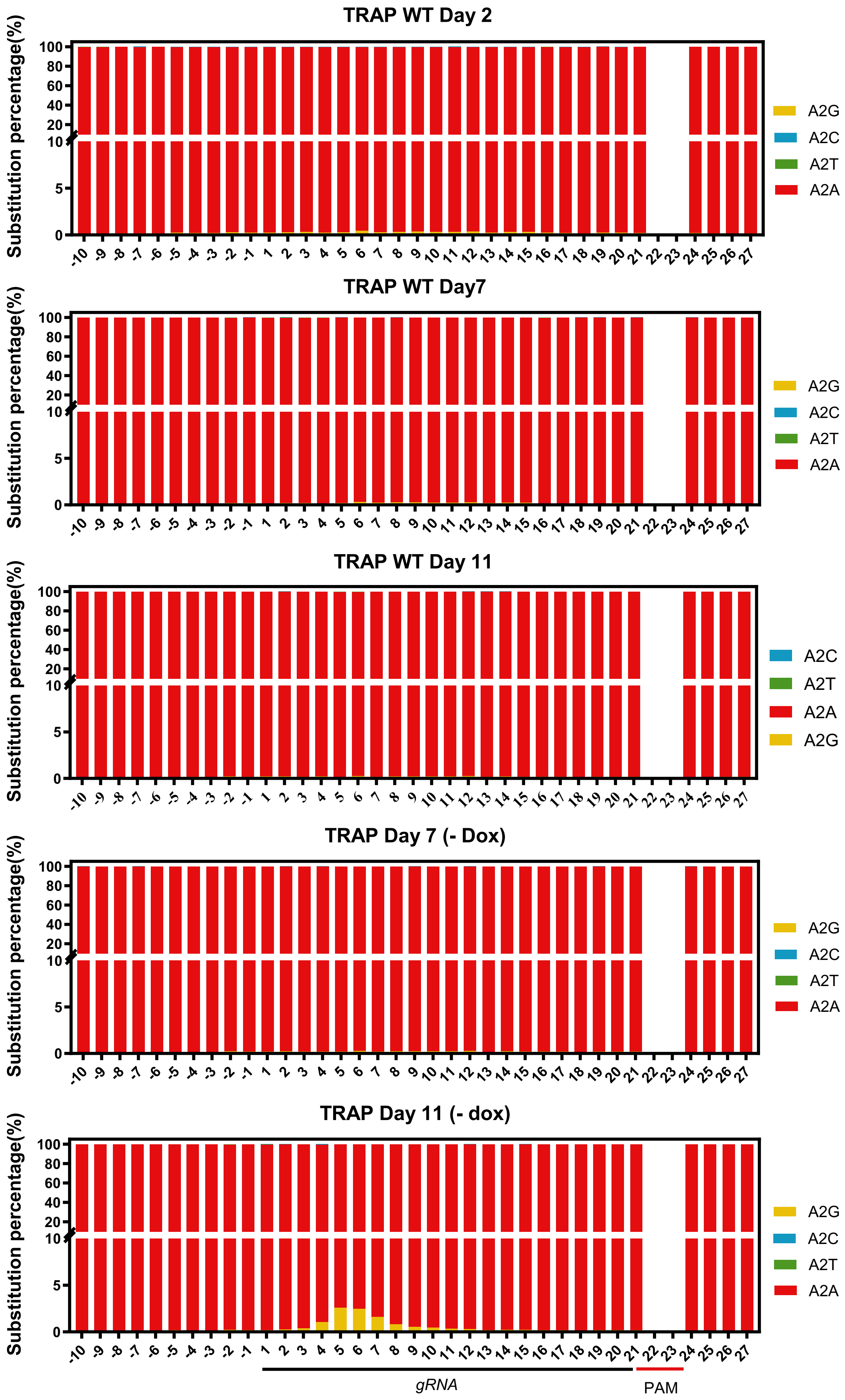
TRAP WT Day 2
TRAP WT Day7
TRAP WT Day 11
TRAP Day 7 (- Dox)
TRAP Day 11 (- dox)
Substitution percentage(%)
A2G
A2C
A2T
A2A
gRNA
PAM

**S26**

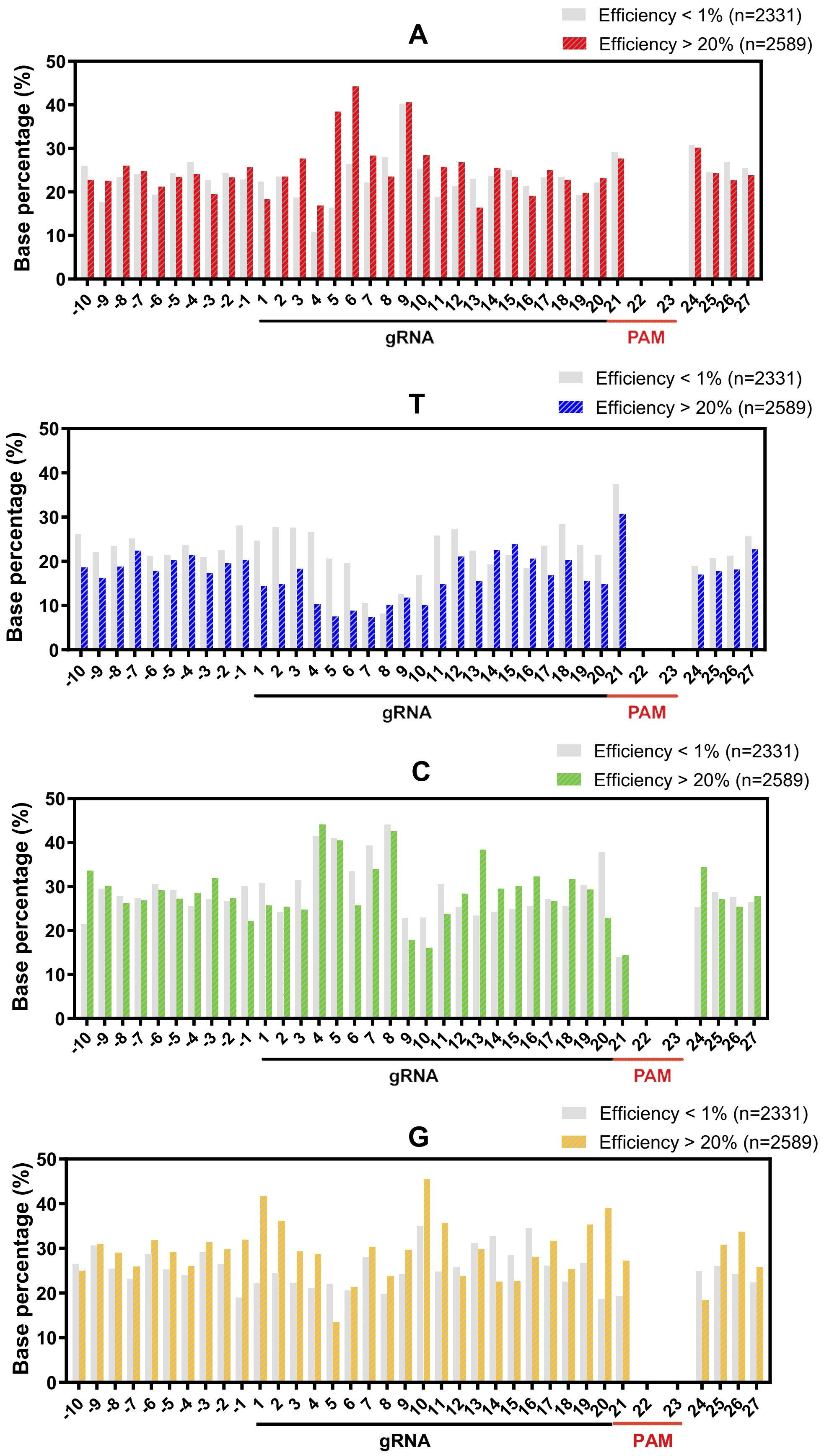
A
Efficiency < 1% (n=2331)
Efficiency > 20% (n=2589)
Base percentage (%)
gRNA
PAM
T
Efficiency < 1% (n=2331)
Efficiency > 20% (n=2589)
Base percentage (%)
gRNA
PAM
C
Efficiency < 1% (n=2331)
Efficiency > 20% (n=2589)
Base percentage (%)
gRNA
PAM
G
Efficiency < 1% (n=2331)
Efficiency > 20% (n=2589)
Base percentage (%)
gRNA
PAM

# S27

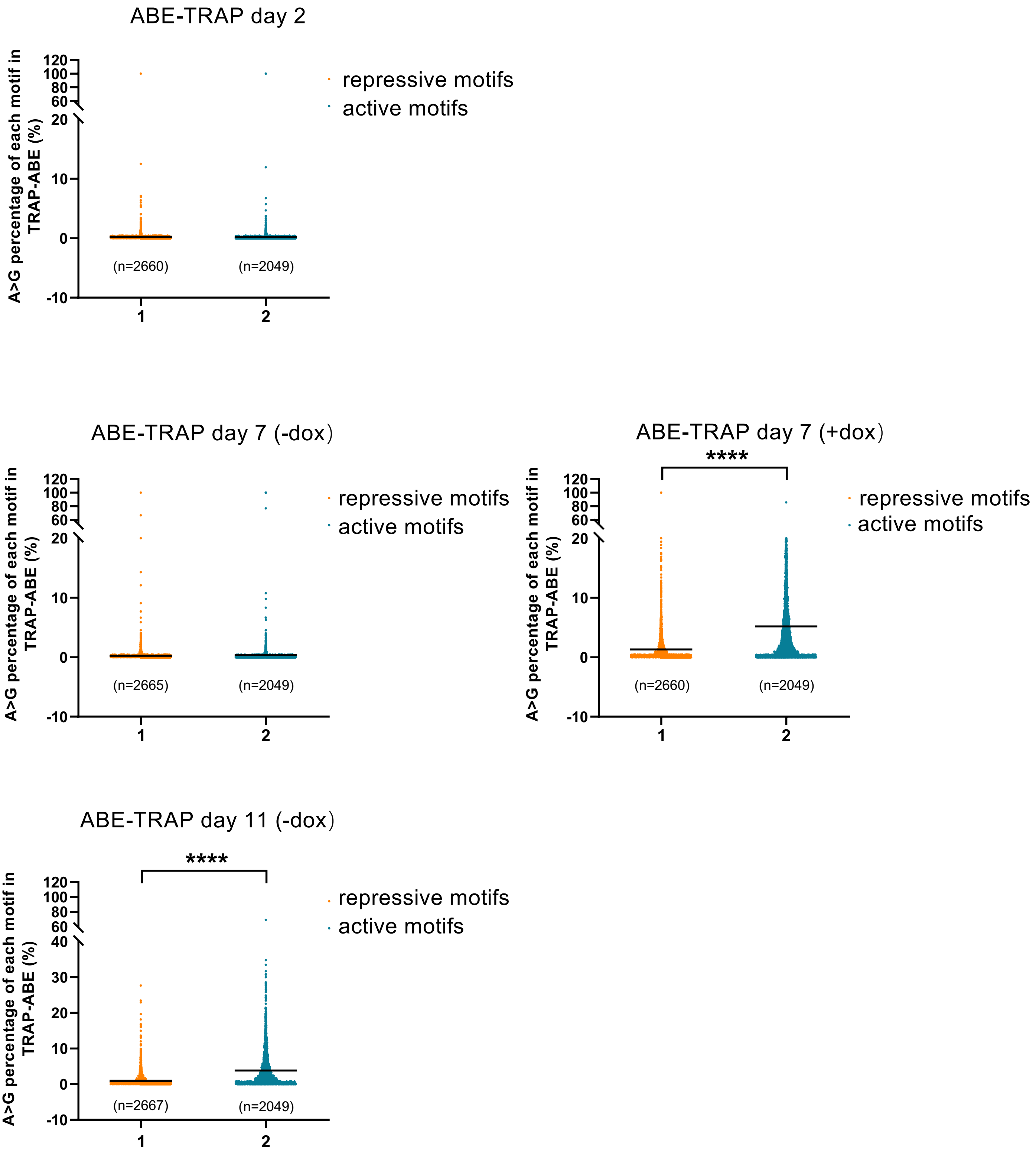
ABE-TRAP day 2
A>G percentage of each motif in TRAP-ABE (%)
repressive motifs
active motifs
(n=2660)
(n=2049)
ABE-TRAP day 7 (-dox)
A>G percentage of each motif in TRAP-ABE (%)
repressive motifs
active motifs
(n=2665)
(n=2049)
ABE-TRAP day 7 (+dox)
****
A>G percentage of each motif in TRAP-ABE (%)
repressive motifs
active motifs
(n=2660)
(n=2049)
ABE-TRAP day 11 (-dox)
****
A>G percentage of each motif in TRAP-ABE (%)
repressive motifs
active motifs
(n=2667)
(n=2049)

**S28**

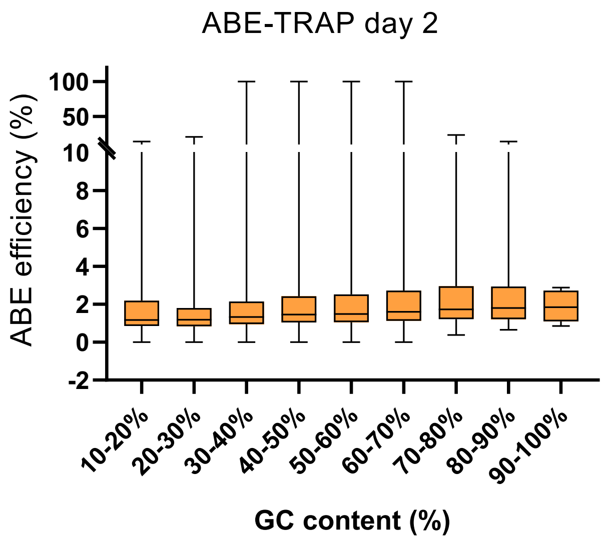
ABE-TRAP day 2
ABE efficiency (%)
100
50
10
8
6
4
2
0
-2
10-20%
20-30%
30-40%
40-50%
50-60%
60-70%
70-80%
80-90%
90-100%
GC content (%)

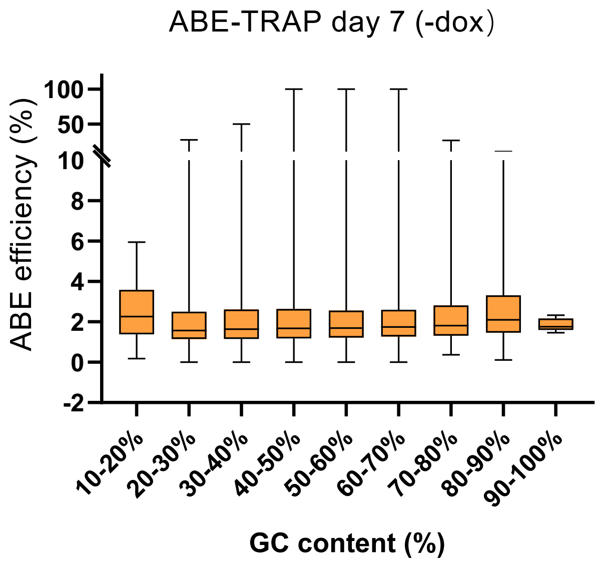
ABE-TRAP day 7 (-dox)
ABE efficiency (%)
100
50
10
8
6
4
2
0
-2
10-20%
20-30%
30-40%
40-50%
50-60%
60-70%
70-80%
80-90%
90-100%
GC content (%)

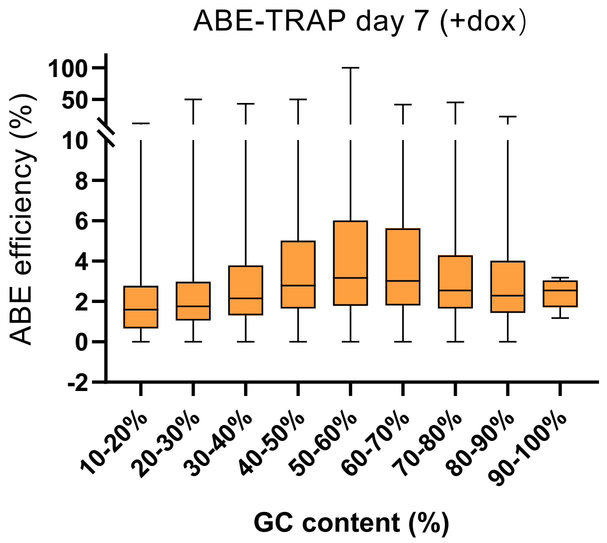
ABE-TRAP day 7 (+dox)
ABE efficiency (%)
100
50
10
8
6
4
2
0
-2
10-20%
20-30%
30-40%
40-50%
50-60%
60-70%
70-80%
80-90%
90-100%
GC content (%)

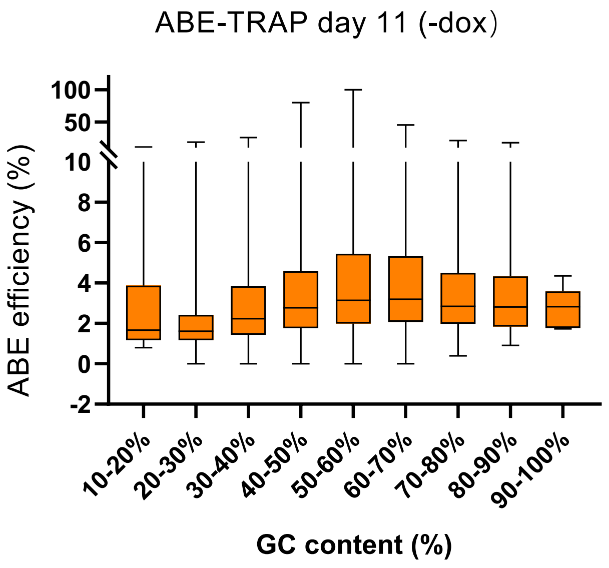
ABE-TRAP day 11 (-dox)
ABE efficiency (%)
100
50
10
8
6
4
2
0
-2
10-20%
20-30%
30-40%
40-50%
50-60%
60-70%
70-80%
80-90%
90-100%
GC content (%)

## S29

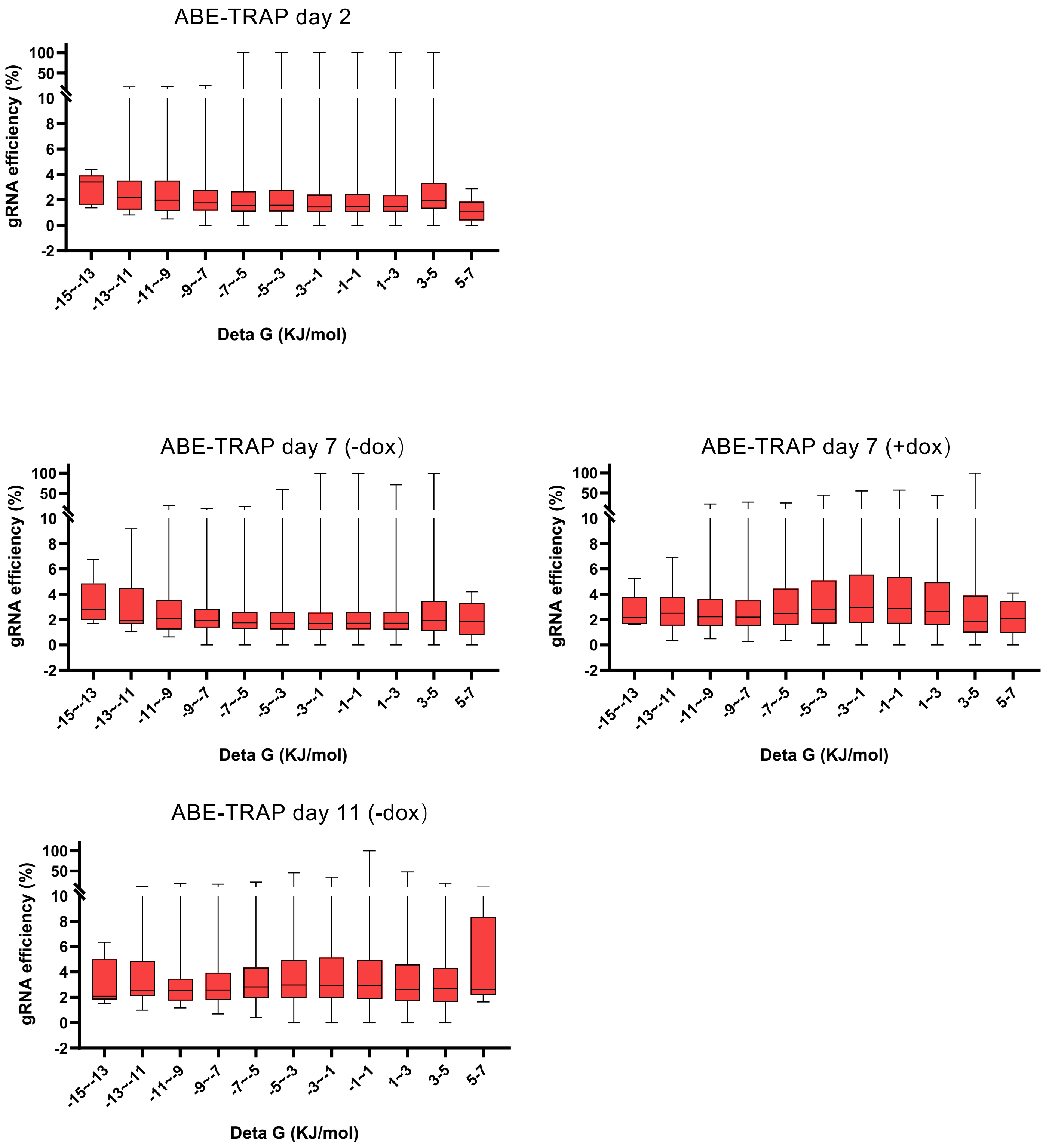
ABE-TRAP day 2
gRNA efficiency (%)
100
50
10
8
6
4
2
0
-2
-15~-13
-13~-11
-11~-9
-9~-7
-7~-5
-5~-3
-3~-1
-1~1
1~3
3-5
5-7
Deta G (KJ/mol)
ABE-TRAP day 7 (-dox)
gRNA efficiency (%)
Deta G (KJ/mol)
ABE-TRAP day 7 (+dox)
gRNA efficiency (%)
Deta G (KJ/mol)
ABE-TRAP day 11 (-dox)
gRNA efficiency (%)
Deta G (KJ/mol)

**S30**

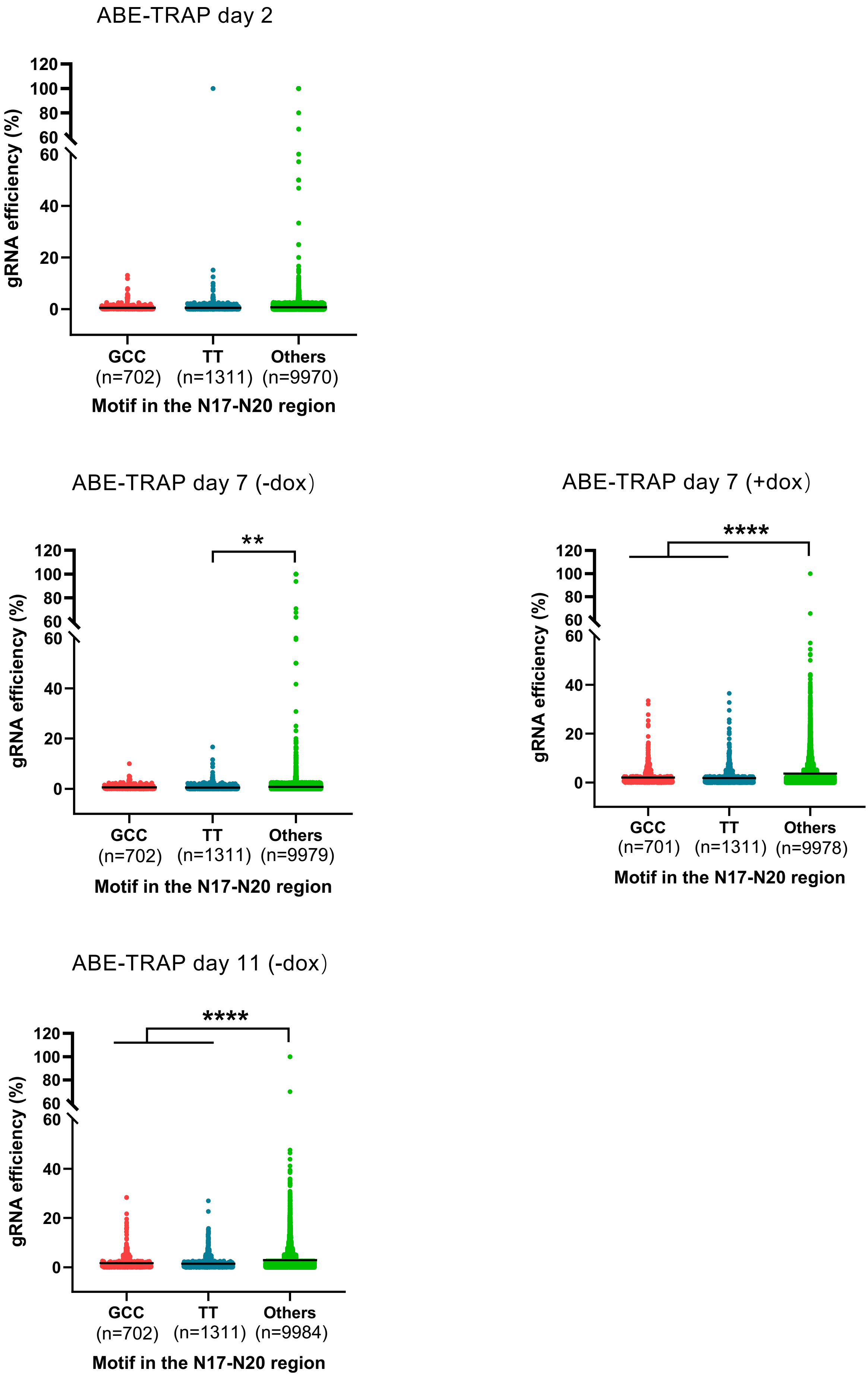
ABE-TRAP day 2
gRNA efficiency (%)
120
100
80
60
60
40
20
0
GCC (n=702)
TT (n=1311)
Others (n=9970)
Motif in the N17-N20 region
ABE-TRAP day 7 (-dox)
**
GCC (n=702)
TT (n=1311)
Others (n=9979)
Motif in the N17-N20 region
ABE-TRAP day 7 (+dox)
****
GCC (n=701)
TT (n=1311)
Others (n=9978)
Motif in the N17-N20 region
ABE-TRAP day 11 (-dox)
****
GCC (n=702)
TT (n=1311)
Others (n=9984)
Motif in the N17-N20 region

S31

a

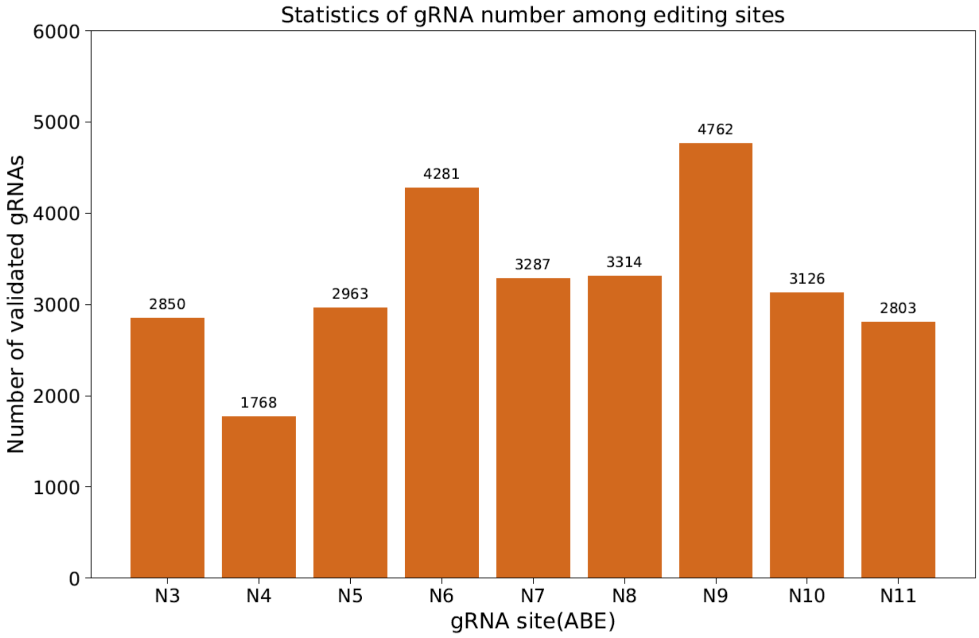
Statistics of gRNA number among editing sites
Number of validated gRNAs
0
1000
2000
3000
4000
5000
6000
2850
1768
2963
4281
3287
3314
4762
3126
2803
N3
N4
N5
N6
N7
N8
N9
N10
N11
gRNA site(ABE)

b

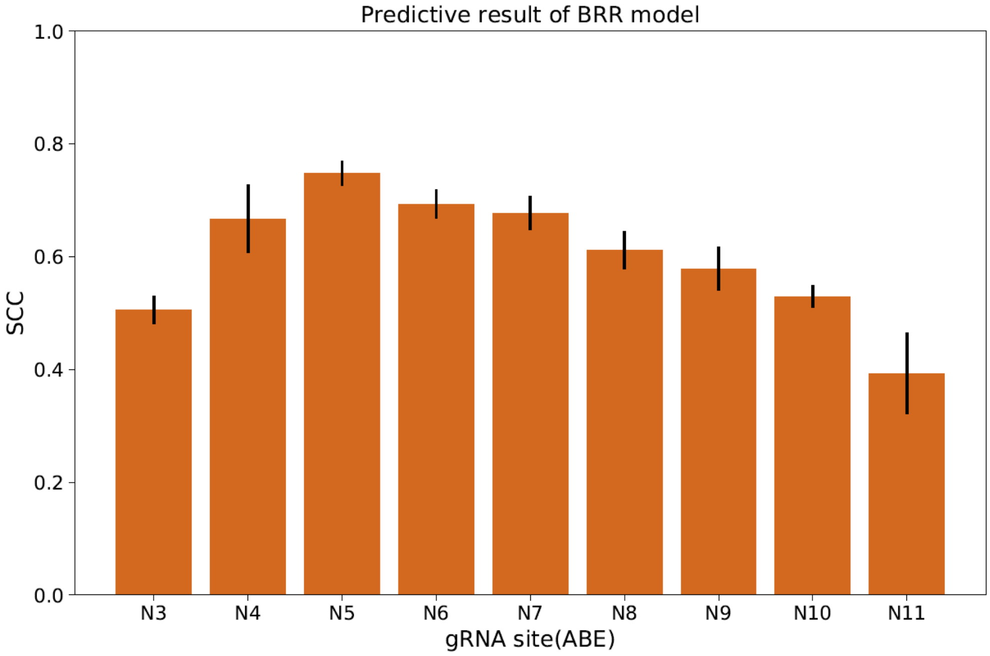
Predictive result of BRR model
SCC
0.0
0.2
0.4
0.6
0.8
1.0
N3
N4
N5
N6
N7
N8
N9
N10
N11
gRNA site(ABE)

# S32

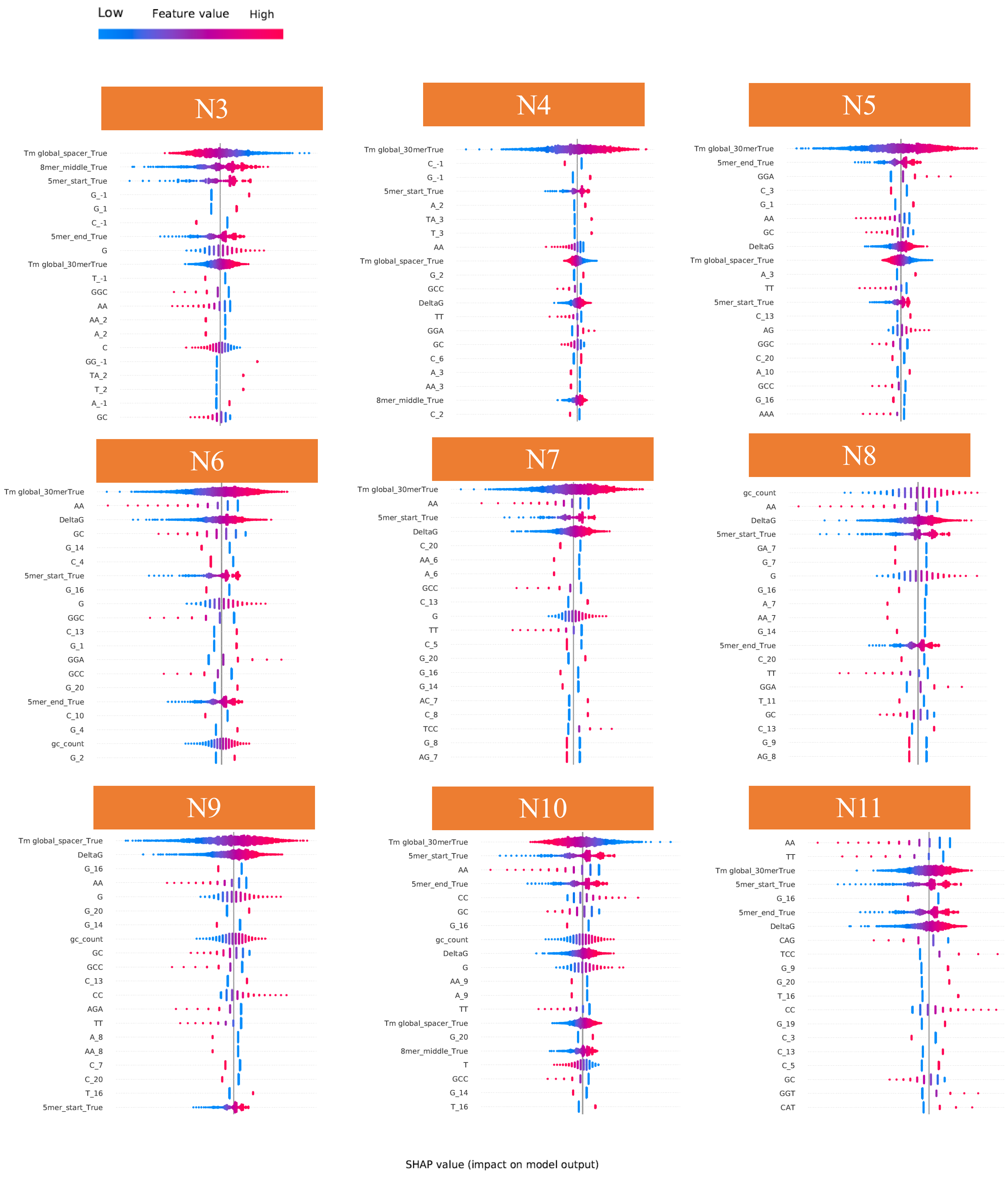
Low
Feature value
High
N3
Tm global_spacer_True
8mer_middle_True
5mer_start_True
G_-1
G_1
C_-1
5mer_end_True
G
Tm global_30merTrue
T_-1
GGC
AA
AA_2
A_2
C
GG_-1
TA_2
T_2
A_-1
GC
N4
Tm global_30merTrue
C_-1
G_-1
5mer_start_True
A_2
TA_3
T_3
AA
Tm global_spacer_True
G_2
GCC
DeltaG
TT
GGA
GC
C_6
A_3
AA_3
8mer_middle_True
C_2
N5
Tm global_30merTrue
5mer_end_True
GGA
C_3
G_1
AA
GC
DeltaG
Tm global_spacer_True
A_3
TT
5mer_start_True
C_13
AG
GGC
C_20
A_10
GCC
G_16
AAA
N6
Tm global_30merTrue
AA
DeltaG
GC
G_14
C_4
5mer_start_True
G_16
G
GGC
C_13
G_1
GGA
GCC
G_20
5mer_end_True
C_10
G_4
gc_count
G_2
N7
Tm global_30merTrue
AA
5mer_start_True
DeltaG
C_20
AA_6
A_6
GCC
C_13
G
TT
C_5
G_20
G_16
G_14
AC_7
C_8
TCC
G_8
AG_7
N8
gc_count
AA
DeltaG
5mer_start_True
GA_7
G_7
G
G_16
A_7
AA_7
G_14
5mer_end_True
C_20
TT
GGA
T_11
GC
C_13
G_9
AG_8
N9
Tm global_spacer_True
DeltaG
G_16
AA
G
G_20
G_14
gc_count
GC
GCC
C_13
CC
AGA
TT
A_8
AA_8
C_7
C_20
T_16
5mer_start_True
N10
Tm global_30merTrue
5mer_start_True
AA
5mer_end_True
CC
GC
G_16
gc_count
DeltaG
G
AA_9
A_9
TT
Tm global_spacer_True
G_20
8mer_middle_True
T
GCC
G_14
T_16
N11
AA
TT
Tm global_30merTrue
5mer_start_True
G_16
5mer_end_True
DeltaG
CAG
TCC
G_9
G_20
T_16
CC
G_19
C_3
C_13
C_5
GC
GGT
CAT
SHAP value (impact on model output)

#### Schematic illustration of the experiment

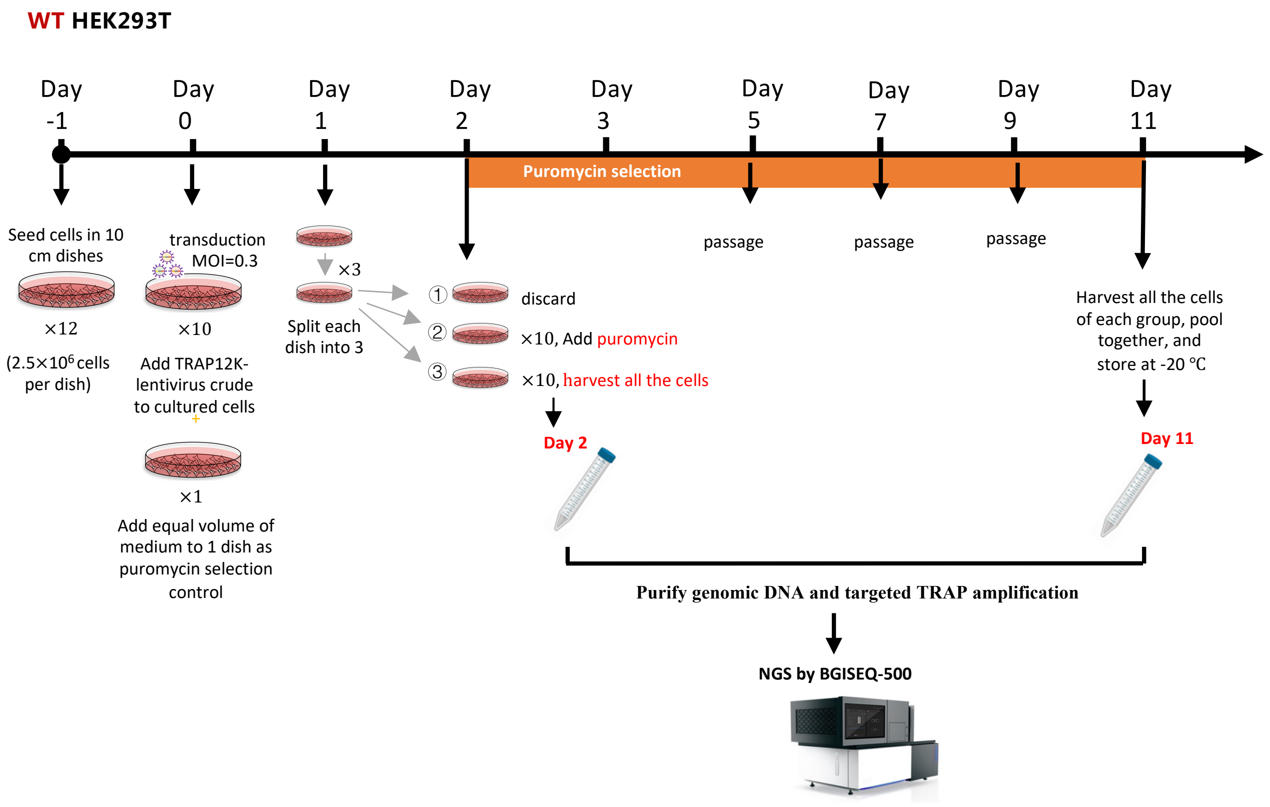

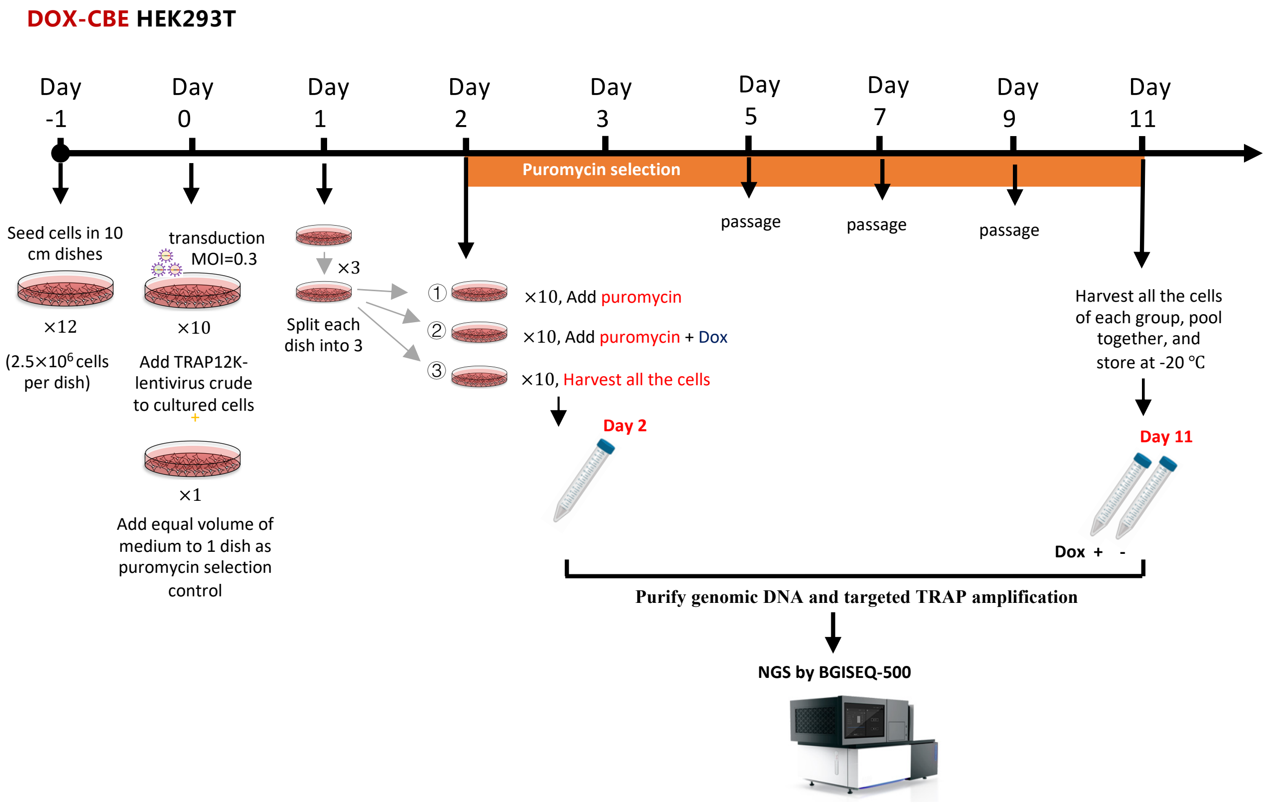

**S34**

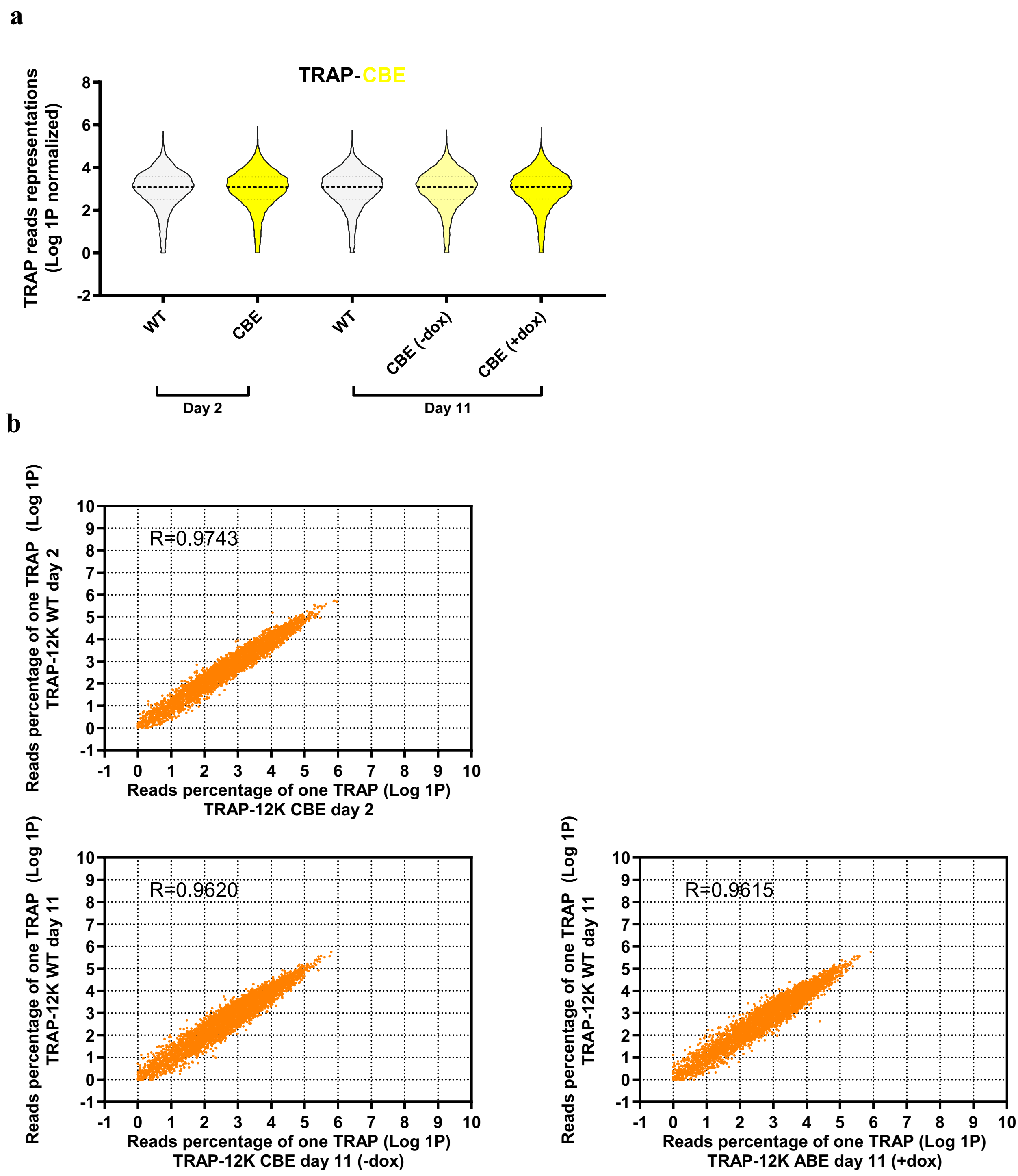
a
TRAP-CBE
TRAP reads representations (Log 1P normalized)
8
6
4
2
0
-2
WT
CBE
WT
CBE (-dox)
CBE (+dox)
Day 2
Day 11
b
R=0.9743
Reads percentage of one TRAP (Log 1P) TRAP-12K WT day 2
Reads percentage of one TRAP (Log 1P) TRAP-12K CBE day 2
R=0.9620
Reads percentage of one TRAP (Log 1P) TRAP-12K WT day 11
Reads percentage of one TRAP (Log 1P) TRAP-12K CBE day 11 (-dox)
R=0.9615
Reads percentage of one TRAP (Log 1P) TRAP-12K WT day 11
Reads percentage of one TRAP (Log 1P) TRAP-12K ABE day 11 (+dox)
-1
0
1
2
3
4
5
6
7
8
9
10

**S35**

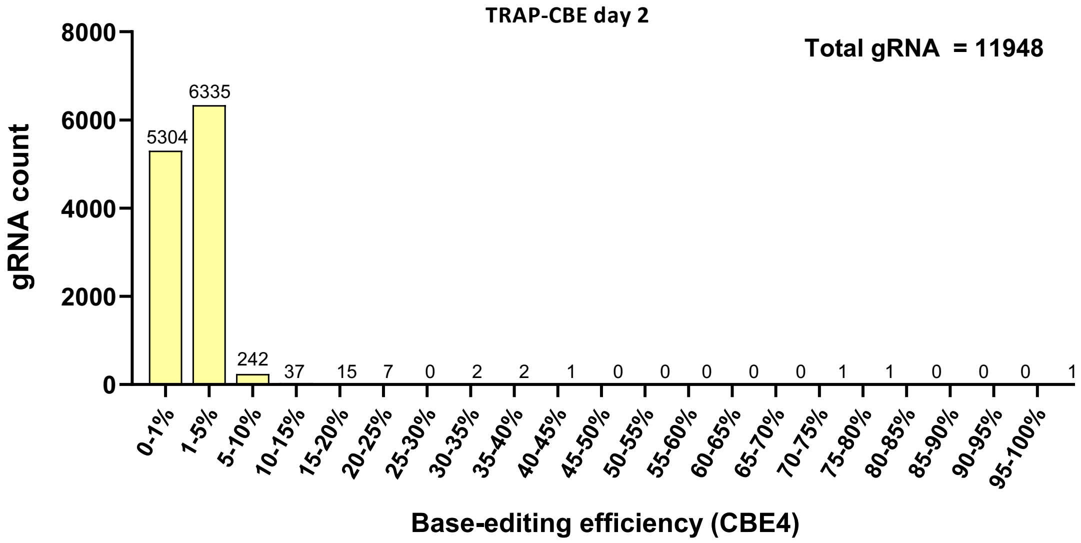
TRAP-CBE day 2
Total gRNA = 11948
gRNA count
8000
6000
4000
2000
0
5304
6335
242
37
15
7
0
2
2
1
0
0
0
0
0
1
1
0
0
0
1
0-1%
1-5%
5-10%
10-15%
15-20%
20-25%
25-30%
30-35%
35-40%
40-45%
45-50%
50-55%
55-60%
60-65%
65-70%
70-75%
75-80%
80-85%
85-90%
90-95%
95-100%
Base-editing efficiency (CBE4)

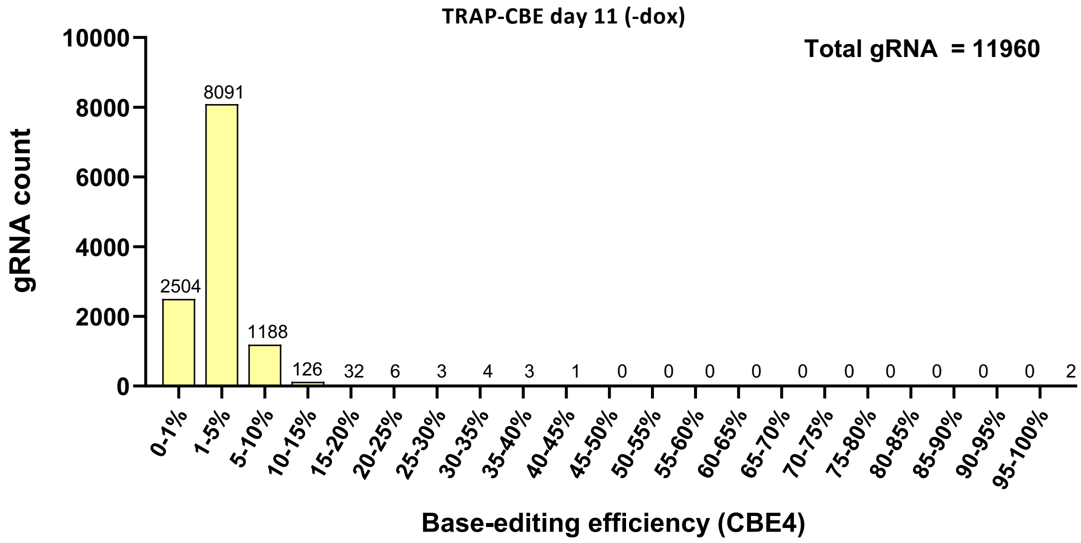
TRAP-CBE day 11 (-dox)
Total gRNA = 11960
gRNA count
10000
8000
6000
4000
2000
0
2504
8091
1188
126
32
6
3
4
3
1
0
0
0
0
0
0
0
0
0
0
2
0-1%
1-5%
5-10%
10-15%
15-20%
20-25%
25-30%
30-35%
35-40%
40-45%
45-50%
50-55%
55-60%
60-65%
65-70%
70-75%
75-80%
80-85%
85-90%
90-95%
95-100%
Base-editing efficiency (CBE4)

**S36**

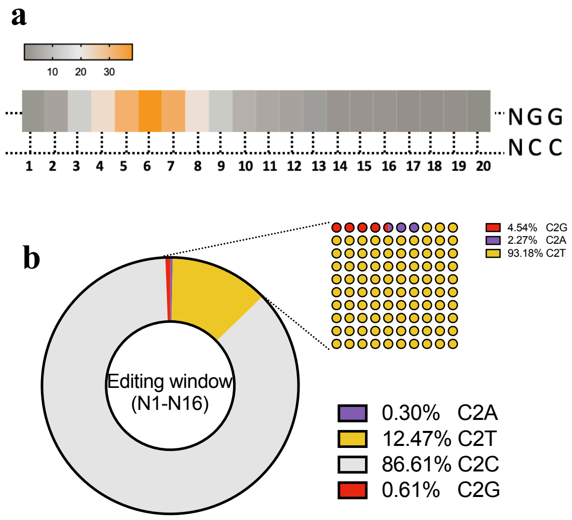
a
10
20
30
NGG
NCC
b
Editing window
(N1-N16)
4.54% C2G
2.27% C2A
93.18% C2T
0.30% C2A
12.47% C2T
86.61% C2C
0.61% C2G

### Examples of STOP codon conversion in TRAP-CBE data

| TRAP ID | Selected data of TRAP-CBE mediated STOP codon conversion (CAG>TAG) | Obs.% |
|---|---|---|
| | A2ML2 — Ser Ser Gln Arg Ser Ile Ile Trp | |
| 3 | CAAGGGCAT TCG AGC CAG CGT TCC ATT ATC TGG AGGC | WT |
| | CAAGGGCAT TCG AGC TAG CGT TCC ATT ATC TGG AGGC | 34.86 |
| | CAAGGGCAT TCG AGT TAG CGT TCC ATT ATC TGG AGGC | 13.14 |
| | CAAGGGCAT TTG AGC TAG CGT TCC ATT ATC TGG AGGC | 7.43 |
| | CAAGGGCAT TTG AGT TAG CGT TCC ATT ATC TGG AGGC | 5.71 |
| | CAAGGGCAT TCG AGC TAG CGT TTC ATT ATC TGG AGGC | 2.86 |
| | CAAGGGCAT TTG AGT TAG CGT TTC ATT ATC TGG AGGC | 2.29 |
| | CAAGGGCAT CCG AGC TAG CGT TCC ATT ATC TGG AGGC | 0.57 |
| | CAAGGGCAT TCG AGG TAG CGT TCC ATT ATC TGG AGGC | 0.57 |
| | CAAGGGCAT TCG AGC CAG CGT TTC ATT ATC TGG AGGC | 6.29 |
| | CAAGAGCAT TCG AGC CAG CGT TCC ATT TTC TGA AGGC | 5.14 |
| | CAAGGGCAT TCG AGC CAG CGT TCC ATT ATC TAG AGGC | 4.00 |
| | CAAGGGCAT TCG AGT TAA CGT TCC ATT ATC TGG AGGC | 2.86 |
| | CAAGGGCAT TTG AGC CAG CGT TCC ATT ATC TGG AGGC | 2.86 |
| | CAAGGGCAT TCG AGC CAG CGT TCC ATT ATC TGG AGTC | 1.71 |
| | CAAGGGCAT TGG AGC CAG CGT TCC ATT ATC TGG AGGC | 1.14 |
| | CAAGGGCAT TCG AGC CAG CGT TGC ATT ATC TGG AGGC | 1.14 |
| | CAAGGGCAT ACG AGC CAG CGT TCC ATT ATC TGG AGGC | 0.57 |
| | | ...... |

| TRAP ID | Selected data of TRAP-CBE mediated STOP codon conversion (CGA>TGA) | Obs.% |
|---|---|---|
| | ABCB5 — Tyr Arg Asp His Ile Gly Val | |
| 61 | ATGTGCGGCAT TAT CGA GAC CAT ATT GGA GTG GTTAG | WT |
| | ATGTGCGGCAT TAT TGA GAC CAT ATT GGA GTG GTTAG | 46.40 |
| | ATGTGCGGCAT TAT TGA GAT CAT ATT GGA GTG GTTAG | 1.99 |
| | ATGTGTGGCAT TAT TGA GAC CAT ATT GGA GTG GTTAG | 1.87 |
| | ATGTGCGGCAT TAT TGA GAT TAT ATT GGA GTG GTTAG | 1.76 |
| | ATGTGCGGCAT TAT TGA GAC TAT ATT GGA GTG GTTAG | 1.58 |
| | ATATGCGGCAT TAT TGA GAC CAT ATT GGA GTG GTTAG | 0.70 |
| | ATGTGCGGCAT TAT TGA GAC CAT ATT GGA GTG GTTAA | 0.53 |
| | ATGAGCGGCAT TAT TGA GAC CAT ATT GGA GTG GTTAG | 0.53 |
| | ATGTGCGGCAT TAT TGA GAG TAT ATT GGA GTG GTTAG | 0.12 |
| | ATGTGCGGCAT TAT CGA GAT TAT ATT GGA GTG GTTAG | 0.12 |
| | ATGTCCGGCAT TAT TGA GAC CAT ATT GGA GTG GTTAG | 0.12 |
| | ATGTGCGGCAC TAT TGA GAC CAT ATT GGA GTG GTTAG | 0.06 |
| | ATGTGCGGCAT TAG TGA GAC CAT ATT GGA GTG GTTAG | 0.06 |
| | ATGTGCGGCAT TGT TGA GAC CAT ATT GGA GTG GTTAG | 0.06 |
| | ATGTGCGGCAG TAT TGA GAC CAT ATT GGA GTG GTTAG | 0.06 |
| | ACGTGCGGCAA TAC TGA GAC GAT ATT GGA GTG GTTAG | 0.06 |
| | ATGTGCGGCAT TAG TGA GAC TAT ATT GGA GTG GTTAG | 0.06 |
| | | ...... |

| TRAP ID | Selected data of TRAP-CBE mediated STOP codon conversion (CAA>TAA) | Obs.% |
|---|---|---|
| | ABCB4 — Gln Pro Gln Ile Leu Leu Leu | |
| 56 | CCCTCATCAGA CAA CCT CAA ATC CTC CTG TTG GATGA | WT |
| | CCCTCATCAGA CAA TTT TAA ATC CTC CTG TTG GATGA | 31.19 |
| | CCCTCATCAGA TAA TTT TAA ATC CTC CTG TTG GATGA | 5.75 |
| | CCCTCATCAGA CAA TTT TAA ATT CTC CTG TTG GATGA | 3.45 |
| | CCCTCATCAGA CAA CTT TAA ATC CTC CTG TTG GATGA | 3.07 |
| | CCCTCATCAGA CAA CCT TAA ATC CTC CTG TTG GATGA | 1.46 |
| | CCCTTATCAGA CAA TTT TAA ATC CTC CTG TTG GATGA | 1.23 |
| | CCCTTATTAGA CAA TTT TAA ATC CTC CTG TTG GATGA | 0.92 |
| | CCCTCATTAGA CAA TTT TAA ATC CTC CTG TTG GATGA | 0.92 |
| | CCCTTATTAGA TAA TTT TAA ATC CTC CTG TTG GATGA | 0.69 |
| | CCCTGATCAGA CAA TTT TAA ATC CTC CTG TTG GATGA | 0.61 |
| | CCCTCATCAGA CAA CCT TAA ATC CTC CTA TTG GATGA | 0.46 |
| | CCCTCATCAGA CAA TAT TAA ATC CTC CTG TTG GATGA | 0.38 |
| | TCCTCATCAGA TAA TTT TAA ATC CTC CTG TTG GATGA | 0.31 |
| | CCCTCATTAGA CAA TTT TAA ATC CTC CTT TTG GATGA | 0.31 |
| | TTTTTATTAGA TAA TTT TAA ATC CTC CTG TTG GATGA | 0.31 |
| | CCCTCATTAGA CAA TTT TAA ATT CTC CTG TTG GATGA | 0.23 |
| | CCCTCATCAGA CAA TTT TAA ATT CTT CTG TTG GATGA | 0.23 |
| | | ...... |

| TRAP ID | Selected data of TRAP-CBE mediated STOP codon conversion (CCA>TTA/TCA) | Obs.% |
|---|---|---|
| | A4GALT sense — Lys Lys Trp Asp Gln Trp Pro | |
| 5 | TCCTCAAAGTA CTT CTT CCA GTC CTG CCA GGG GATGG | WT |
| | TCCTCAAAGTA CTT CTT CCA GTC CTG CCA GGG GATGG | 28.38 |
| | TCCTCAAAGTA CTT TTT TTA GTC CTG CCA GGG GATGG | 24.55 |
| | TCCTCAAAGTA CTT TTT TCA GTC CTG CCA GGG GATGG | 18.14 |
| | TCCTCAAAGTA CTT TTT CCA GTC CTG CCA GGG GATGG | 15.09 |
| | TCCTCAAAGTA CTT TTT TTA GTT CTG CCA GGG GATGG | 1.25 |
| | TCCTCAAAGTA CTT CTT TCA GTC CTG CCA GGG GATGG | 1.09 |
| | TCCTTAAAGTA CTT TTT TTA GTT TTG CCA GGG GATGG | 1.02 |
| | TCCTCAAAGTA CTT TTT TTA GTT TTG CCA GGG GATGG | 0.78 |
| | TCCTCAAAGTA CTT ATT TTA GTC CTG CCA GGG GATGG | 0.78 |
| | TCCTCAAAGTA CTT TTT TGA GTC CTG CCA GGG GATGG | 0.70 |
| | TCCTCAAAGTA CTT ATT TCA GTC CTG CCA GGG GATGG | 0.55 |
| | TCCTCAAAGTA CTT TTT ATA GTC CTG CCA GGG GATGG | 0.55 |
| | TCCTCAAAGTA CTT CTG GCA GTC CTG CCA GGG GATGG | 0.55 |
| | TCCTCAAAGTA CTT CTT TTA GTT TTG CCA GGG GATGG | 0.55 |
| | TCCTCAAAATA CTT TTT TTA GTC CTG CCA GGG GATGG | 0.47 |
| | TCCTCAAAGTA CTT CTT TTA GTT CTG CCA GGG GATGG | 0.39 |
| | TCCTCAAAGTA CTT TTT GCA GTC CTG CCA GGG GATGG | 0.31 |
| | | ...... |

| TRAP ID | Selected data of TRAP-CBE mediated STOP codon conversion (CCA>CTA/TTA) | Obs.% |
|---|---|---|
| | ADRB1 sense — Arg Trp Trp His Met Leu Ile | |
| 339 | CGCTCTCCGC CCG CCA CCA GTG CAT GAG GAT GGGCAG | WT |
| | CGCTCTCCGC CCG CCA CCA GTG CAT GAG GAT GGGCAG | 35.42 |
| | CGCTCTCCGC CCG CTA TTA GTG CAT GAG GAT GGGCAG | 28.59 |
| | CGCTCTCCGC CCG CCA TTA GTG CAT GAG GAT GGGCAG | 6.45 |
| | CGCTCTCCGC CCG TTA TTA GTG CAT GAG GAT GGGCAG | 6.35 |
| | CGCTCTCCGC CCG CCA TCA GTG CAT GAG GAT GGGCAG | 5.68 |
| | CGCTCTCCGC CCG CCA GCA GTG CAT GAG GAT GGGCAG | 2.31 |
| | CGCTCTCCAC CCG CTA TTA GTG CAT GAG GAT GGGCAG | 1.06 |
| | CGCTCTCCGC CCG CTA TCA GTG CAT GAG GAT GGGCAG | 0.96 |
| | CGCTTTCCGC CCG CTA TTA GTG CAT GAG GAT GGGCAG | 0.96 |
| | CGCTCTCAGC CCG CCA TTA GTG CAT GAG GAT GGGCAG | 0.96 |
| | CGCTTTTTGC CCG TTA TTA GTG CAT GAG GAT GGGCAG | 0.96 |
| | CGCTCTCCGC CCG CTA CCA GTG CAT GAG GAT GGGCAG | 0.77 |
| | CGCTTTTTGC CCG CTA TTA GTG CAT GAG GAT GGGCAG | 0.77 |
| | CGCTCTTCGC CCG TTA TTA GTG CAT GAG GAT GGGCAG | 0.58 |
| | CGCCCTCCGC CCG CTA TTA GTG CAT GAG GAT GGGCAG | 0.48 |
| | CGCTTTTCGC CCG CTA TTA GTG CAT GAG GAT GGGCAG | 0.48 |
| | CGCTCTCCGC CCG CCA CTA GTG CAT GAG GAT GGGCAG | 0.48 |
| | | ...... |

| TRAP ID | Selected data of TRAP-CBE mediated STOP codon conversion (CCA>TTA/CTA) | Obs.% |
|---|---|---|
| | A4GALT sense — Asn Trp Val His Val Ala Tyr | |
| 4 | GGCTCTTCTT GTT CCA CAC GTG GAC AGC ATA GGTGGC | WT |
| | GGCTCTTCTT GTT TTA TAC GTG GAC AGC ATA GGTGGC | 49.23 |
| | GGCTCTTCTT GTT TTA CAC GTG GAC AGC ATA GGTGGC | 8.95 |
| | GGCTCTTCTT GTT TCA CAC GTG GAC AGC ATA GGTGGC | 7.40 |
| | GGCTCTTCTT GTT TTA TAT GTG GAC AGC ATA GGTGGC | 6.71 |
| | GGCTTTTCTT GTT TTA TAC GTG GAC AGC ATA GGTGGC | 3.27 |
| | GGCTCTTCTT GTT CCA TAC GTG GAC AGC ATA GGTGGC | 2.07 |
| | GACTCTTCTT GTT TTA TAC GTG GAC AGC ATA GGTGGC | 1.89 |
| | GGCTCTTCTT GTT ACA CAC GTG GAC AGC ATA GGTGGC | 1.72 |
| | GGCTTTTTTT GTT TTA TAC GTG GAC AGC ATA GGTGGC | 1.72 |
| | GGCTCTTTTT GTT TTA TAT GTG GAC AGC ATA GGTGGC | 1.55 |
| | GGCTTTTTTT GTT TTA TAT GTG GAC AGC ATA GGTGGC | 1.38 |
| | GGCTCTTTTT GTT TTA TAC GTG GAC AGC ATA GGTGGC | 1.03 |
| | GACTCTTCTT GTT TTA CAC GTG GAC AGC ATA GGTGGC | 1.03 |
| | GGCTCTTCTT GTT CTA TAC GTG GAC AGC ATA GGTGGC | 0.86 |
| | GGCTCTTCTT GTT CTA CAC GTG GAC AGC ATA GGTGGC | 0.86 |
| | GGCTCTTCTT GTT ATA TAC GTG GAC AGC ATA GGTGGC | 0.86 |
| | GGCTTTTCTT GTT TTA TAT GTG GAC AGC ATA GGTGGC | 0.69 |
| | | ...... |

**S38**

**a**

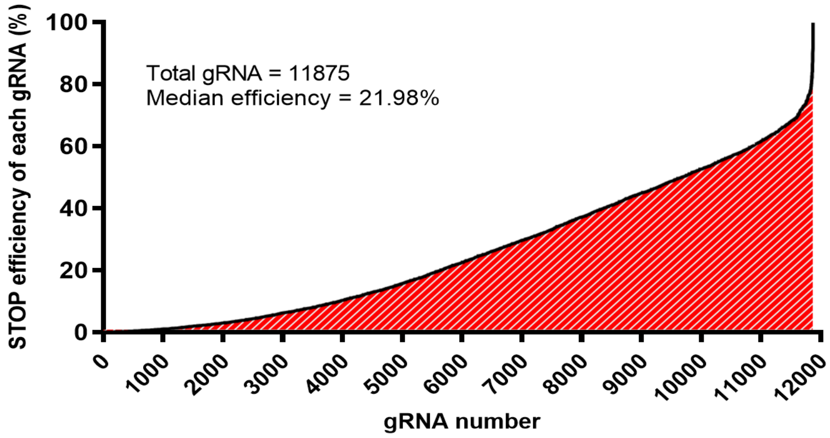

STOP efficiency of each gRNA (%)
100
80
60
40
20
0
Total gRNA = 11875
Median efficiency = 21.98%
0
1000
2000
3000
4000
5000
6000
7000
8000
9000
10000
11000
12000
gRNA number

**b**

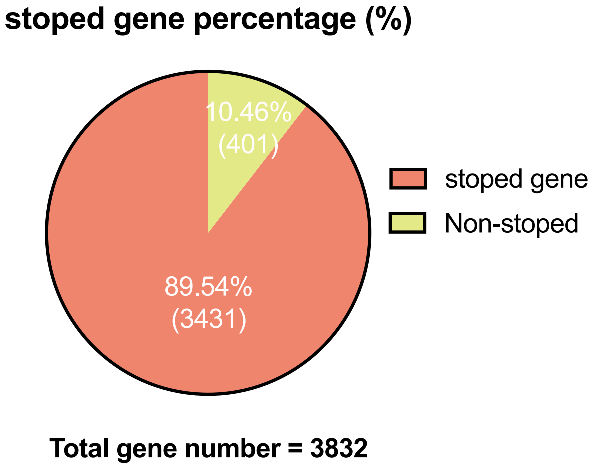

stoped gene percentage (%)
10.46%
(401)
89.54%
(3431)
stoped gene
Non-stoped
Total gene number = 3832

**S39**

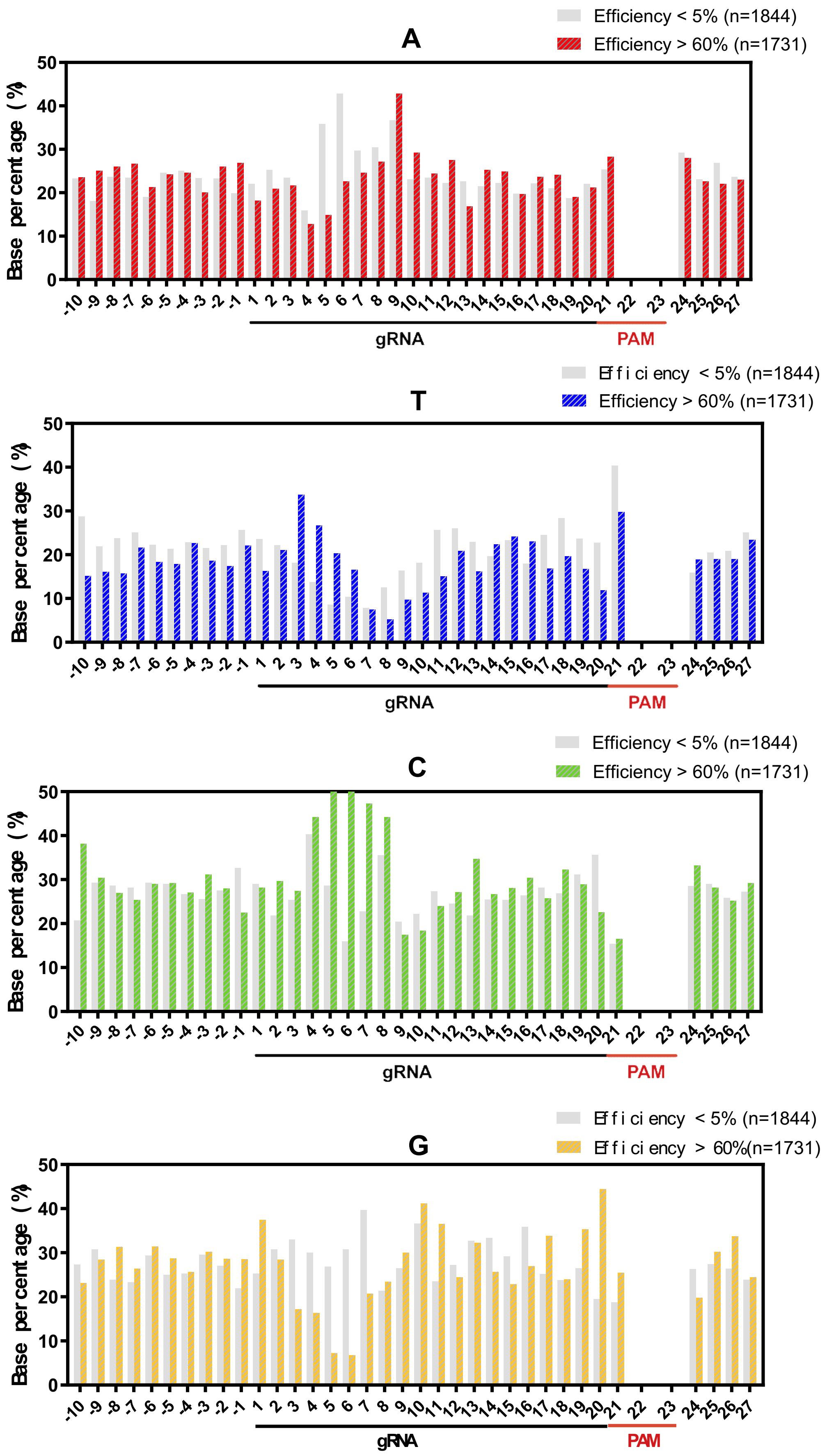
A
Efficiency < 5% (n=1844)
Efficiency > 60% (n=1731)
Base percentage (%)
50
40
30
20
10
0
-10 -9 -8 -7 -6 -5 -4 -3 -2 -1 1 2 3 4 5 6 7 8 9 10 11 12 13 14 15 16 17 18 19 20 21 22 23 24 25 26 27
gRNA
PAM
T
Efficiency < 5% (n=1844)
Efficiency > 60% (n=1731)
Base percentage (%)
gRNA
PAM
C
Efficiency < 5% (n=1844)
Efficiency > 60% (n=1731)
Base percentage (%)
gRNA
PAM
G
Efficiency < 5% (n=1844)
Efficiency > 60%(n=1731)
Base percentage (%)
gRNA
PAM

S40

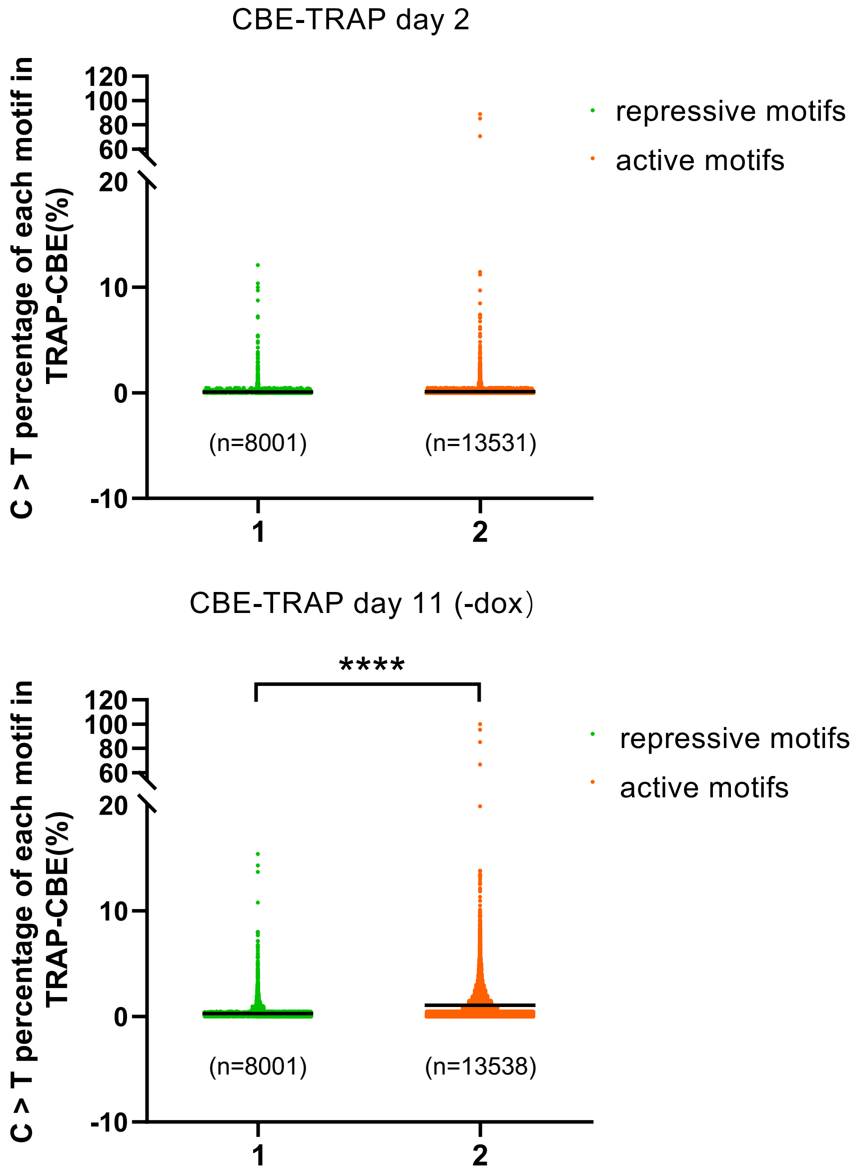

CBE-TRAP day 2
C > T percentage of each motif in TRAP-CBE(%)
120
100
80
60
20
10
0
-10
(n=8001)
(n=13531)
1
2
repressive motifs
active motifs
CBE-TRAP day 11 (-dox)
****
C > T percentage of each motif in TRAP-CBE(%)
120
100
80
60
20
10
0
-10
(n=8001)
(n=13538)
1
2
repressive motifs
active motifs

**S41**

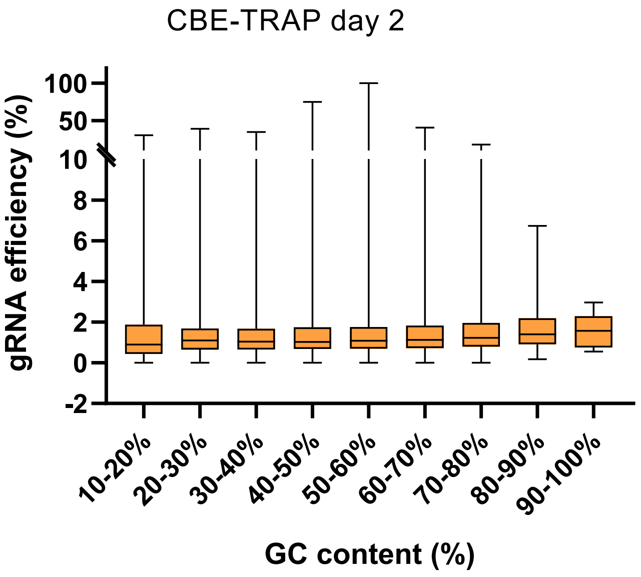

CBE-TRAP day 2
gRNA efficiency (%)
100
50
10
8
6
4
2
0
-2
10-20%
20-30%
30-40%
40-50%
50-60%
60-70%
70-80%
80-90%
90-100%
GC content (%)

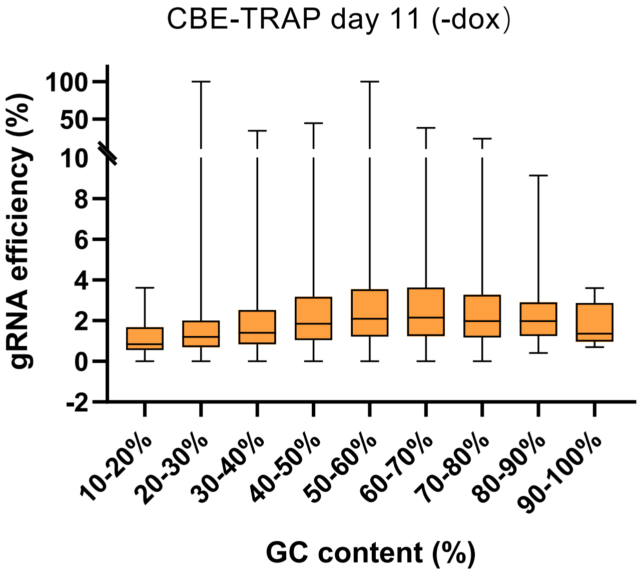

CBE-TRAP day 11 (-dox)
gRNA efficiency (%)
100
50
10
8
6
4
2
0
-2
10-20%
20-30%
30-40%
40-50%
50-60%
60-70%
70-80%
80-90%
90-100%
GC content (%)

S42

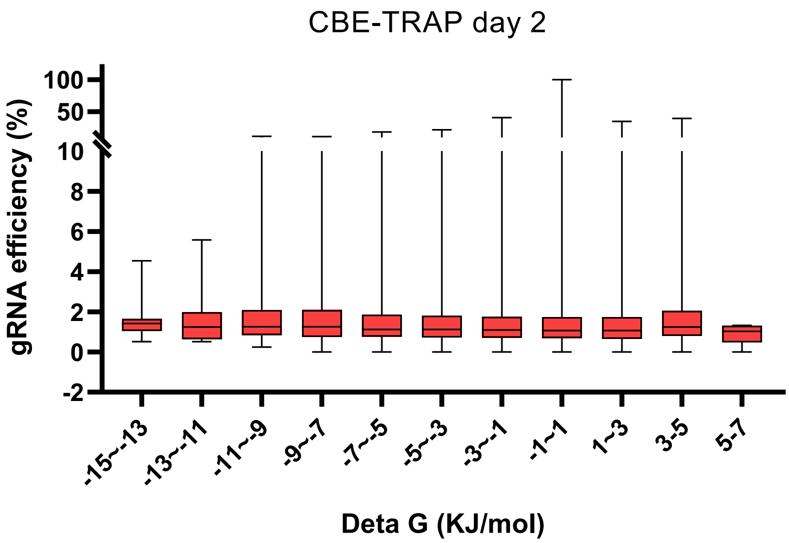
CBE-TRAP day 2
gRNA efficiency (%)
100
50
10
8
6
4
2
0
-2
-15~-13
-13~-11
-11~-9
-9~-7
-7~-5
-5~-3
-3~-1
-1~1
1~3
3-5
5-7
Deta G (KJ/mol)

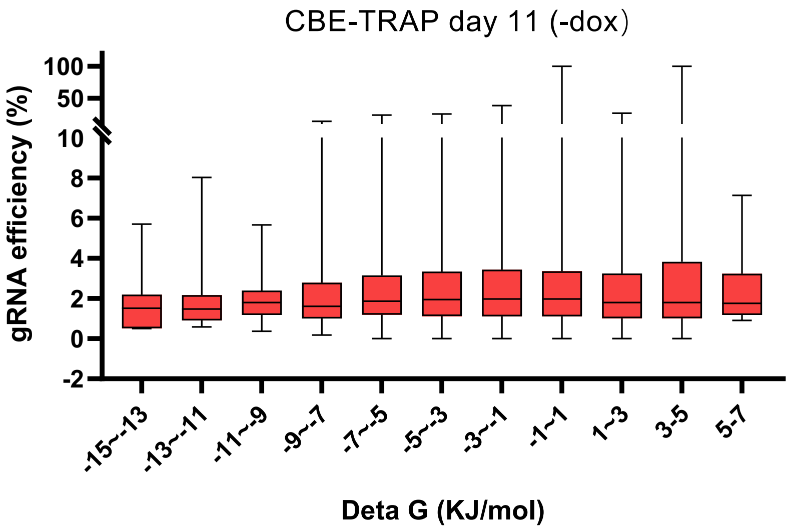
CBE-TRAP day 11 (-dox)
gRNA efficiency (%)
100
50
10
8
6
4
2
0
-2
-15~-13
-13~-11
-11~-9
-9~-7
-7~-5
-5~-3
-3~-1
-1~1
1~3
3-5
5-7
Deta G (KJ/mol)

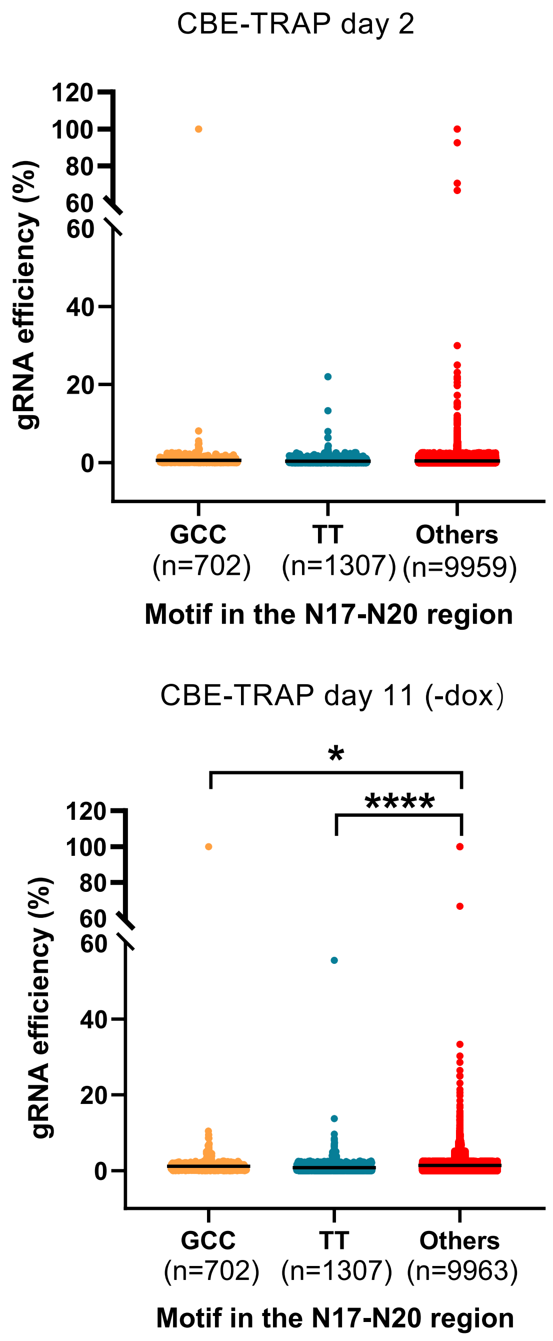
CBE-TRAP day 2
120
100
80
60
60
40
20
0
gRNA efficiency (%)
GCC
(n=702)
TT
(n=1307)
Others
(n=9959)
Motif in the N17-N20 region
CBE-TRAP day 11 (-dox)
*
****
120
100
80
60
60
40
20
0
gRNA efficiency (%)
GCC
(n=702)
TT
(n=1307)
Others
(n=9963)
Motif in the N17-N20 region

S44

**a**

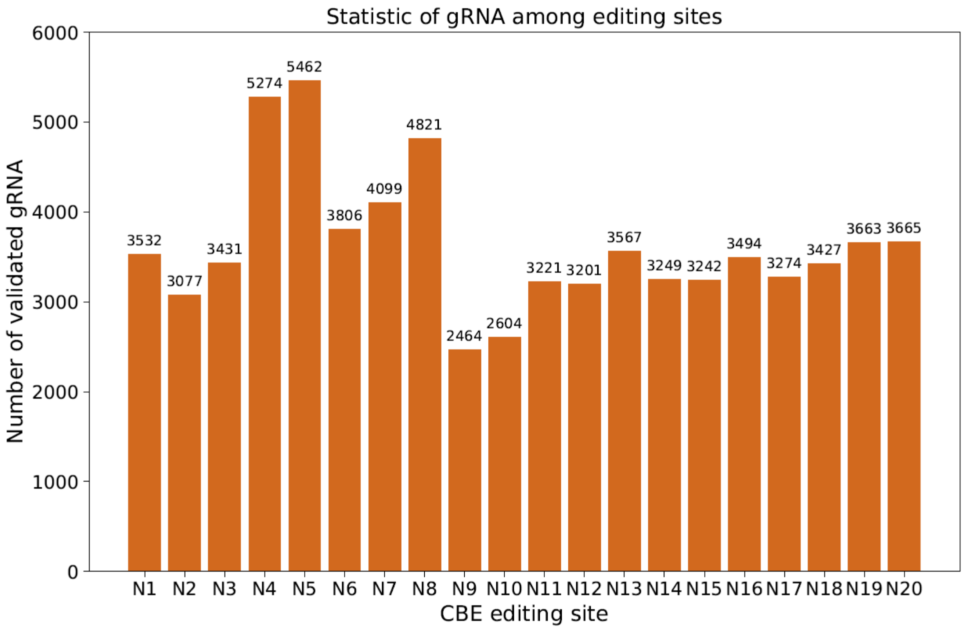

Statistic of gRNA among editing sites
Number of validated gRNA
CBE editing site
0
1000
2000
3000
4000
5000
6000
N1 N2 N3 N4 N5 N6 N7 N8 N9 N10 N11 N12 N13 N14 N15 N16 N17 N18 N19 N20
3532 3077 3431 5274 5462 3806 4099 4821 2464 2604 3221 3201 3567 3249 3242 3494 3274 3427 3663 3665

**b**

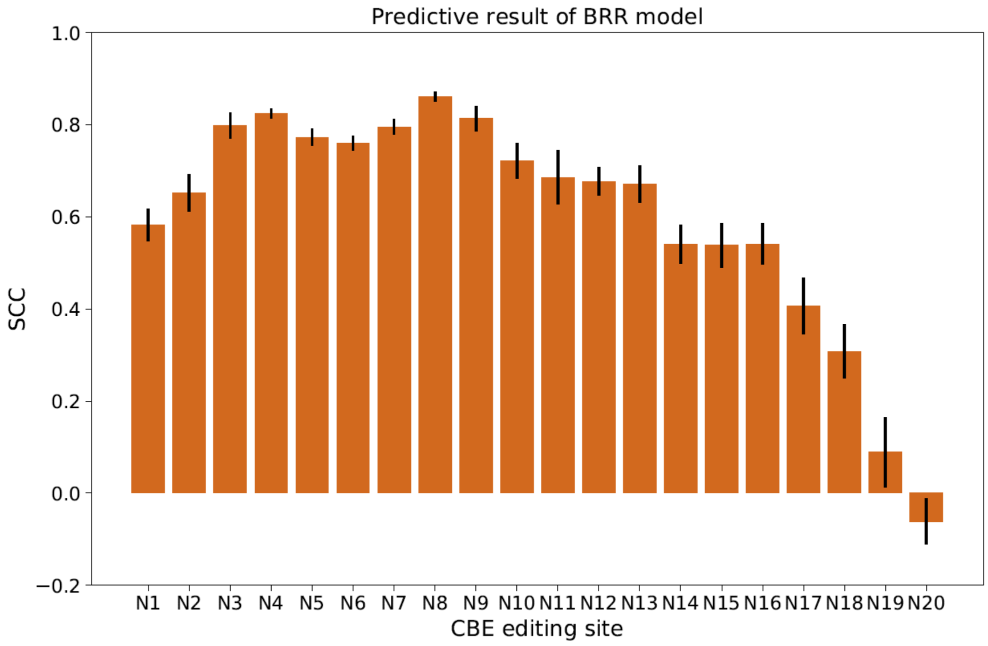

Predictive result of BRR model
SCC
CBE editing site
−0.2
0.0
0.2
0.4
0.6
0.8
1.0
N1 N2 N3 N4 N5 N6 N7 N8 N9 N10 N11 N12 N13 N14 N15 N16 N17 N18 N19 N20

# S45

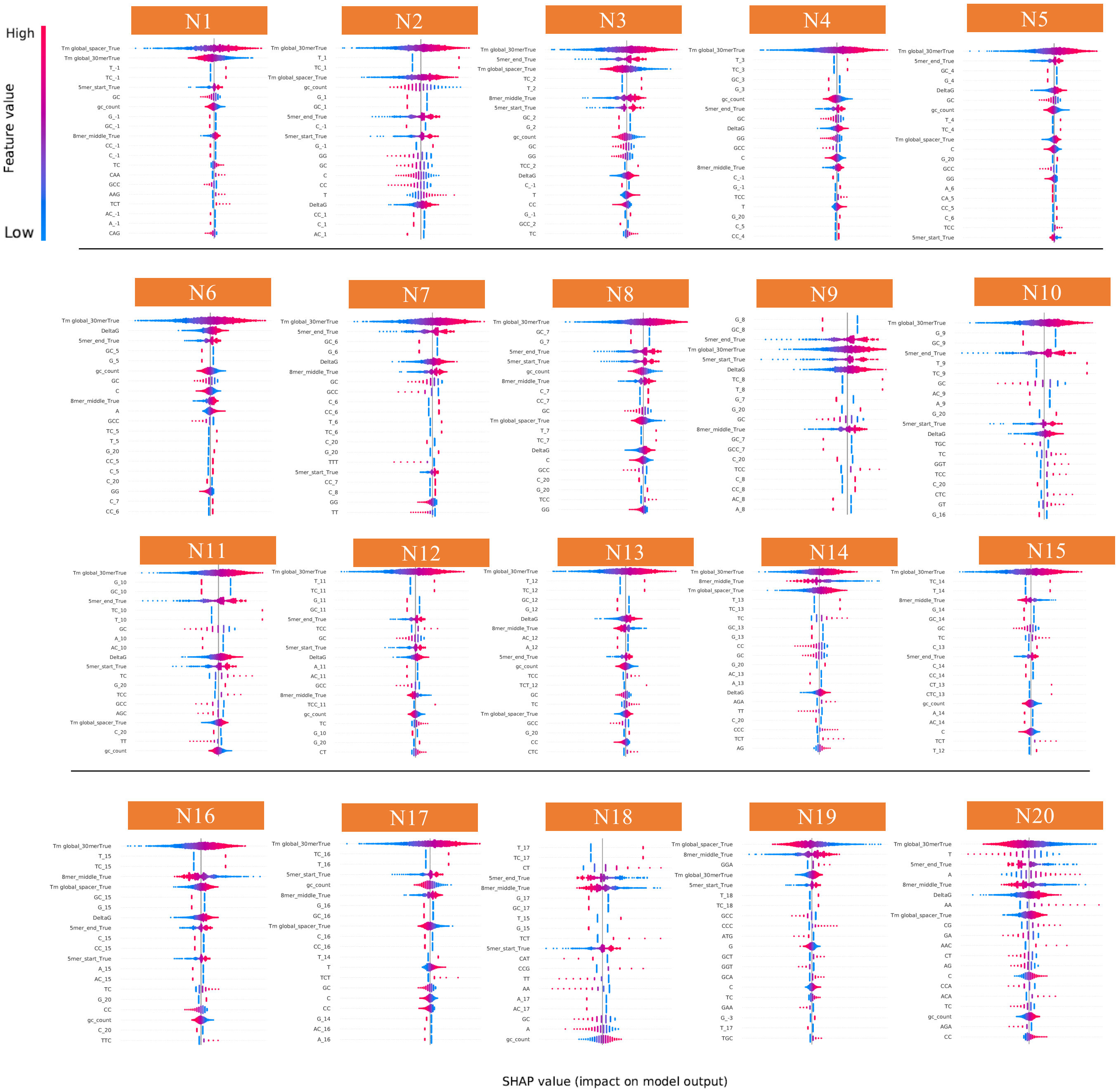

High
Feature value
Low
N1
N2
N3
N4
N5
N6
N7
N8
N9
N10
N11
N12
N13
N14
N15
N16
N17
N18
N19
N20
SHAP value (impact on model output)
